# *Azospirillum*: effective plasmid management

**DOI:** 10.64898/2026.09.14.751567

**Authors:** Natalia O. Dranenko, Mikhail S. Gelfand

## Abstract

*Azospirillum* is a genus of nitrogen-fixing alpha-proteobacteria whose genomes consist of a chromosome and several large secondary replicons. The organization and evolution of these replicons remain poorly understood. We analyzed 22 complete *Azospirillum* genomes, containing six to ten replicons, and characterized their gene composition, replication, segregation, and dimer resolution systems.

The largest replicon contains most of the essential genes and is the chromosome. Other replicons can be classified based on genecomposition, similarity of the plasmid replication protein RepA, and similarity of the ParAB segregation proteins. Overall, groups obtained using different approaches agree with each other. Analysis of gene composition also identified fusions or splits of secondary replicons.

None of the large secondary replicons of *Azospirillum* strains contain the *repABC* replication system, and all of them are iteron plasmids. For all groups of replicons, the RepA box motifs and the structure of the plasmid replication origin *oriV* were predicted.

Most replicons use the ParABS system for segregation. In rare cases, the replicon contains *repA* from one group and *parAB* from another. The position of the *parAB* operon relative to the replication origin is determined by the *parAB* group, and not by *repA*.

Most secondary replicons apparently use an FtsK-dependent pathway for dimer resolution, relying instead on the chromosomally encoded XerCD proteins. In most cases, the identified *dif* motifs do not distinguish groups of replicons.

**Significance statement:** Approximately 10 % of bacteria have genomes with large secondary replicons. Some of them contain more than one such replicon, five to nine in *Azospirillum* spp. Such secondary replicons are usually present in a genome as a single copy; however, the mechanisms underlying their replication, segregation, stable inheritance, and evolution remain largely uncharacterized. We analyzed 22 complete *Azospirillum* genomes and described how they replicate, segregate, and resolve dimers. Chromosomes and secondary replicons rely on distinct but coordinated replication and segregation systems. These findings reveal general principles of how multipartite bacterial genomes are organized and evolve.

## Introduction

*Azospirillum* is a genus of plant growth-promoting bacteria which form symbiotic relationships with plant roots and may act as a biofertilizer (Pelagio-Flores et al. 2025). Due to its importance for agriculture, *Azospirillum* has been studied in detail in the context of its effect on plant growth and yield increase, but the genome of these bacteria itself is of particular interest. *Azospirillum* belongs to about 10% of bacteria whose genome structure, unlike that of the majority (diCenzo and Finan 2017), contains large secondary replicons, chromids or megaplasmids (Gao et al. 2019 Aug 8; Dranenko et al. 2023). Such genomic structure is well-known for bacteria associated with plants. *Azospirillum* is even more unusual, as bacteria of this genus have remarkably many large, stable secondary replicons.

### Chromosome and plasmid replication systems

Hence, there arises a problem of coordinated replication of these replicons and their segregation in daughter cells after division. The replication of the primary replicon and the secondary ones should happen consistently and end at approximately the same time (Fournes et al. 2018). The chromosome replication starts at the conserved *oriC* region and is orchestrated by the DnaA protein, binding a 7 bp sequence, the DnaA-box (Sernova and Gelfand 2008; Hansen and Atlung 2018). Despite the fact that essential genes may be present on secondary replicons, only the primary replicon replication starts at the *oriC* origin, while secondary replicons follow plasmid replication pathways (Fournes et al. 2018).

The plasmid-type replication starts at plasmid origin of replication *oriV* and may occur independently during the cell cycle and is typical for multicopy plasmids (Nordström and Dasgupta 2006). Replication of low-copy plasmids, such as megaplasmids, chromids, and secondary chromosomes (the terms are traditional for taxa and not very consistent (Dranenko et al. 2023)), also is of the plasmid type, but under much stricter control (diCenzo and Finan 2017). As each of the daughter cells needs a copy of a plasmid that exists in a small number of copies in the maternal cell, the process is regulated at several stages: 1) the plasmids replicate frequently enough so that there are copies for both daughter cells; 2) the plasmids must be correctly partitioned to be physically distributed to the daughter cells; 3) the plasmid dimers must be resolved into monomers for proper distribution (Pinto et al. 2012).

Some *Alphaproteobacteria* have secondary replicons with the copy number comparable to the number of copies of chromosomes (Suzuki et al. 2001; Döhlemann et al. 2017). Most of them use RepABC-type systems for replication (Pinto et al. 2012). Three genes located in the same operon, *repA*, *repB*, and *repC*, control plasmid replication and segregation. RepC is essential and sufficient for replication, RepA and RepB manage the partitioning and are members of the ParA and ParB partitioning protein families respectively (Żebracki et al. 2015). The replication origin is inside the *repC* gene (Pinto et al. 2012). It is different by structure from classical plasmid origins as it does not contain groups of direct repeats, or iterons. RepC of the Ti plasmid of *Agrobacterium tumefaciens* binds an AT-rich DNA sequence that includes imperfect dyad symmetry (Pinto et al. 2011). Moreover, the RepC protein initiates replication only of the plasmid in which it is encoded, but not of other RepC-dependent plasmids (Cervantes-Rivera et al. 2011; Pinto et al. 2011).

Many plasmids use a different replication system, containing replication activator RepA that binds iterons. In such plasmids, iterons, short direct repeats, play a crucial role in the initiation of replication and control of the plasmid copy number (Konieczny et al. 2014). The iteron length varies from 17 to 22 nucleotides (Konieczny et al. 2014). The number of iterons varies from three to seven (Chattoraj 2000; Loftie-Eaton and Rawlings 2010), sometimes reaching ten (Konieczny et al. 2014; Maurya et al. 2023), moreover, in some cases they form several positionally independent groups (Page et al. 2001). The number and position of iterons control the plasmid copy number (Maurya et al. 2023), more iterons yielding fewer copies (Chattoraj 2000; Loftie-Eaton and Rawlings 2010). Initiation of replication in such plasmids involves the plasmid initiator protein from the Rep family and the chromosomal replication initiator DnaA, the latter binding to one or more DnaA-boxes at the plasmid origin (Konieczny et al. 2014).

### Partition of replicons

Correct partition of replicons into daughter cells is particularly important for genomes with several significant secondary replicons. The commonest chromosomal partition system in bacteria is ParABS (Jalal and Le 2020). It consists of the ParA and ParB proteins, as well as the *parS* site encoded on the same chromosome (Wang et al. 2013). ParA is an ATPase motor protein, ParB is a centromere-binding protein and *parS* is a centromere-like DNA site, which is generally palindromic (Livny et al. 2007; Chu et al. 2024). During the partitioning process, ParB first binds to *parS* and recruits additional ParB proteins to non-specific DNA in the surrounding area, forming a nucleoprotein complex (Jalal and Le 2020). Then, interaction of ParB-*parS* complex with ParA leads to the DNA movement (Jalal and Le 2020).

In most studied cases the *parA* and *parB* genes form an operon located relatively close to the origin of replication (Surtees and Funnell 2003). The *parS* site is also located near the origin and segregates first during the chromosome replication; so this localization is believed to be essential (Jalal and Le 2020). For example, in *Caulobacter crescentus*, the *parS* sites are located at 8 kbp distance from the replication origin and segregation stalls until the *parS* sites are replicated (Menikpurage et al. 2023).

In RepABC-type plasmids of *Alphaproteobacteria*, the operon structure is different from the chromosomal one (Pinto et al. 2012). In the chromosome, the *parAB* operon and the *dnaA* gene are independent, while in alphaproteobacterial *repABC* plasmids, replication and partition genes form one operon (Pinto et al. 2012). The *repA*, *repB*, and *repC* genes are transcribed from promoter positioned upstream of *repA*. Sometimes one more presumably protein-coding gene, *repD*, is positioned between *repA* and *repB*. Due to the diversity of *repD* sequences in related plasmids, it is assumed that the function of *repD* is to control expression of downstream genes rather than to encode a protein (Chai and Winans 2005). Between *repB* and *repC*, there is also a gene encoding a short antisense RNA, which downregulates *repC* (Pinto et al. 2012).

### Resolution of replicon dimers

During replication of circular DNA in bacteria, dimers can be formed following events of homologous recombination (Castillo et al. 2017). Although there are cellular mechanisms that preclude crossover necessary for homologous recombination, resolution of dimers is still necessary (Castillo et al. 2017). Dimers or multimers of higher orders result from homologous recombination between sister replicons, which reduces the number of independent replicon copies (Fournes et al. 2025). Chromosomes or large secondary replicons, which are usually present in the genome in one copy, contain genes whose loss strongly affects fitness, and hence such events will be fatal for the cell (Val et al. 2008).

Resolution of multimers in bacteria is performed by two or, less frequently, one site-specific recombinase (Castillo et al. 2017). The most common systems which work for chromosomes consist of two site-specific tyrosine recombinases, XerC and XerD, encoded on chromosomes and working on *dif* sites (Castillo et al. 2017). A *dif* site is a pseudopalindromic sequence of 28-30 bp. It consists of two 11 bp binding sites for XerC and XerD separated by a 6-8 bp linker (Castillo et al. 2017). It is located near the terminus of replication, usually close to the GC-skew turning point (Kono et al. 2011; Bhowmik et al. 2018). The area of about 10 kbp around the *dif* site contains multiple oppositely oriented KOPS (FtsKOrienting Polar Sequences) directing DNA translocase FtsK; the latter segregates sister chromosomes and activates XerCD recombination in case of dimerization. This area also contains *matS* sites recognized by the MatP protein, which delays segregation of the replication terminus *ter* and hence prevents premature separation of sister chromosomes (Fournes et al. 2025).

Secondary replicons can either contain a *dif* site and also use chromosomally encoded XerC and XerD proteins, or carry their own site-specific recombinases binding *dif*-like sites (Val et al. 2008; Kono et al. 2011; Fournes et al. 2025). Dimer resolution for secondary replicons can follow the chromosomal scenario and be FtsK-dependent or, as in the case of *Escherichia coli* secondary replicons, FtsK-independent (Fournes et al. 2025). FtsK-independent dimer resolution requires different accessory proteins and, sometimes, around 200 bp accessory sequences, which is flanking the XerC binding site (Val et al. 2008; Fournes et al. 2025). For *Enterobacteriaceae*, it has also been shown that the mechanism of dimer resolution differs depending on the size and copy number of a secondary replicon (Fournes et al. 2025). For large replicons, a FtsK-dependent dimer resolution pathway involving elements controlling FtsK, directional bias of KOPS, and *matS* sites is typical (Fournes et al. 2025). Plasmids of 30-100 kbp long tend to have no *dif* or *dif*-like site, while in plasmids of 25 kbp and smaller such sites are usually present (Fournes et al. 2025).

Here, we present a comprehensive study of genes and functional sites involved in replication, segregation, and dimer resolution during *Azospirillum* cell division. Using a comparative bioinformatic analysis of 22 complete *Azospirillum* genomes (6–10 replicons each), we show that replicon similarity groups based on gene content are congruent with those based on RepA replication initiator proteins and *oriV* similarity, indicating that the “identity” of a replicon is defined by its replication system. In contrast, *parAB* genes encoding segregation proteins are less stable in terms of replicon similarity groups. We also predict *dif* sites for chromosome dimer resolution and demonstrate that they are present on all but the 1–2 smallest replicons per genome, suggesting that these replicons employ the canonical FtsK-dependent dimer resolution system, whereas the smallest replicons lacking *dif* sites may rely on an alternative mechanism.

## Results

### *Azospirillum* phylogeny and genome composition

We analyzed 22 complete *Azospirillum* genomes, each composed of six to ten replicons (**Supplementary Table 1**). A phylogenetic tree of these strains (**Figure 1**) featured two major clades and two single-strain branches. The largest replicon (2.5–3.3 Mb) in each strain is presumed to be the chromosome. Most secondary replicons are large, with only eleven shorter than 100 kbp and only two of these shorter than 50 kbp. The replicon sizes are more stable in the clade with shorter internal branches compared to the clade with longer internal branches.

**Fig. 1.**
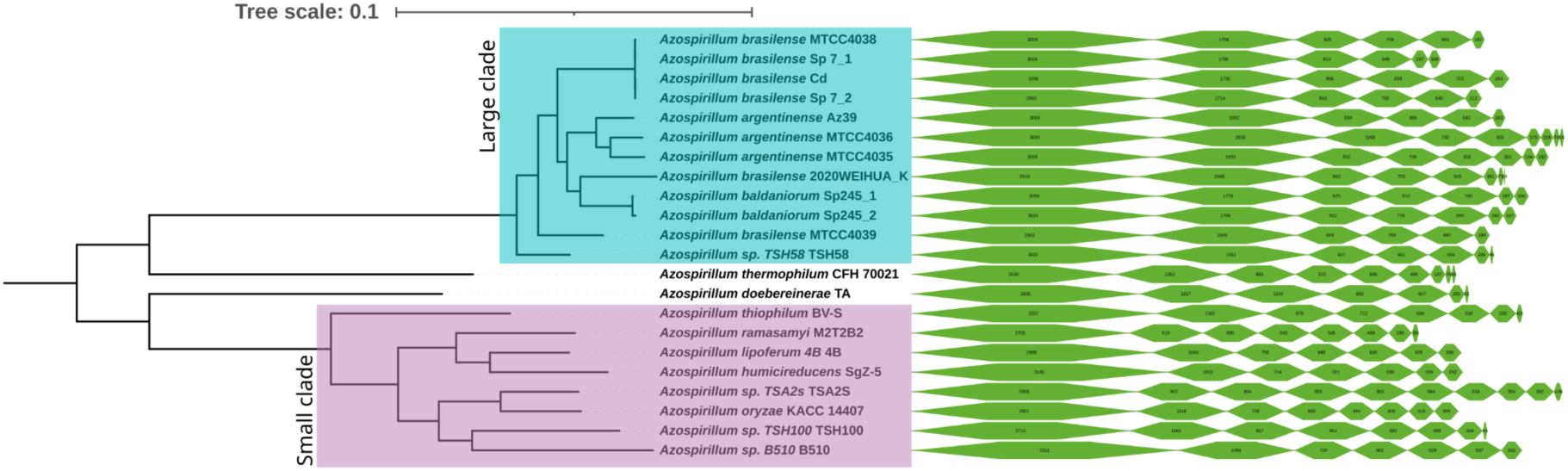
The phylogenetic tree of *Azospirillum* strains rooted on long branches with the composition of replicons sorted by size.

The average GC-content of the chromosomes is 68.3%, the average GC-content of all other replicons is lower, 67.9%, and for the smallest replicons of each strain, the average GC-content is 66.4% (**Supplementary Figure 1**). However, there is no direct relationship between the replicon size and its GC-content.

### Location of essential genes in replicons

For all available strains, we checked to what replicons belong genes encoding essential bacterial domains (see Methods). The gene encoding the UBA/TS-N protein domain (PF00627) was not present in *Azospirillum* genomes. Three out of 22 genomes missed also other essential genes: in one genome the gene encoding signal peptidase II was missing; in two strains, the S20 small ribosome subunit gene has frameshift at the sequence in file start point, however, growth without it was observed for some bacteria (Nikolaeva et al. 2021); finally, one strain had frameshifts in genes encoding two phenylalanyl– and arginyl-tRNA synthetases, likely caused by sequencing errors.

138 essential bacterial domains were present in 110 different gene groups. 89 of these groups contained exactly one gene per genome, in four groups mentioned above, some strains did not have the respective gene, and in 17 groups, more than one gene with the same function was present. Moreover, in seven cases, an essential domain was observed in different orthologous groups.

Single-copy genes encoding proteins with essential functions were usually, in 63 cases out of 89, located in the largest replicon, less often in the second one in size, and are almost never found in smaller replicons. In all but one case of essential genes present in more than one copy, there were paralogs in secondary replicons.

In two strains, *Azospirillum brasilense* Sp 7_2 and *Azospirillum brasilense* Cd, rRNA genes are located in main and secondary replicons and do not form operons. Two strains have only one rRNA operon, in *Azospirillum humicireducens* SgZ-5 it lies in the chromosome and in *Azospirillum baldaniorum* Sp245_1, in a secondary replicon. The remaining 18 strains contain from five to nine rRNAs in operons located in three to six replicons. Moreover, rRNA sequences on the same replicon may differ significantly (**Supplementary Figure 2**). In some cases, rRNA sequences from the same strain or even replicon differ more than between different strains.

Thus, if an essential gene is unique, it is usually located in the chromosome, and the mechanism of chromosome segregation during division guarantees transfer of these genes to the daughter cells. Additional genes encoding the same essential bacterial domains differ from chromosomal ones and are often located in secondary replicons.

### Phylogenetic relationships between replicons

Replicons can be grouped based on various characteristics, in particular, by size, by gene composition, by the sequence of replication genes, and by the sequence of segregation genes. **Table 1** presents these classifications for all replicons of the strains under consideration. In most cases, the classifications are similar, except for the grouping by size. Each of the methods for classifying replicons is described in detail in the following sections.

**Table 1.**
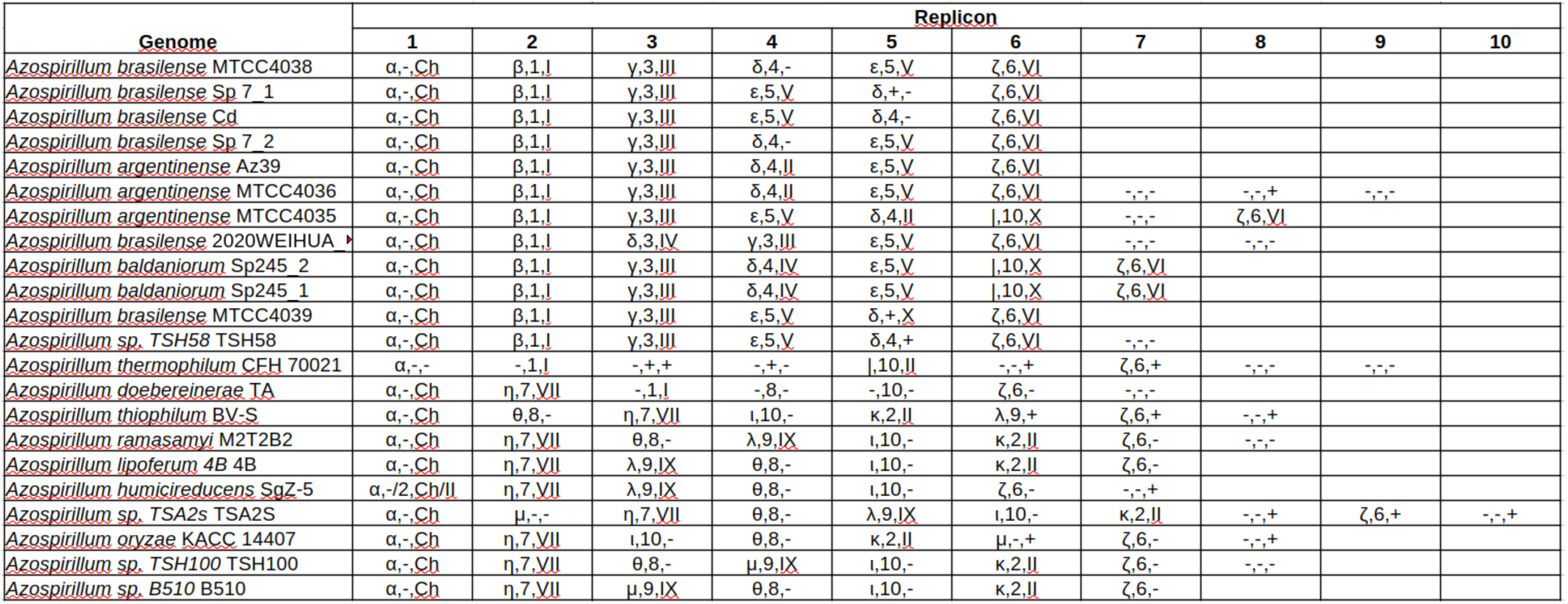
Different approaches to classifying *Azospirillum* replicons: based on gene composition (Greek letters), replication proteins RepA (Arabic numerals), and segregation proteins ParAB (Roman numerals).

To reconstruct the evolutionary relationships in terms of gene composition between replicons two independent approaches were used (see Methods). The groups of related replicons determined using each of the methods are shown in **Figure 2**. Both methods lead to generally the same results. As a rule, one group includes replicons of close size.

**Fig. 2.**
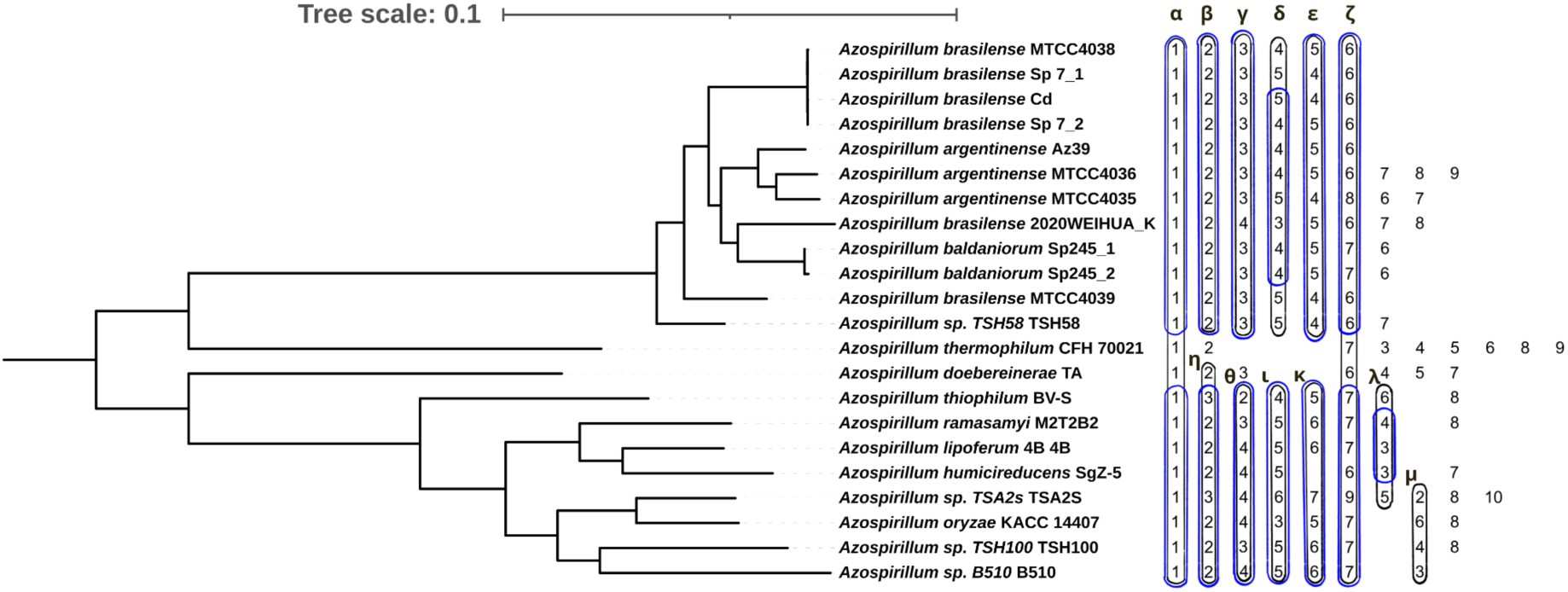
The phylogenetic tree of *Azospirillum* strains rooted on long branches and the composition of replicons sorted by groups of similarity. The replicons of each strain are numbered by size from the largest replicon (1), chromosome, to the smallest one. The replicons are sorted by similarity. The frame color reflects the way the replicons are matched: blue – the Jaccard coefficient for orthologous groups with no paralogs, black – the modified Jaccard coefficient (see Methods).

The replicon similarity follows the logic of the tree structure: the main replicon, which contains the largest number of core genes, retains its composition in all *Azospirillum* strains; the remaining replicons are divided into similar groups in two large clades; two strains on long branches contain mostly unique replicons.

When analyzing the similarity of replicons, we considered both the best bidirectional hits and the best unidirectional ones (**Supplementary Figure 3**). In many cases the best bidirectional hit connects two replicons of comparable size, while smaller replicons are linked to the same replicons as unidirectional best hits.This suggest replicon merge/split events.

### Replication of the main replicon

The largest replicon, the chromosome, contains the traditional chromosomal replication origin, *oriC*. The chromosome replication origin sequence is known for *Azospirillum baldaniorum* Sp245_2. It is located between the *yqfL* and *hemE* genes. Based on the origin sequence, the DnaA box motif was predicted. The *oriC* contains six DnaA boxes, grouped as two remote co-directed sites and four repeats, the first, second and fourth of which lie in one direction, and the third in the other (**Figure 5**). The *parAB* operon is located at a distance of about 12.4 kbp from the origin of replication. Two palindromic *parS* sites are located downstream of the *parB* gene at distances of about 600 and 2500 bp from *parB* stop and at about 11.8 and 9.9 bp from *oriC*, respectively. Chromosomal *parS* sites are palindromic and have a length of 16 nucleotides (Livny et al. 2007).

### Replication of secondary replicons

Unlike typical large plasmids of alpha-proteobacteria (Pinto et al. 2012), none of the large secondary replicons of *Azospirillum* strains contain the *repABC* replication system. All of them are iteron plasmids.

We searched for genes encoding plasmid replication proteins in all *Azospirillum* strains and found 127 candidate RepA proteins. All but ten of these proteins form ten clades in the phylogenetic tree (**Figure 3**).

**Fig. 3.**
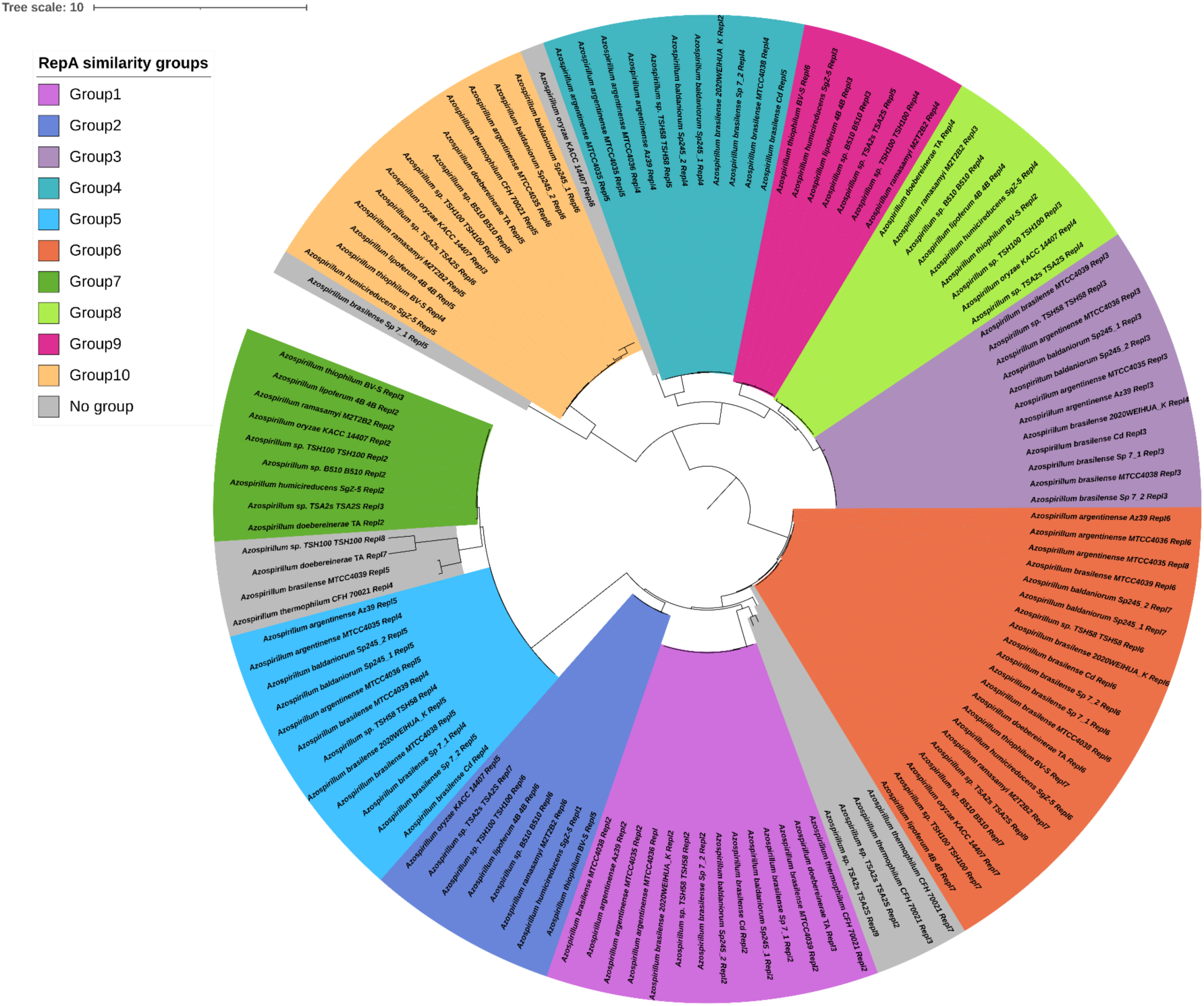
RepA proteins tree. The tree is rooted at the point between two main *Azospirillum* tree clades. The colored clades are defined by protein similarity with 82% identity threshold. The proteins colored in gray are not closely similar to RepA from other replicons.

These ten clades correspond to the replicon similarity groups determined by gene composition taking into account the presence of two large branches on the tree (**Figure 4**). In addition, the similarity in RepA makes it possible to extend some components of the similarity made by gene composition.

**Fig. 4.**
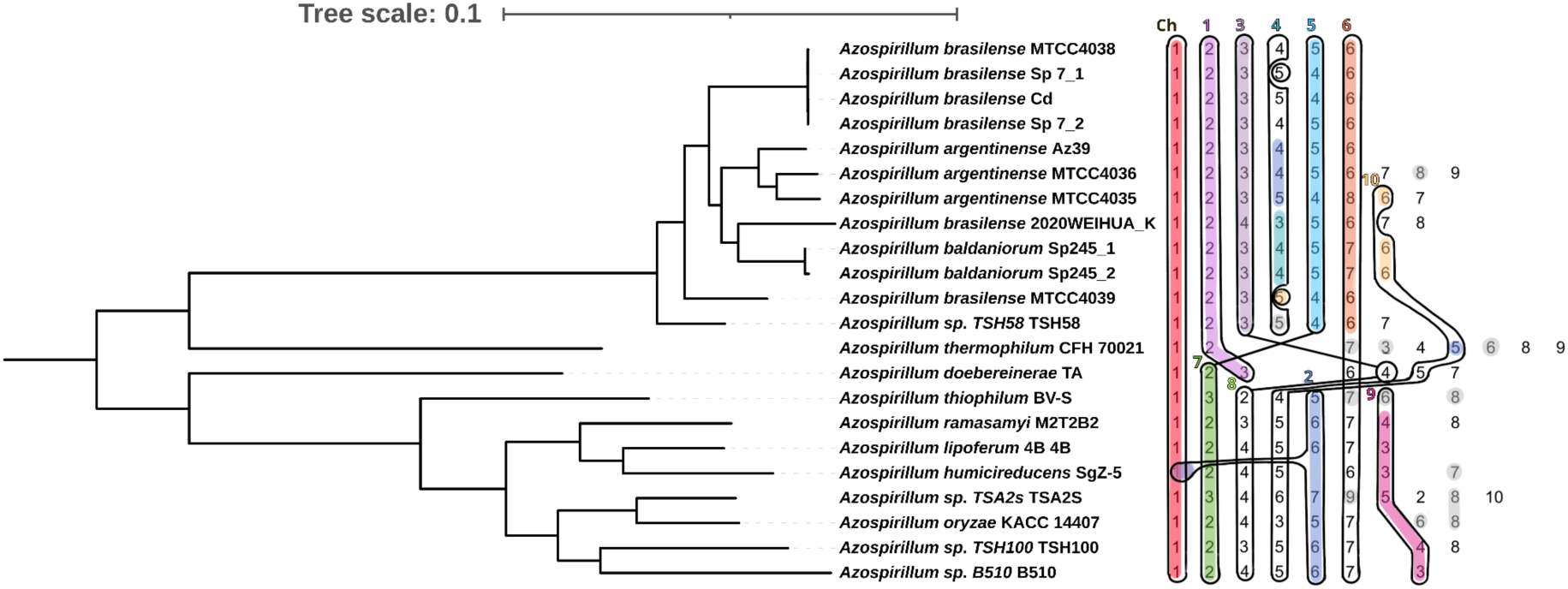
The phylogenetic tree of *Azospirillum* strains rooted on long branches and RepA proteins similarity groups. Replicons with genes encoding RepA proteins of one similarity group are combined into one black frame and the boldface colored numbers at the top correspond to RepA similarity groups from Figure 3. The frames for proteins of close clades are connected. The replicon numbers within the RepA similarity frames are colored according to the *parAB* genes detected on these replicons (see the text). Grey replicon marks indicate *parAB* genes that are not included in similarity groups. In *Azospirillum argentinense* MTCC4036, *parA* in replicon 5 is formed by two adjacent, partial ORFs; the figure features the full form.

The similarity of proteins within a group is more than 82%, and if a group is represented in only one branch of the tree, it is more than 90%, while the similarity of proteins between groups is usually about 35%, the exception being groups 5 and 7 similar, on average, at about 80%, and groups 3 and 8 similar at about 65%.

A gene encoding the RepA protein, similar to proteins from relatively small secondary replicons in other strains (Group 2), was also found in the chromosome of *Azospirillum humicireducens* SgZ-5 (**Figure 4**). The one-sided best similarities also showed a similarity between this chromosome and the group 2 of secondary replicons (**Figure 3**). This suggests that the chromosome of *A. humicireducens* SgZ-5 contains a fragment or an entire secondary replicon, which may result from integration of the plasmid into the chromosome or an assembly error.

For all groups of secondary replicons determined by the RepA protein similarity, we were able to find the position of the replication origin and the sequence of iterons in them (**Figure 5, Supplementary Figure 4**). For all groups including chromosomes, the replication origin structure is shown with the DnaA boxes and iterons positions and logos.

**Fig. 5.**
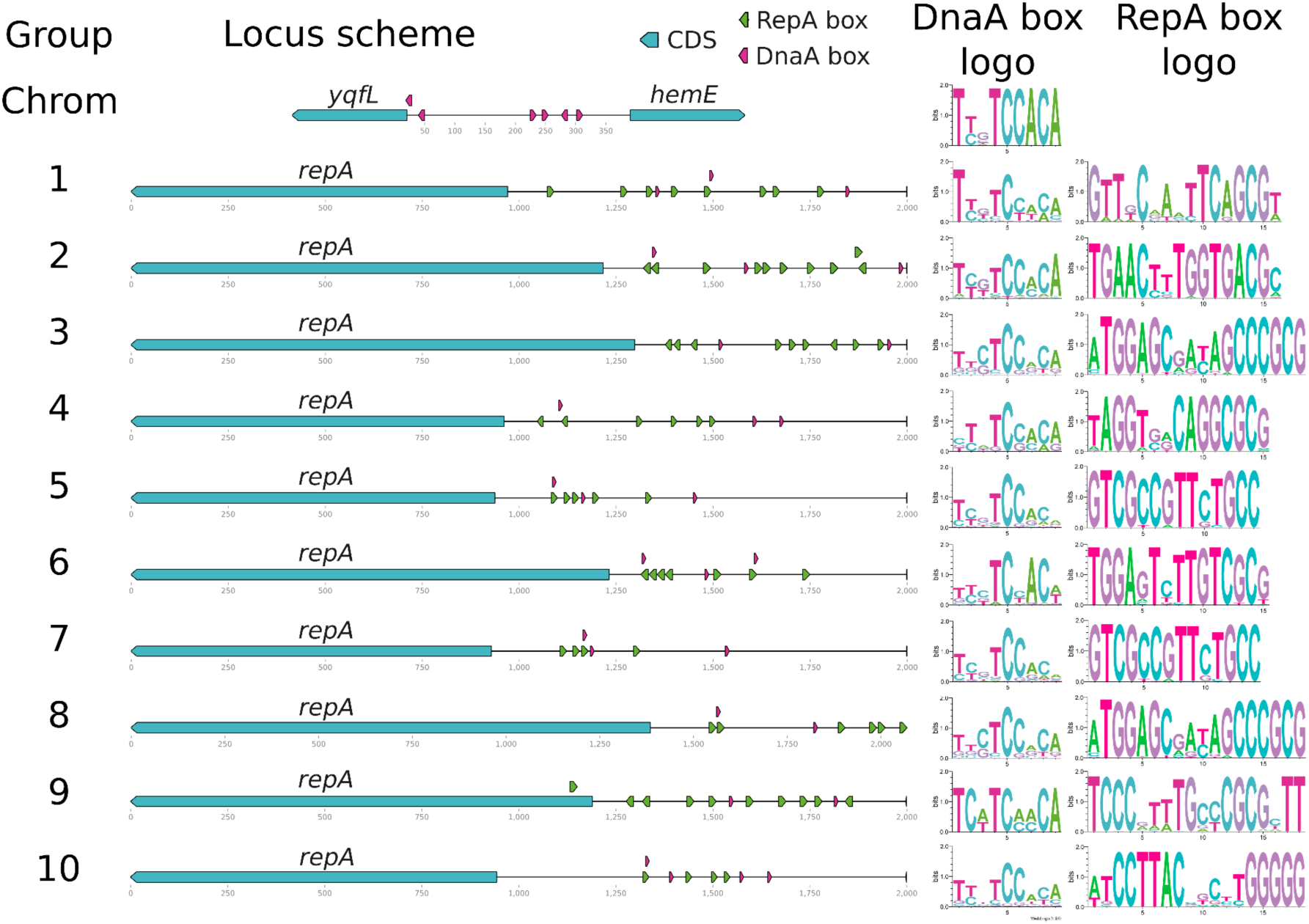
Replication origin structures for chromosome and replicon groups 1-10. Protein coding sequences are shown as blue arrows, DnaA boxes are shown as pink arrows, RepA boxes are green arrows. Logos of the corresponding motifs are given for each group of binding sites.

The structure of the replication origin is generally preserved within the replicon group, although particular candidate RepA or DnaA binding sites at particular positions upstream of *repA* may be missing, as some sites are present in all members, while others are found only in subsets. Nevertheless, in each group of secondary replicons, there are at least three iterons and one DnaA box directed opposite to *repA*. The iteron lengths vary by group, and range from 14 to 18 nucleotides. In groups 5 and 7, the RepA proteins are much more similar compared to other pairs of groups and, consistently, the structures of replication origins and the RepA box motifs are similar too. For groups 3 and 8, also more similar than average, the RepA box motifs, but not the structures of *oriV*, are similar.

Almost all replicon groups had remote RepA boxes directed opposite to the main group of iterons, which may contribute to the copy-number control. It was previously shown that on plasmid RK2, the presence of a singleton in the opposite direction to the main group of iterons affects plasmid replication and reduces the copy number nearly two-fold (Maurya et al. 2023).

In all replicons but one there is only one *repA* gene and *oriV* structure. The fifth replicon of *A. argentinense* MTCC4035 has two identical *repA* with the same iteron *oriV* structure upstream of each of these genes. This is a part of a long, about 5.5 kbp, duplication, which may be an assembly error.

### Segregation of secondary replicons

While linkage of replicons by the overall gene composition is in good agreement with clustering by replication systems, clustering by genes of segregation proteins ParAB is slightly different (**Figure 4, Supplementary Figure 5, Supplementary Figure 6**). Group 6 is divided into subgroups by *parAB*, since these genes are present in only one of them. Group 4 is divided into two subgroups with different *parAB*, one of them being unique and the other similar to Group 2 *parAB* genes, and one replicon with *parAB* genes unique in our data. Group 10 consists of three replicons with common *parAB*, one replicon with *parAB* similar to that of Group 2, and remaining replicons in which *parAB* were not identified. One of the replicons in Group 9 has *parAB* genes unique in our dataset. In Group 8 no candidates in *parAB* genes were identified.

In Group 3, two sets of *parAB* operons are present on all replicons of the group. Both are located quite far from the replication origin at a relatively stable distance to it (∼40 kpb and ∼200 kbp). The presence of two sets of genes for the same segregation system in bacterial replicons is not typical although cases of the presence of several different systems are known (Siguier et al. 2023). In a Group 5 replicon from *A. brasilense* Cd, *parAB* from a set of Group3 *parAB* more remote from *repA* is also found along with surrounding *parS* sites.

For all available *parAB* groups, we determined the structure of the *parABS* locus. Positions of *parS* sites and the logo motif of the proposed *parS* site are shown in **Figure 6**. In case of chromosomal *parABS*, as well as *parABS* from Groups 1, 2, 5, and 7, the distance between the *parAB* operon and the *repA* gene start is stable with two exceptions in Group 5. In Group 4, either the sequence and the distance between *repA* and the *parAB* is similar to that of Group 2, or both are different. Overall, this distance is the largest, about 12600 bp, in chromosomes, while in Groups 1, 2, 4, 5, and 7 of secondary replicons it ranges from 1000 to 7000, and there is no correlation between this distance and the size of replicons in a group (**Supplementary Figure 7**). In Groups 3, 6, 8, 9, 10 of secondary replicons, the distances between the *parAB* operon and the *repA* gene start are diverse even within one group (**Supplementary Table 2**).

**Fig. 6.**
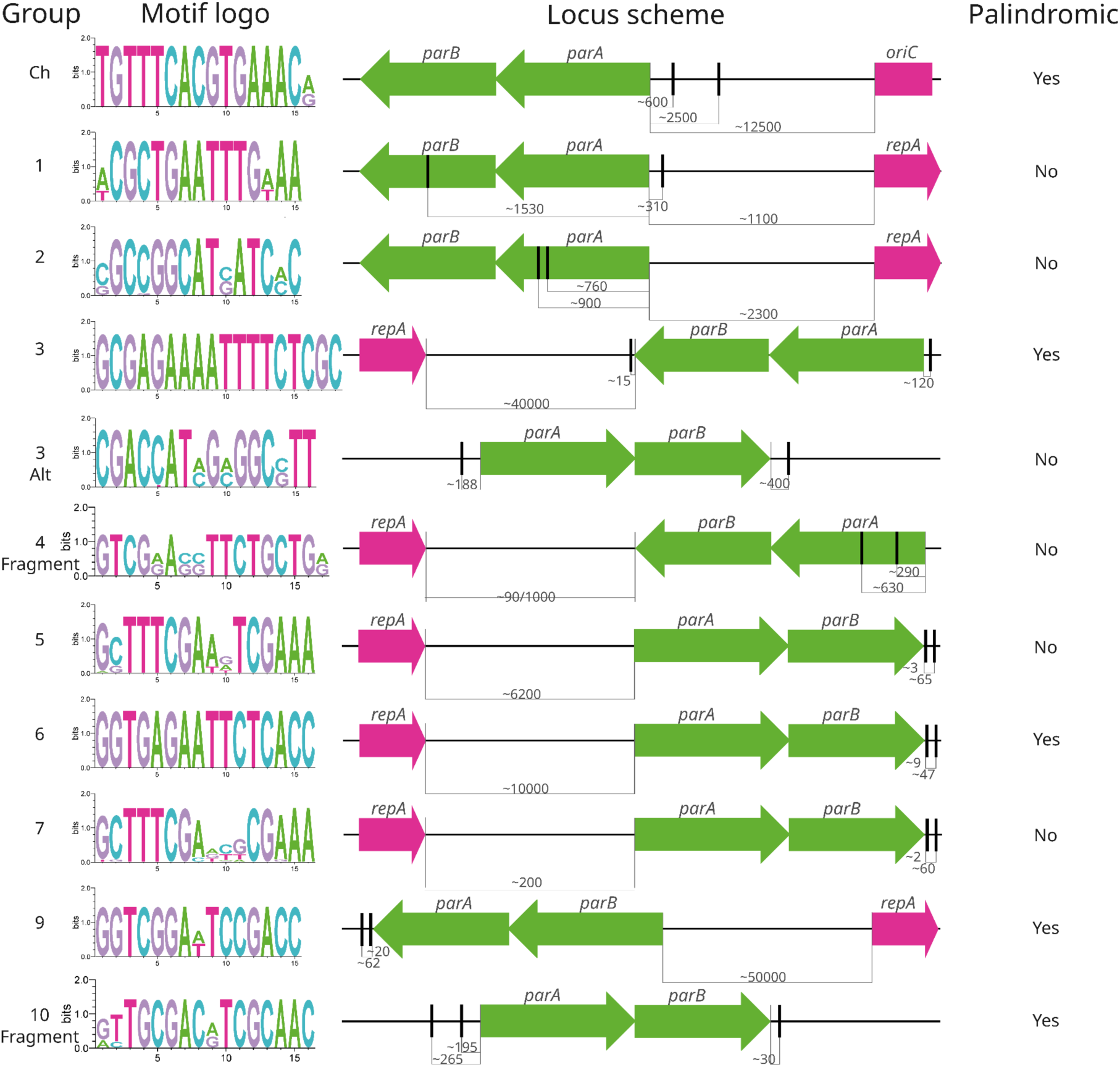
The *parABS* loci for all replicon groups, for which *parAB* genes were predicted. Motifs logos are built for *parS* sites in the *par* locus. The *parAB* genes are represented by green arrows, the *parS* sites are marked by thick vertical lines, the *repA* genes are shown as magenta arrows.

Traditional palindromic *parS* were found in five of the *parABS* groups. In the remaining six groups, non-canonical non-palindromic potential *parS* were identified. The presence of plasmids with the *parABS* segregation system with palindromic and non-palindromic sites in one organism has been observed, for example, in *Rhizobium leguminosarum* bv. *trifolii* (Koper et al. 2016).

### Resolution of dimers

Dimer resolution in bacteria occurs at *dif* or *dif*-like sites by a pair of XerC-XerD recombinases or, less often, a single site-specific recombinase (Castillo et al. 2017). All *Azospirillum* strains in our dataset contain *xerC* and *xerD* genes in the chromosome and some additional tyrosine recombinases in secondary replicons. Only two orthogroups of tyrosine recombinases, XerC and XerD, are present in all strains and their trees match but are slightly different from the strains tree (**Supplementary Figure 8**).

Based on *dif* sites from diverse multichromosomal alphaproteobacteria (Kono et al. 2011), we predicted *dif* sites in chromosomes and secondary replicons of *Azospirillum* (**Figure 7**). All predicted *dif* sites had a 6 bp linker between the XerC and XerD binding sites. The obtained *dif* motifs correlate with the tree structures of the *Azospirillum* strains and XerC-XerD (**Supplementary Figure 8**) and are more similar within two large clades of these trees than between them. There is also an agreement between the *dif* motif and grouping of replicons by similarity of RepA. The *dif* motif of the chromosomes of the large clade and Group 1 significantly differs from other replicons of that clade, whereas in the small clade, all *dif* sites, including the chromosomal ones, are highly similar.

**Fig. 7.**
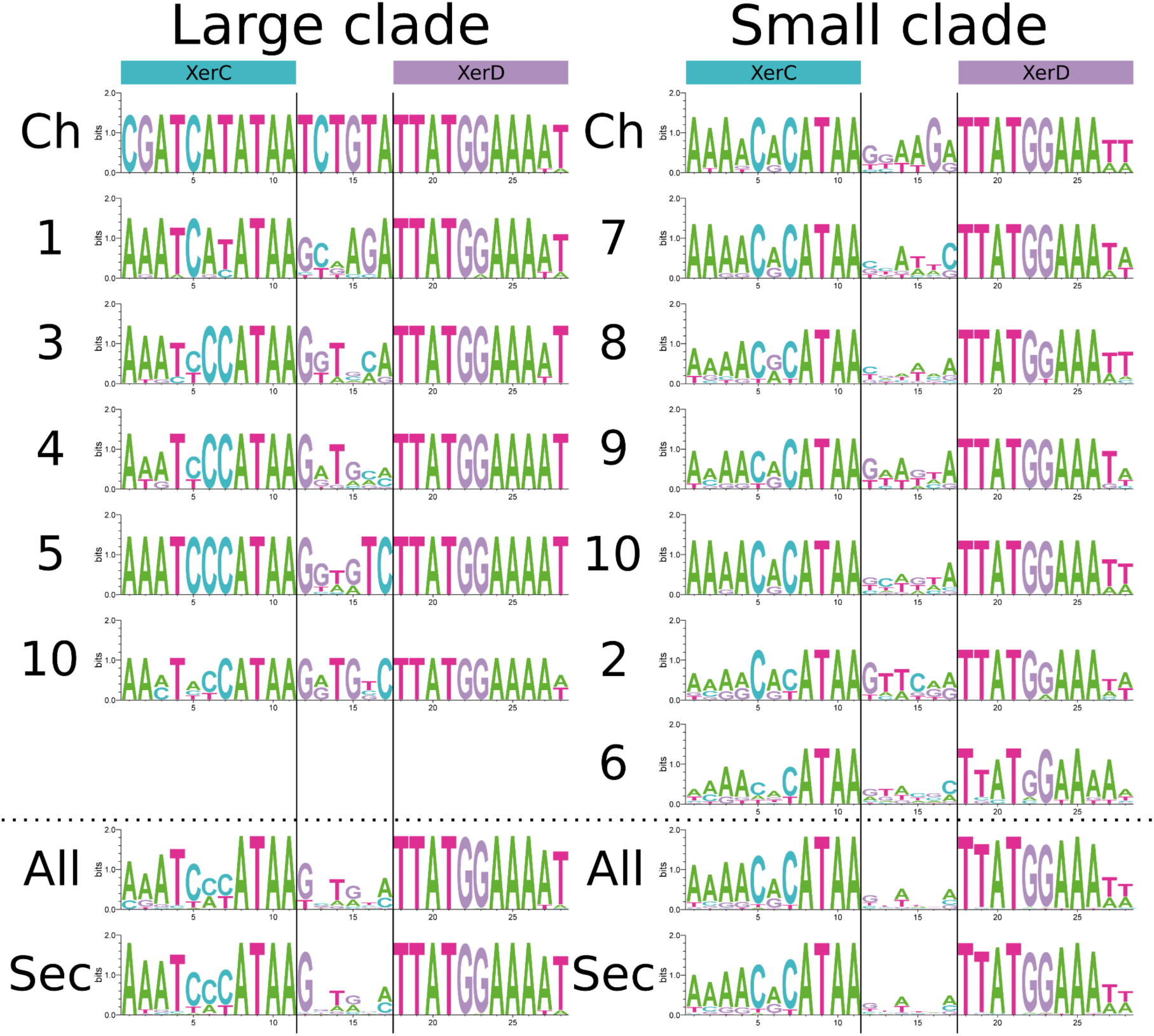
*dif* motifs for replicon groups. Groups containing replicons from two clades are divided according to the XerC and XerD phylogenetic trees. All: all replicons, Sec: only secondary replicons.

All *dif* sites but three are located in intergenic regions, the exceptions seem to be caused by gene annotation errors. Single *dif* sites are present in all replicons except chromosomes of *A. brasilense* Sp 7_2 and *A. brasilense* Cd that have two sites each, a chromosomal-type one and one similar to *dif* from Groups 3-5 and 10 in the large clade. Intergenic regions harboring *dif* sites are similar in the larger replicons of the large clade, but they differ in smaller replicons of the large clade and all replicons of the small clade. The median length of the intergenic region on the side of the XerC box is 116 bp, and on the side of the XerD box – 114 bp. No potential, conserved accessory sequences were found either in the RepA-based groups or in any other group of replicons.

In replicons, the presence of a *dif* site generally corresponds to replicon size. With the exception of *A. thermophilum* CFH 70021, where even some large replicons lacked a *dif* site, the remaining strains lacked this site in at most the two smallest replicons. Among the RepA similarity groups, the absence of *dif* sites was observed exclusively in Group 6, specifically within the large clade, where replicons are significantly smaller than those in the small clade of the same group.

On the chromosomes was also found a candidate *ftsK* gene, so we tested all 4 variants of the KOPS (GGGNAGGG) for presence on *Azospirillum* chromosomes. Based on the location of the sites, their number on the chromosome, and a more pronounced skew signal than the corresponding GC skew in the replicon, the KOPS motif was identified: GGGCAGGG (see **Methods**).

For all groups of RepA similarity and replicons with *dif* that were not included in the similarity groups, as well as replicons without predicted *dif* sites (as a control), we analyzed location of KOPS sites (**Supplementary Figure 9**).

In the chromosome, location of KOPS corresponds to the turnover points in the cumulative GC-skew (**Supplementary Figure 10**). In most cases, *dif* is also located near the turning point of the cumulative GC-skew. As a result, for 17 out of 22 studied strains, the distance between *dif* and the chi-square statistic maximum point does not exceed 5 kbp (**Supplementary Table 3**). In the remaining five cases, the greater distance is due to the specific features of the replicon structure. In three strains, as can be seen from the location of KOPS and the shape of the cumulative GC-skew, an inversion occurred, encompassing *dif*, resulting in two points where the direction of KOPS changes. In two more cases, the KOPS statistics reach a plateau between dif and KOPS direction change point. Thus, the predicted dif sequence is also supported by data on the location of KOPS.

In secondary replicons, *dif* predicted by the sequence and the KOPS direction change point are usually also close, however, there are situations when the KOPS turnover point becomes more noticeable near the origin rather than the replication terminus (**Supplementary Figure 10**). The smallest replicons often lack both *dif* and a sufficient density of KOPS.

## Discussion

*Azospirillum* is a genus of nitrogen-fixing bacteria that are actively used as fertilizers in agriculture (Cassán and Diaz-Zorita 2016; Cassán et al. 2020; Pelagio-Flores et al. 2025). Although they have been studied for more than a century (Pelagio-Flores et al. 2025), the genetic organization of these bacteria leaves many open questions about co-regulation of replication, segregation, and dimer resolution of multiple replicons of comparable size.

*Azospirillum* strains from our dataset had six to ten replicons, the largest of which is the bacterial chromosome. For them, we studied the location of genes encoding proteins with essential functions. We have shown that, as a rule, if, in a genome, there is only one gene with an essential function, this gene is located in a chromosome, while if there is another gene with the same function, the sequences of the encoded proteins differ significantly, and the second gene is located on a secondary replicon. Such secondary-replicon genes either had duplicated and translocated from the chromosome in a distant past and had diverged significantly in sequence from the original gene, while retaining the function, or, more likely, they had entered the genome from an external source, possibly as a part of the replicon ancestor. rRNA operons are usually located in the first four replicons in terms of size. rRNAs encoded in the chromosome and in the secondary replicons may differ, even being closer to rRNAs of other strains than to paralogs from the same genome. Again, horizontal transfer of entire replicons, containing rRNA genes, from a closely related organism is a likely explanation..

For most replicons, we were able to predict genes involved in the initiation of replication, as well as their binding motifs and the structure of loci where the replication begins. The ParAB proteins involved in the segregation of sister copies of replicons were identified and their binding sites *parS* were predicted. The same was done for the XerCD proteins involved in resolving dimers and their binding sites *dif*. Finally, the structure and relative arrangement of loci responsible for the replication and segregation of replicons was described.

All these functional elements, as well as the gene composition and the size of replicons, can be used to group replicons by evolutionary origin. Overall, groups obtained using different approaches agree with each other. Analysis of gene composition identified fusions or splits of secondary replicons.

The phylogenetic tree of the RepA proteins, involved in initiating plasmid replication, can also serve to group replicons. While few published studies had focused on the replication of *Azospirillum* spp., one of them described a relatively small *Azospirillum* plasmid as an iteron plasmid, controlled by the RepA protein (Vande Broek et al. 2000). Here, we expand the list of both known *repA* genes and replication origin structures and provide coordinates of candidate iterons and DnaA boxes. Although plasmids in alphaproteobacteria are often *repABC* plasmids (Petersen et al. 2009; Pinto et al. 2012), no such plasmids have been found in *Azospirillum*. The phylogeny reconstruction for the predicted RepA proteins yields a classification of *Azospirillum* replicons into ten major groups, and several replicons whose RepA sequences are not similar to the others. The grouping of replicons by RepA also aligns well with the grouping by gene composition, with a minor exception. By gene composition, Group 9 by RepA splits into two groups, λ and μ, the latter including also several more replicons with diverse RepA.

In addition to the RepA protein sequence, the RepA binding motif and the structure of the replication origin are also conserved in groups of secondary replicons. As a rule, the plasmid origin of replication in *Azospirillum* is represented by at least one dense group of three iterons and several more distant iterons. It can be assumed that the observed structure of *Azospirillum* origins strictly controls the number of copies in order to maintain the balance between the chromosome and the plasmids.

In *repABC* plasmids, replication and partition systems are strongly related and encoded in a replicon as one operon (Cervantes-Rivera et al. 2011; Pinto et al. 2012), whereas in *repA* plasmids and megaplasmids, the partition system is independent of the replication system. Thus, in *Azospirillum*, the sequence and location of the *parAB* operon relative to the origin of replication can serve to classify replicons. Indeed, *parAB* was not detected in all replicons, since, although candidates for *parA* were found in most replicons, in some cases a corresponding *parB* could not be identified. Hence these replicons likely use a different segregation mechanism, relying on proteins not belonging to the ParA and ParB families (Aylett and Löwe 2012; Jalal and Le 2020; Siguier et al. 2023). The ParAB proteins form nine similarity groups, most of which match the RepA similarity groups. Since the distance from *parAB* to *repA* appears to be stable within the ParAB families and is maintained in cases when *repA* are different while *parAB* are the same, it can be assumed that this distance is functionally significant. Natural configurations of *parABS* and replication origin were mainly studied in *Caulobacter crescentus* (a-proteobacteria), *Vibrio cholerae* (g-proteobacteria), and *Pseudomonas aeruginosa* (g-proteobacteria) and they have short distances between *oriC* and *parS* sites, at most 10 kbp (David et al. 2014; Lagage et al. 2016; Tran et al. 2018). In *Azospirillum*, we see much larger distances between *parAB* and *repA*; even in chromosomes, this distance is already 12 kb, and in secondary replicons, distances exceeding 100 kb have been observed. Experiments with moving *parS* across chromosomes from the origin to the terminus demonstrated that *parS* sites remained functional at distances exceeding 500 kbp (David et al. 2014; Lagage et al. 2016; Tran et al. 2018). Thus, *parABS* distant from the origin in *Azospirillum* replicons are likely functional. Why such localization of *parABS* is conserved in some replicon groups remains an open question.

Proteins involved in dimer resolution and their binding motifs in some cases, for example, in *E. coli* or *Vibrio* (Demarre et al. 2014; Castillo et al. 2017; Fournes et al. 2025), make it possible to distinguish between plasmids or primary and secondary replicons. Previously, *dif* sites for primary and secondary replicons were predicted for sixteen Alphaproteobacteria with secondary chromosomes (Kono et al. 2011). Our study identified *dif* sites for most replicons and they are rather unusual for Alphaproteobacteria. The *dif* motifs in strains of the small tree clade are highly similar to each other, both for large and relatively small replicons. In strains of the large clade, the *dif* motifs clearly distinguish chromosomes and Group 1 replicons (based on RepA or ParAB similarity), while in replicons of other groups, the *dif* motifs coincide.

In some cases, we also observed several representatives of the same system on a single replicon. In the fifth replicon of *A. argentinense* MTCC4035, there are two identical *repA* genes with a matching environment. In Group 3 (by RepA similarity), there are two *parAB* operons with different sequences on different distances from the *repA* gene. In *A. humicireducens* SgZ-5, a fragment was translocated from a secondary replicon to the chromosome. Situations of this kind may be the result of assembly errors or natural fusions (Ostermayer et al. 2025).

Overall, this study demonstrates that fine details of replication and segregation can be described in bacteria with multiple, large replicons by applying diverse comparative genomic techniques. Of course, the final answer will be provided by experimental validation of these predictions.

## Material & Methods

### Genomes and annotation

Twenty two complete genomes of *Azospirillum* available in `GenBank` at December of 2023 were analyzed (**Supplementary Table 1**). To annotate the genomes, a combination of `prokka` (Seemann 2014) and `prodigal` (Hyatt et al. 2010) tools was used as a module of the `PanACoTA` pipeline (Perrin and Rocha 2021). Completeness of the genomes was assessed using the list of essential bacterial domains (Rinke et al. 2013).

### Alignment and phylogenetic trees

Protein alignment was constructed using `MUSCLE` (Edgar 2004) with default parameters. Preliminary orthologous groups for the tree were constructed using `MMseqs2` (Steinegger and Söding 2017) with 0.6 similarity threshold, universal single-copy genes were selected and aligned (concatenated protein-induced nucleotide alignment), and the phylogenetic tree of *Azospirillum* strains was constructed using the `iqtree2` tool (Minh et al. 2020) with 1000 bootstraps by modules of the `PanACoTA` pipeline (Perrin and Rocha 2021). The tree was visualised using online `iTOL` (Letunic and Bork 2024).

### Orthologous groups

Construction of orthologous groups for detailed protein analysis was made with `OrthoFinder` (Emms et al. 2026).

### Identification of RepA, ParA, ParB, and FtsK proteins

Candidate RepA proteins were identified by `HMMER` search (Potter et al. 2018) using RepA (PF10134.16) and TrfA profiles (PF07042.18). In addition, `BLAST` search for RepA from *Azospirillum brasilense* (Vande Broek et al. 2000) and three plasmid replication proteins (AWJ92039.1, WP_149235675.1, WP_149233155.1) was used.

The *parAB* operon was identified by `HMMER` search (Potter et al. 2018) with the ParB HTH domain (PF17762.4) and ParA (PF10609.12) profiles (Mistry et al. 2021) against all predicted proteins. Among the hits, pairs of adjacent genes arranged sequentially in the order *parA* followed by *parB* were selected.

Candidate FtsK proteins were identified by `BLAST` search for FtsK from two distant *Azospirillum* strains (KAA1056527.1, WP_149165488.1). The complete orthogroup was taken as candidate FtsK proteins.

Protein trees were built with the `iqtree2` tool (Minh et al. 2020).

### Relationships between replicons

Similarity of replicons was defined either as the Jaccard index calculated for orthogroups without paralogs, 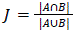, where A and B – genes from one and the other replicons, respectively, |*A*∩*B*| – the genes common to these replicons, and |*A*∪*B*| – genes found on at least one of the replicons, or as a generalisation of the Jaccard index, namely the ratio of the sum of the minimum numbers of paralogs in each orthogroup and the sum of the maximum numbers of paralogs in each orthogroup: 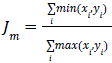, where 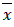 and 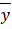 are the vectors composed of the number of paralogs for each orthogroup *i* represented in at least one of the compared replicons.

For each pair of genomes, the Jaccard coefficients and modified Jaccard coefficients were calculated for all pairs of replicons. Replicons were grouped according to bidirectional best hits in terms of the index used.

### Search for DNA motifs

For *de novo* prediction of DNA motifs, the `meme` suite was used (Bailey et al. 2015). To scan DNA for known DNA motifs a `python` application based on the `MOODS` library was used (GitHub – jhkorhonen/MOODS: MOODS: Motif Occurrence Detection Suite). Known XerCD-binding *dif* sites were taken from (Kono et al. 2011). Motif logos were constructed using the `WebLogo` generator (Crooks et al. 2004).

### Identification of KOPS sites

At the first step the motif GGGNAGGG was used to predict positions of KOPS. In *Azospirillum* chromosomes, sites with all four possible variants in the motif sequence were identified. Sites were accepted if they formed clusters with the density of at least one per 30 kbp and a pronounced direction skew over the replichores (**Supplementary Figure 11**). As a result, at the second step, the GGGCAGGG variant was selected as the motif.

To predict position of the KOPS direction change point in each replicon, for points at each 100 nucleotides the chi-square statistic value was calculated for the difference in the number of sites in quarter-replicons at each side of the point. The maximum of this statistic closest to the predicted *dif* was considered to be the change point for the KOPS direction.

To check that the change in the directionality of candidate sites at the replication terminus was not solely due to a combination of the GC-skew and the high G content of the motif, we also checked chromosomal positions of all motifs obtained by shuffling positions of the original motif (**Supplementary Table 4**). For the original motive, the chi-square statistic was the highest among all other motive options.

Similarly, the value of the chi-square statistic was calculated at the point corresponding to the position predicted by the *dif* sequence. *dif* position predicted by two methods were compared.

### Prediction of replication origins

To predict chromosomal replication origins, the `Ori-Finder2022` web service was used (Dong et al. 2022). The exact match of chromosomal *oriC* was also found in the `DoriC` database (ORI97017823) (Dong et al. 2023). Plasmid replication origins *oriV* were determined as an upstream region of *repA* gene with multiple RepA boxes and at least one DnaA box. The origin structures were drawn with the python library `DNA Features Viewer` (GitHub – Edinburgh-Genome-Foundry/DnaFeaturesViewer).

## Author statements

### Authors contribution

M.S.G. conceived the study. N.O.D. performed the study. N.O.D. and M.S.G. analyzed the results. N.O.D. drafted the paper. M.S.G. edited the paper.

### Ethical approval

Not applicable.

### Consent for publication

Not applicable.

### Data Availability Statement

All genome sequences analyzed in this study are publicly available in NCBI GenBank (see Supplementary Table 1 for accession numbers of 22 complete *Azospirillum* genomes and their replicons) (Benson et al. 2013). Other materials used are also publicly available; the record IDs and corresponding databases are provided in the Methods section. Supplementary materials are available at https://github.com/zaryanichka/Azospirillum.

## Acknowledgements

The study was initiated at the Summer Schools of Molecular and Theoretical Biology with Darina Karpova, Polina Pilipenko, Arina Tkacheva, Elizaveta Dialektova, and Agnia Ligay. The authors are grateful to Olga O. Bochkareva for inspiration and useful discussions.

## Supplementary materials

**Supplementary Table 1.** Genbank accessories and the sizes of the replicons for all the genomes used.

**Supplementary Table 2.** A summary table of information about the coordinates of the replication origin, the *parAB* operon, the distances between them, and the position of *dif*.

**Supplementary Table 3.** Comparison of chi-square statistics for predicting the position of *dif* based on the sequence and the KOPS direction change point.

**Supplementary Table 4.** Comparison of chi-square statistics for permuted versions of KOPS in chromosomes.

**Supplementary Figure 1**. The relation between the size of a replicon and its GC composition. The values for chromosomes are highlighted in orange, the values for the shortest replicons of each strain are highlighted in red, and the remaining dots are blue.

**Supplementary Figure 2**. Unrooted phylogenetic trees of (A) 5S, (B) 16S, and (C) 23S rRNA from rRNA operons. If there are non-monophyletic rRNAs from the same strain, they are marked in the same color, consistent between the trees.

**Supplementary Figure 3**. The phylogenetic tree of *Azospirillum* strains rooted on long branches with the best bidirectional hits between replicons marked by pink arrows and the best unidirectional hits marked by blue arrows. The replicons of each strain are numbered by size from the largest replicon (1), chromosome, to the smallest one.

**Supplementary Figure 4**. Schemes of replication origins for each group of replicons. The *repA* genes are shown as blue arrows, the DnaA boxes are shown as pink arrows, the RepA boxes are green arrows.

**Supplementary Figure 5**. Phylogenetic tree of the ParA proteins, rooted at the midpoint.

**Supplementary Figure 6**. Phylogenetic tree of the ParB proteins, rooted at the midpoint.

**Supplementary Figure 7**. The relationship between the size of the replicon and the distance from replicon origin to the start of the *parAB* operon across replicon similarity groups.

**Supplementary Figure 8**. Phylogenetic tree of site-specific XerC and XerD recombinases.

**Supplementary Figure 9**. KOPS positions for all replicons of the same group (by RepA). The color of height 1 ticks (lime and dark blue) indicates the direction of the site. The red height 2 tick indicates the position of *dif*. The short black tick indicates the maximum coordinate for the replicon.

**Supplementary Figure 10**. Cumulative GC skew, chi-square statistics for a sliding window with a step of 100, and the position of KOPS in both directions for all replicons of the strain, grouped by strain.

**Supplementary Figure 11**. KOPS positions in all chromosomes. The green and blue ticks of height 1 indicate KOPS in different directions. The pink and blue ticks of height 1.5 indicate the origin and terminus of replication. The red tick of height 2 indicates the position of the predicted *dif* site.

