## Supplementary material for "*Azospirillum*: effective plasmid management": AzospirillumSupplementaryFiguresAll.pdf

**Supplementary Figure 1.** The relation between the size of a replicon and its GC composition. The values for chromosomes are highlighted in orange, the values for the shortest replicons of each strain are highlighted in red, and the remaining dots are blue.

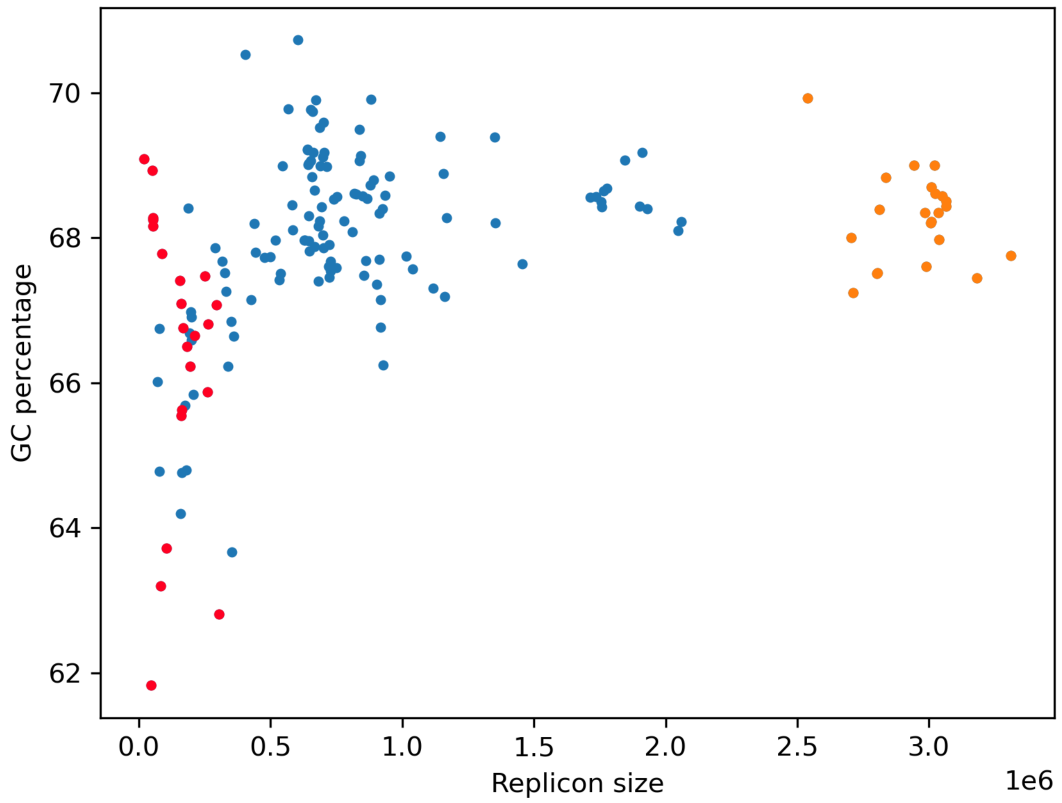

**Supplementary Figure 2.** Unrooted phylogenetic trees of (A) 5S, (B) 16S, and (C) 23S rRNA from rRNA operons. If there are non-monophyletic rRNAs from the same strain, they are marked in the same color, consistent between the trees.

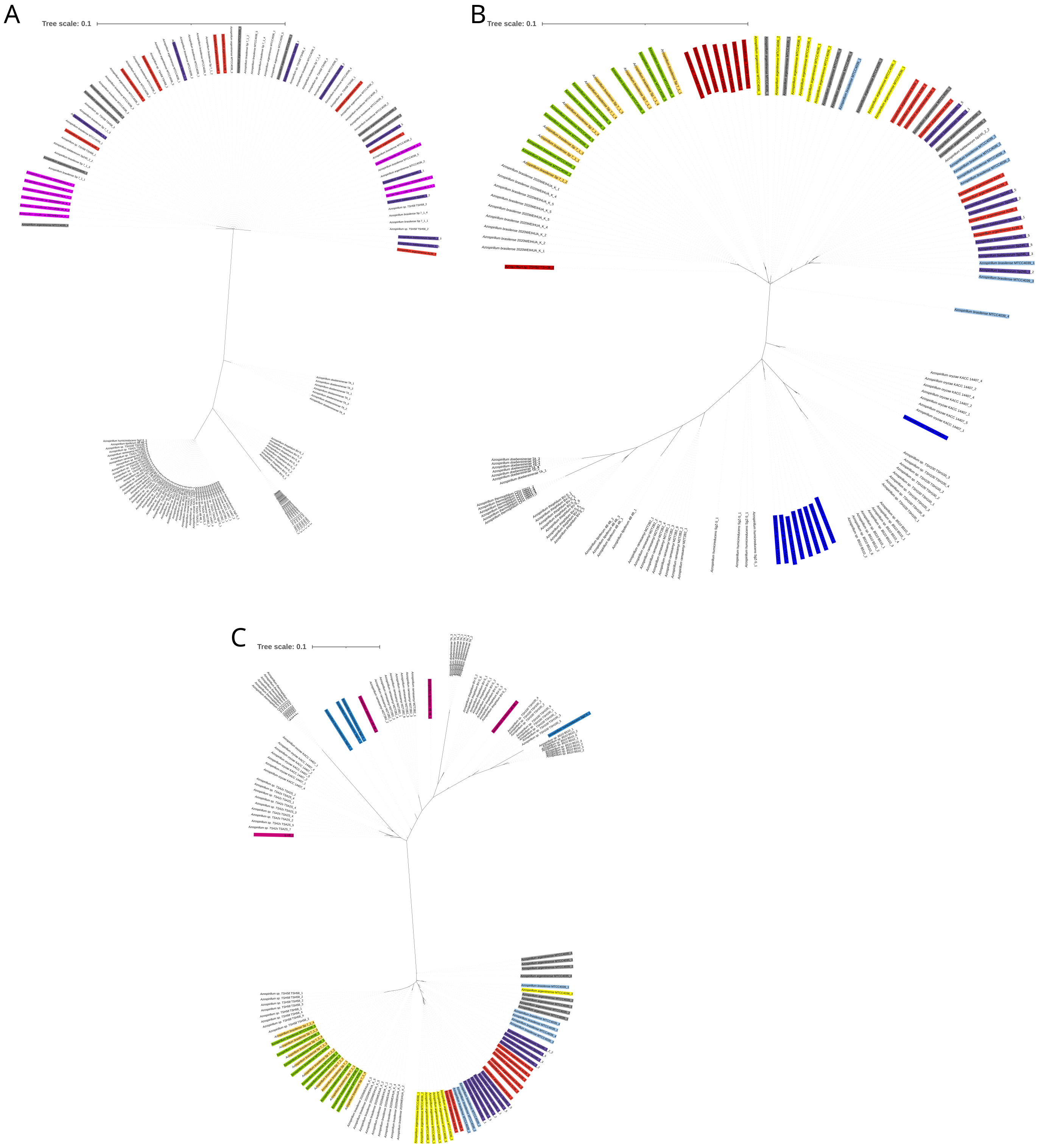

**Supplementary Figure 3.** The phylogenetic tree of Azospirillum strains rooted on long branches with the best bidirectional hits between replicons marked by pink arrows and the best unidirectional hits marked by blue arrows. The replicons of each strain are numbered by size from the largest replicon (1), chromosome, to the smallest one.

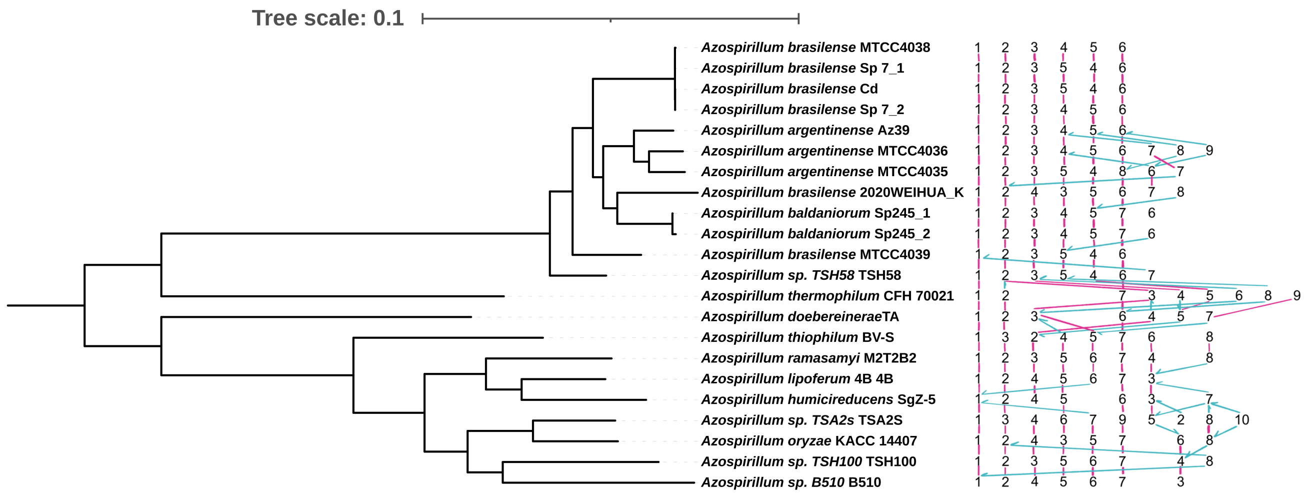

**Supplementary Figure 4.** Schemes of replication origins for each group of replicons. The repA genes are shown as blue arrows, the DnaA boxes are shown as pink arrows, the RepA boxes are green arrows.

Chromosome

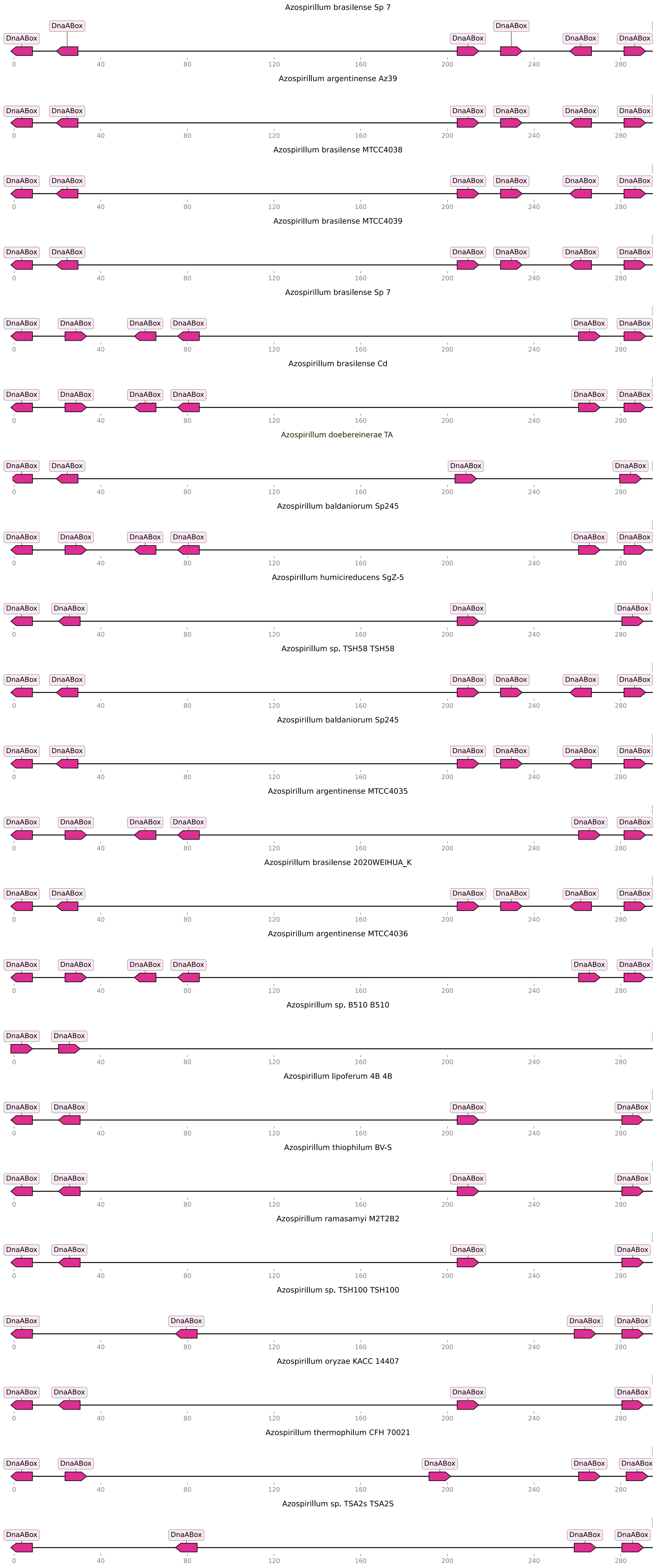

### Group1

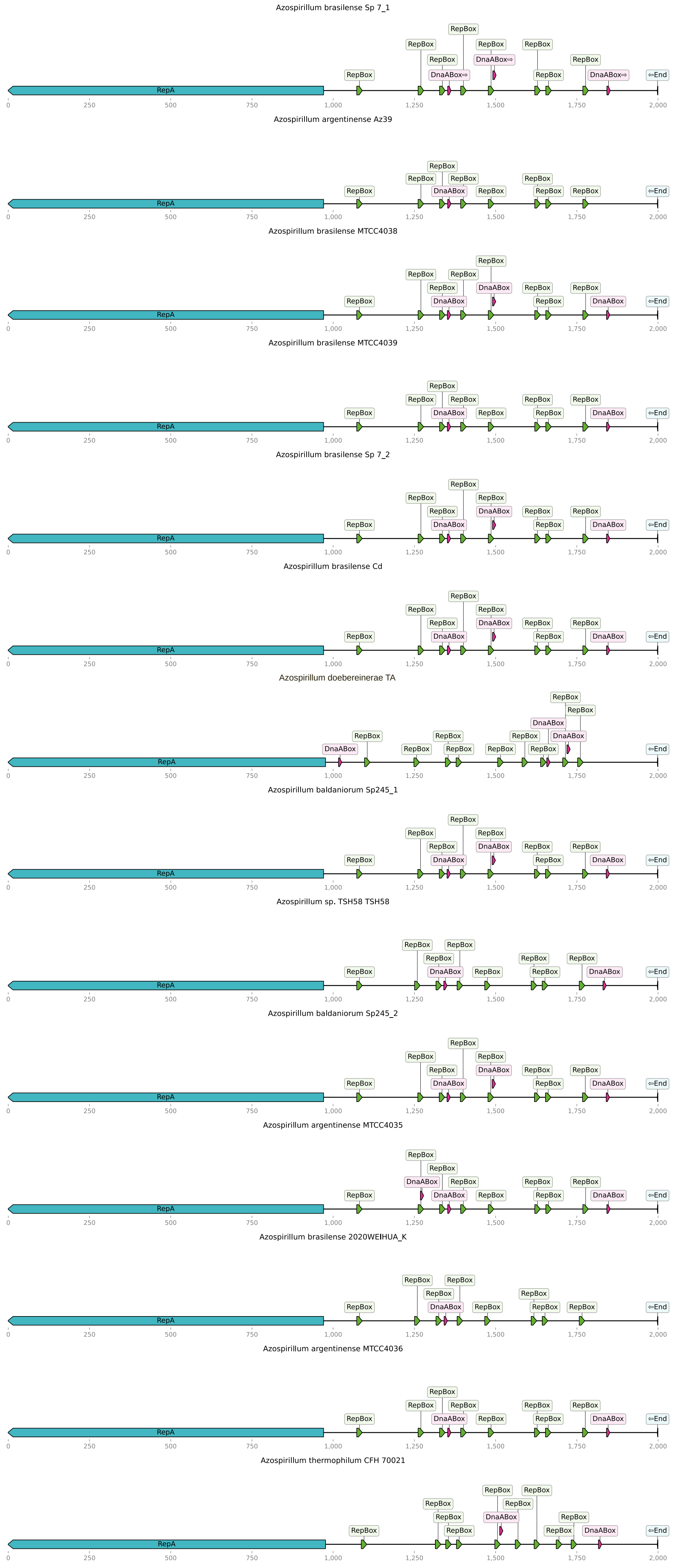
Azospirillum brasilense Sp 7_1
Azospirillum argentinense Az39
Azospirillum brasilense MTCC4038
Azospirillum brasilense MTCC4039
Azospirillum brasilense Sp 7_2
Azospirillum brasilense Cd
Azospirillum doebereinerae TA
Azospirillum baldaniorum Sp245_1
Azospirillum sp. TSH58 TSH58
Azospirillum baldaniorum Sp245_2
Azospirillum argentinense MTCC4035
Azospirillum brasilense 2020WEIHUA_K
Azospirillum argentinense MTCC4036
Azospirillum thermophilum CFH 70021
RepA
RepBox
DnaABox
⇦End
0
250
500
750
1,000
1,250
1,500
1,750
2,000

### Group2

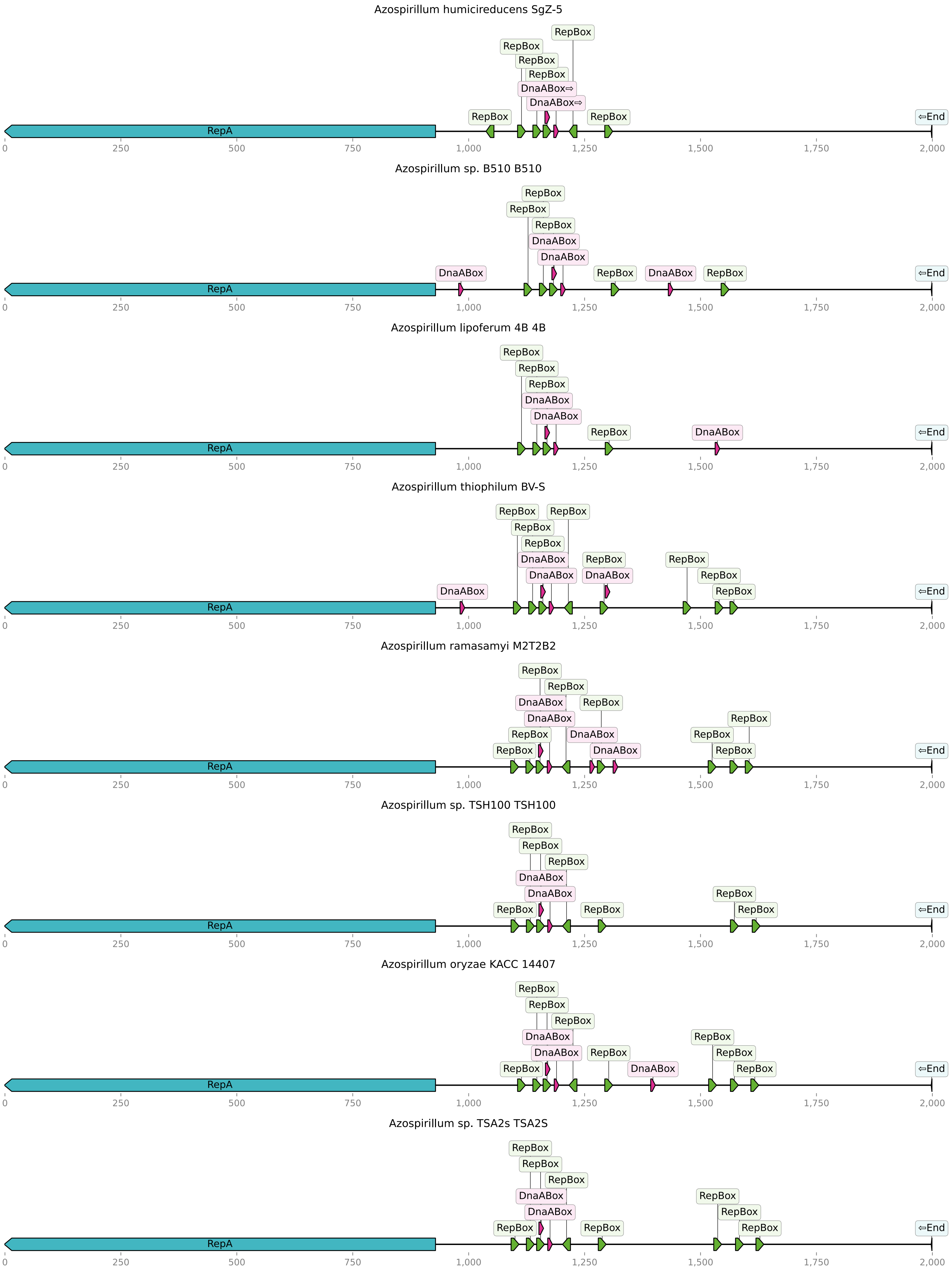
Azospirillum humicireducens SgZ-5
RepBox
DnaABox⇨
RepA
⇦End
0
250
500
750
1,000
1,250
1,500
1,750
2,000
Azospirillum sp. B510 B510
RepBox
DnaABox
RepA
⇦End
Azospirillum lipoferum 4B 4B
RepBox
DnaABox
RepA
⇦End
Azospirillum thiophilum BV-S
RepBox
DnaABox
RepA
⇦End
Azospirillum ramasamyi M2T2B2
RepBox
DnaABox
RepA
⇦End
Azospirillum sp. TSH100 TSH100
RepBox
DnaABox
RepA
⇦End
Azospirillum oryzae KACC 14407
RepBox
DnaABox
RepA
⇦End
Azospirillum sp. TSA2s TSA2S
RepBox
DnaABox
RepA
⇦End

### Group3

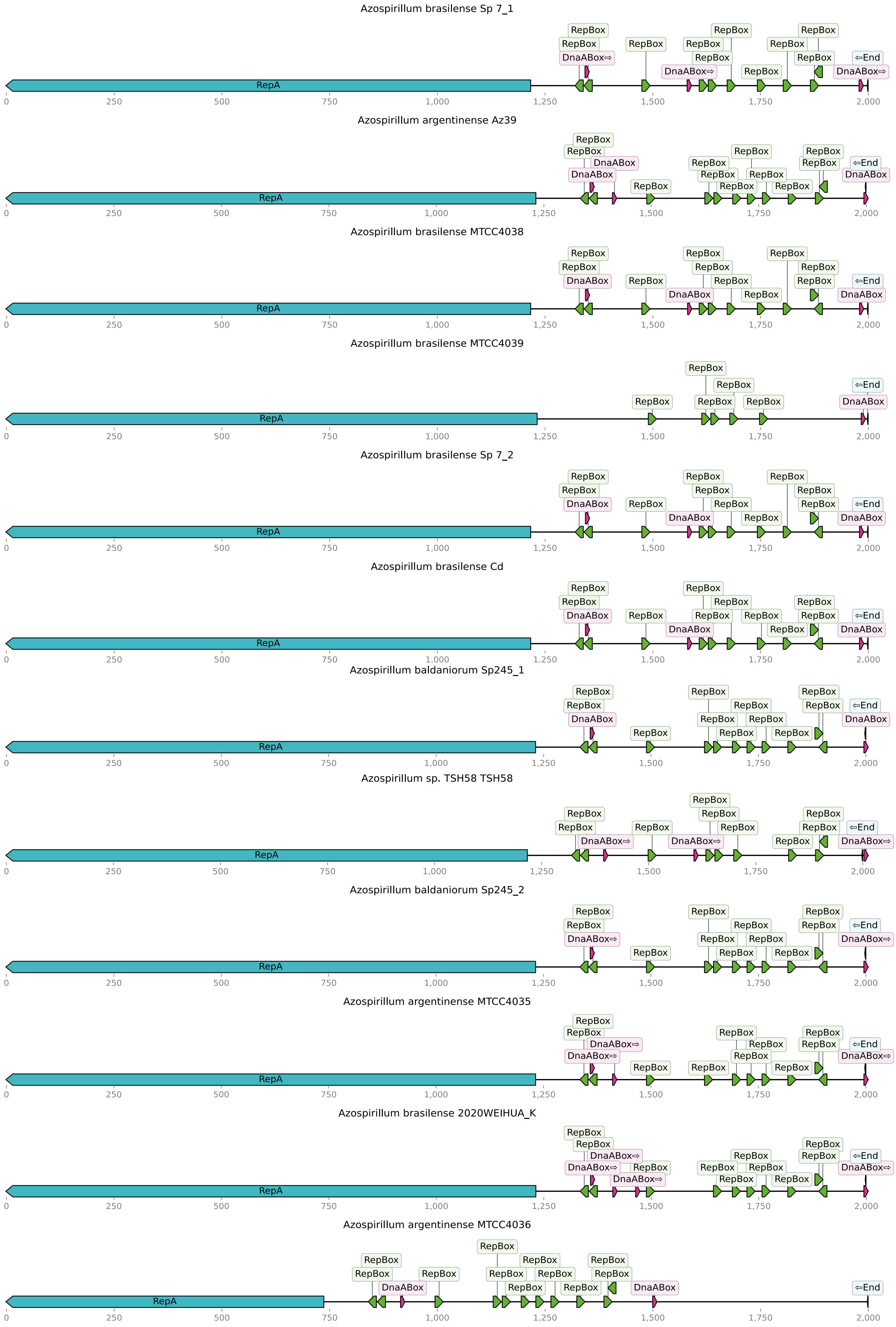

Azospirillum brasilense Sp 7_1
RepA
RepBox
DnaABox⇨
⇦End
0
250
500
750
1,000
1,250
1,500
1,750
2,000
Azospirillum argentinense Az39
RepA
RepBox
DnaABox
⇦End
Azospirillum brasilense MTCC4038
RepA
RepBox
DnaABox
⇦End
Azospirillum brasilense MTCC4039
RepA
RepBox
DnaABox
⇦End
Azospirillum brasilense Sp 7_2
RepA
RepBox
DnaABox
⇦End
Azospirillum brasilense Cd
RepA
RepBox
DnaABox
⇦End
Azospirillum baldaniorum Sp245_1
RepA
RepBox
DnaABox
⇦End
Azospirillum sp. TSH58 TSH58
RepA
RepBox
DnaABox⇨
⇦End
Azospirillum baldaniorum Sp245_2
RepA
RepBox
DnaABox⇨
⇦End
Azospirillum argentinense MTCC4035
RepA
RepBox
DnaABox⇨
⇦End
Azospirillum brasilense 2020WEIHUA_K
RepA
RepBox
DnaABox⇨
⇦End
Azospirillum argentinense MTCC4036
RepA
RepBox
DnaABox
⇦End

### Group4

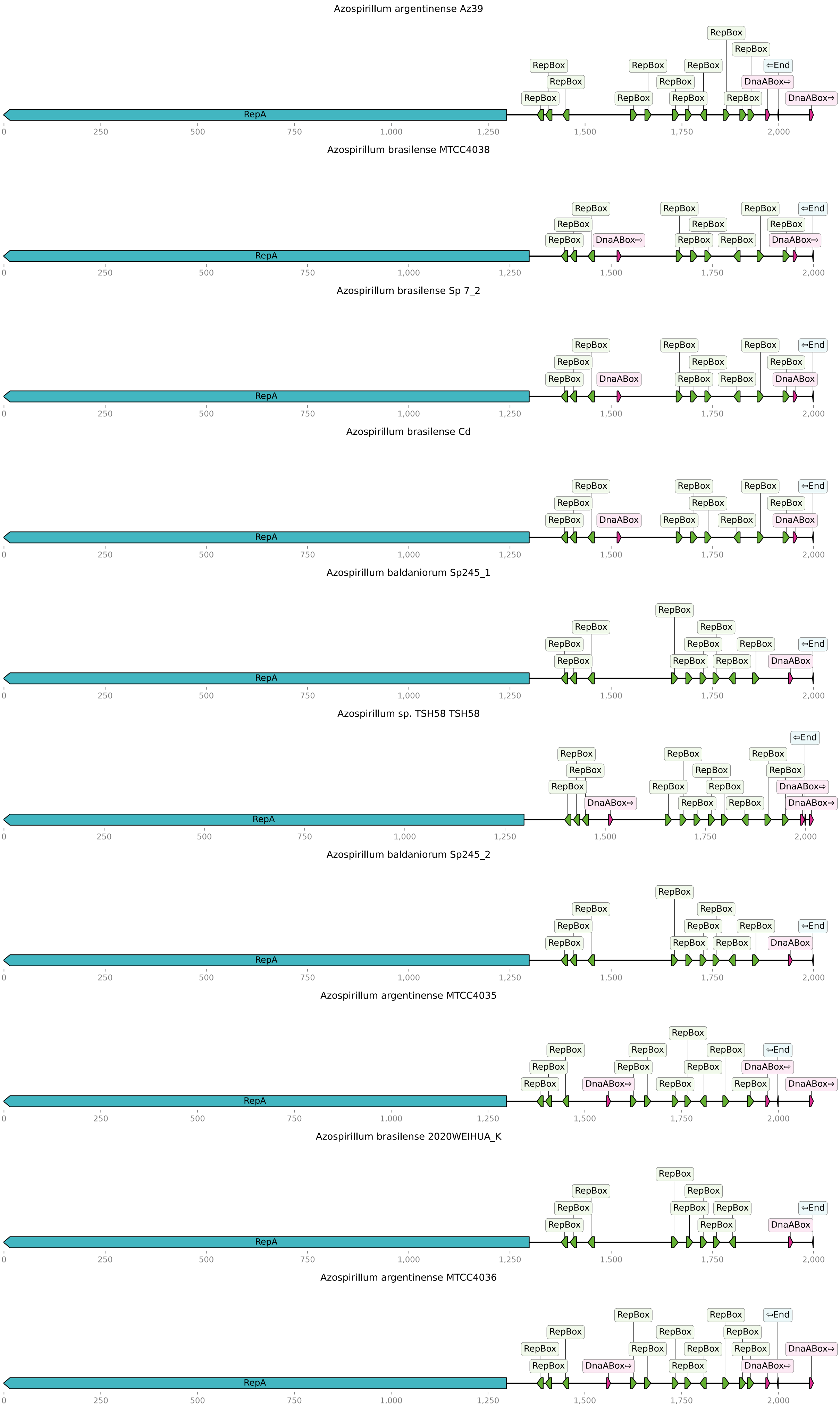

Azospirillum argentinense Az39
Azospirillum brasilense MTCC4038
Azospirillum brasilense Sp 7_2
Azospirillum brasilense Cd
Azospirillum baldaniorum Sp245_1
Azospirillum sp. TSH58 TSH58
Azospirillum baldaniorum Sp245_2
Azospirillum argentinense MTCC4035
Azospirillum brasilense 2020WEIHUA_K
Azospirillum argentinense MTCC4036
RepA
RepBox
DnaABox⇨
DnaABox
⇦End
0
250
500
750
1,000
1,250
1,500
1,750
2,000

### Group5

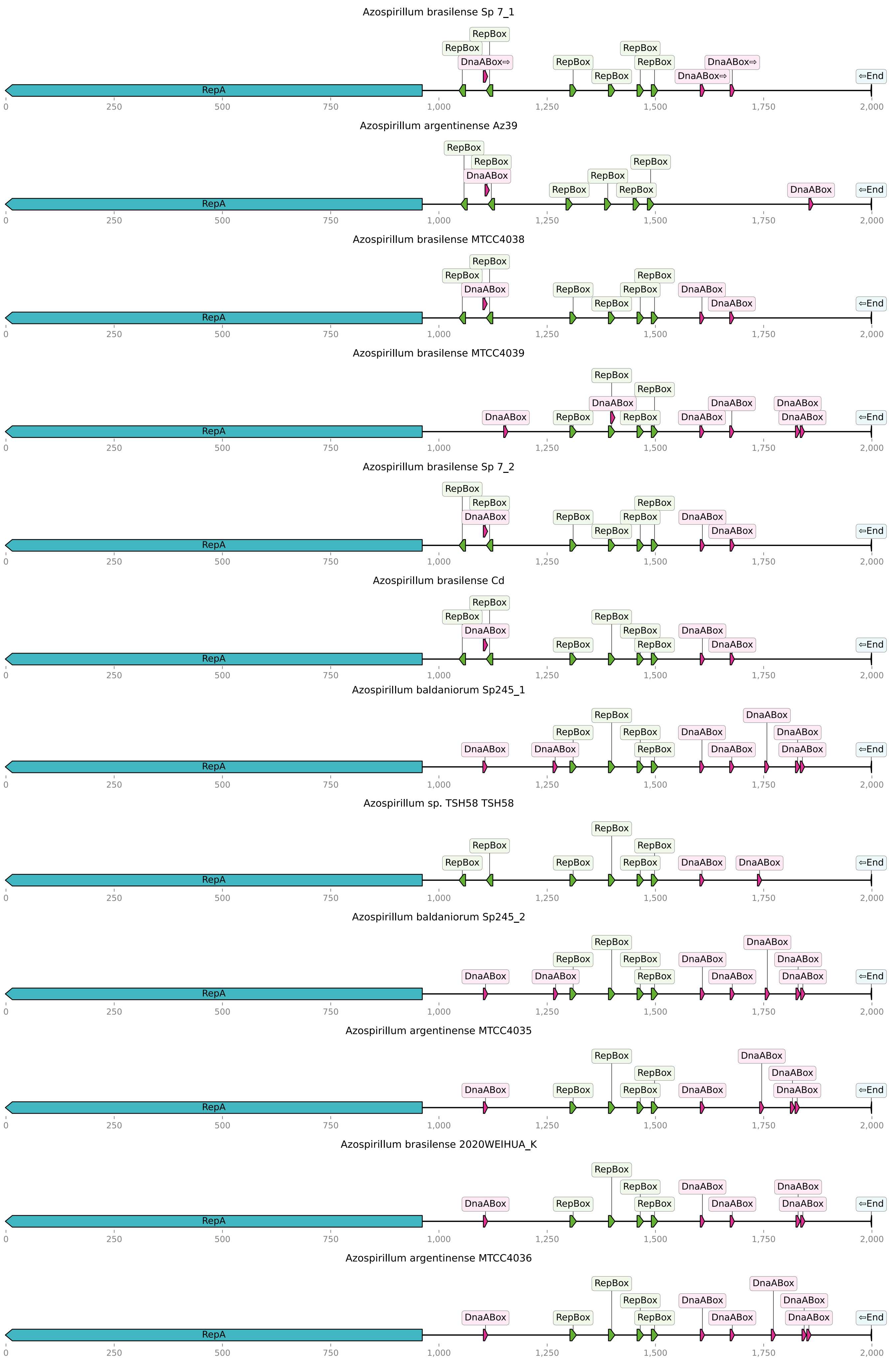

Azospirillum brasilense Sp 7_1
RepBox
RepBox
DnaABox⇨
RepBox
RepBox
RepBox
RepBox
RepBox
DnaABox⇨
DnaABox⇨
⇦End
RepA
0
250
500
750
1,000
1,250
1,500
1,750
2,000
Azospirillum argentinense Az39
RepBox
RepBox
DnaABox
RepBox
RepBox
RepBox
RepBox
DnaABox
⇦End
RepA
Azospirillum brasilense MTCC4038
RepBox
RepBox
DnaABox
RepBox
RepBox
RepBox
RepBox
DnaABox
DnaABox
⇦End
RepA
Azospirillum brasilense MTCC4039
RepBox
RepBox
DnaABox
DnaABox
RepBox
RepBox
DnaABox
DnaABox
DnaABox
DnaABox
⇦End
RepA
Azospirillum brasilense Sp 7_2
RepBox
RepBox
DnaABox
RepBox
RepBox
RepBox
RepBox
DnaABox
DnaABox
⇦End
RepA
Azospirillum brasilense Cd
RepBox
RepBox
DnaABox
RepBox
RepBox
RepBox
RepBox
DnaABox
DnaABox
⇦End
RepA
Azospirillum baldaniorum Sp245_1
RepBox
RepBox
RepBox
RepBox
DnaABox
DnaABox
DnaABox
DnaABox
DnaABox
DnaABox
DnaABox
⇦End
RepA
Azospirillum sp. TSH58 TSH58
RepBox
RepBox
RepBox
RepBox
RepBox
RepBox
DnaABox
DnaABox
⇦End
RepA
Azospirillum baldaniorum Sp245_2
RepBox
RepBox
RepBox
RepBox
DnaABox
DnaABox
DnaABox
DnaABox
DnaABox
DnaABox
DnaABox
⇦End
RepA
Azospirillum argentinense MTCC4035
RepBox
RepBox
RepBox
RepBox
DnaABox
DnaABox
DnaABox
DnaABox
DnaABox
⇦End
RepA
Azospirillum brasilense 2020WEIHUA_K
RepBox
RepBox
RepBox
RepBox
DnaABox
DnaABox
DnaABox
DnaABox
DnaABox
⇦End
RepA
Azospirillum argentinense MTCC4036
RepBox
RepBox
RepBox
RepBox
DnaABox
DnaABox
DnaABox
DnaABox
DnaABox
DnaABox
⇦End
RepA

### Group6

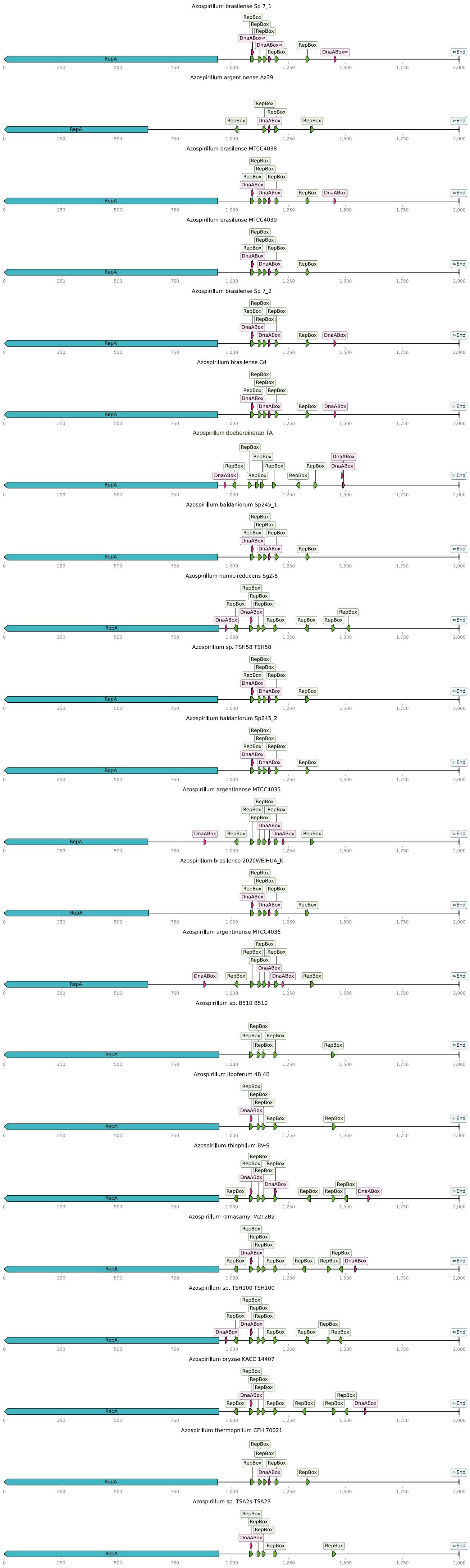
Azospirillum brasilense Sp 7_1
Azospirillum argentinense Az39
Azospirillum brasilense MTCC4038
Azospirillum brasilense MTCC4039
Azospirillum brasilense Sp 7_2
Azospirillum brasilense Cd
Azospirillum doebereinerae TA
Azospirillum baldaniorum Sp245_1
Azospirillum humicireducens SgZ-5
Azospirillum sp. TSH58 TSH58
Azospirillum baldaniorum Sp245_2
Azospirillum argentinense MTCC4035
Azospirillum brasilense 2020WEIHUA_K
Azospirillum argentinense MTCC4036
Azospirillum sp. B510 B510
Azospirillum lipoferum 4B 4B
Azospirillum thiophilum BV-S
Azospirillum ramasamyi M2T2B2
Azospirillum sp. TSH100 TSH100
Azospirillum oryzae KACC 14407
Azospirillum thermophilum CFH 70021
Azospirillum sp. TSA2s TSA2S
RepA
RepBox
DnaABox
⇐End
0
250
500
750
1,000
1,250
1,500
1,750
2,000

### Group7

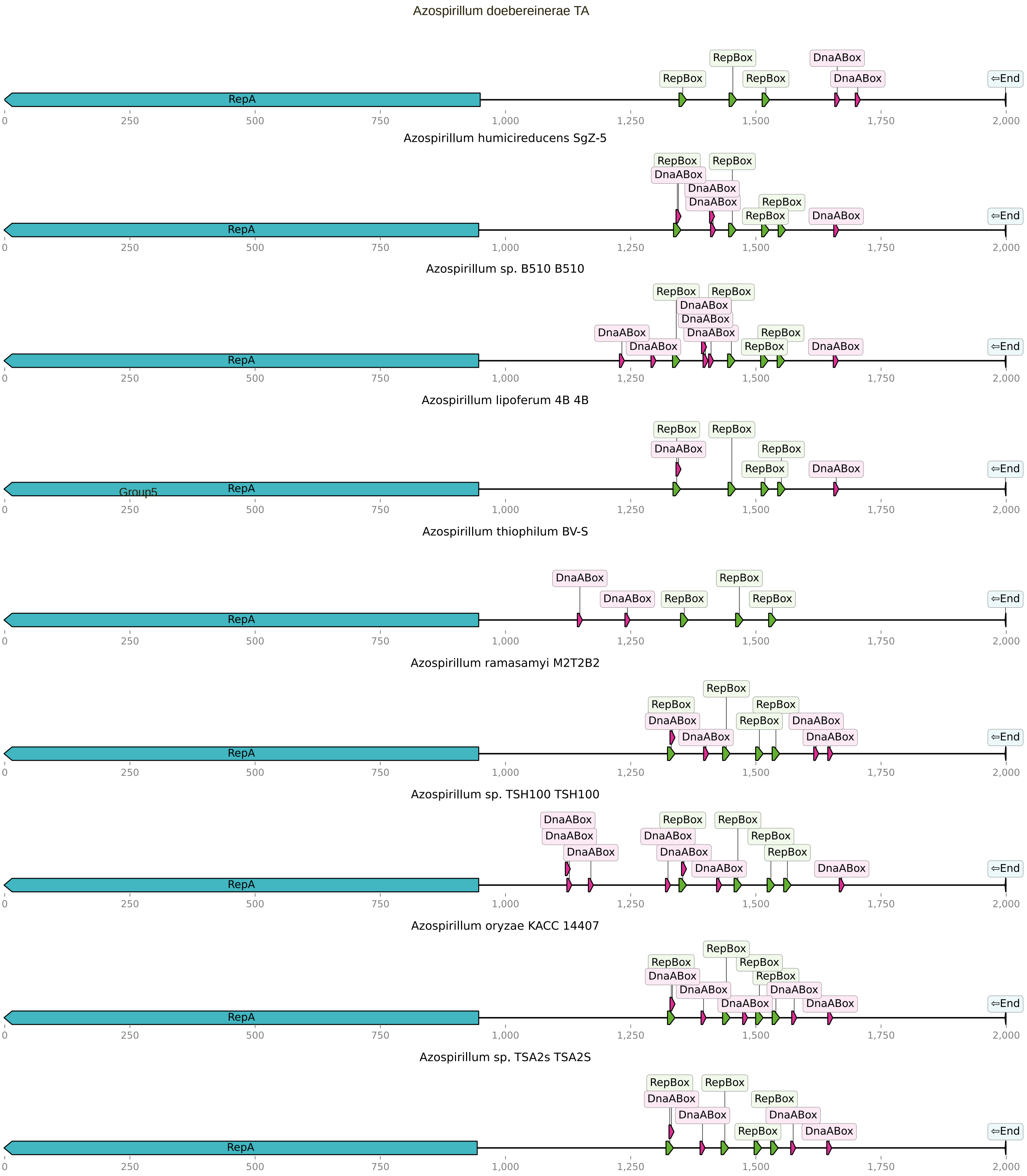

Azospirillum doebereinerae TA
RepA
RepBox
RepBox
RepBox
DnaABox
DnaABox
⇦End
0
250
500
750
1,000
1,250
1,500
1,750
2,000
Azospirillum humicireducens SgZ-5
RepA
RepBox
DnaABox
RepBox
DnaABox
DnaABox
RepBox
RepBox
DnaABox
⇦End
Azospirillum sp. B510 B510
RepA
DnaABox
DnaABox
RepBox
RepBox
DnaABox
DnaABox
DnaABox
RepBox
RepBox
DnaABox
⇦End
Azospirillum lipoferum 4B 4B
RepA
Group5
RepBox
DnaABox
RepBox
RepBox
RepBox
DnaABox
⇦End
Azospirillum thiophilum BV-S
RepA
DnaABox
DnaABox
RepBox
RepBox
RepBox
⇦End
Azospirillum ramasamyi M2T2B2
RepA
RepBox
DnaABox
DnaABox
RepBox
RepBox
RepBox
DnaABox
DnaABox
⇦End
Azospirillum sp. TSH100 TSH100
RepA
DnaABox
DnaABox
DnaABox
DnaABox
RepBox
DnaABox
DnaABox
RepBox
RepBox
RepBox
DnaABox
⇦End
Azospirillum oryzae KACC 14407
RepA
RepBox
DnaABox
DnaABox
RepBox
DnaABox
RepBox
RepBox
DnaABox
DnaABox
⇦End
Azospirillum sp. TSA2s TSA2S
RepA
RepBox
DnaABox
DnaABox
RepBox
RepBox
RepBox
DnaABox
DnaABox
⇦End

### Group8

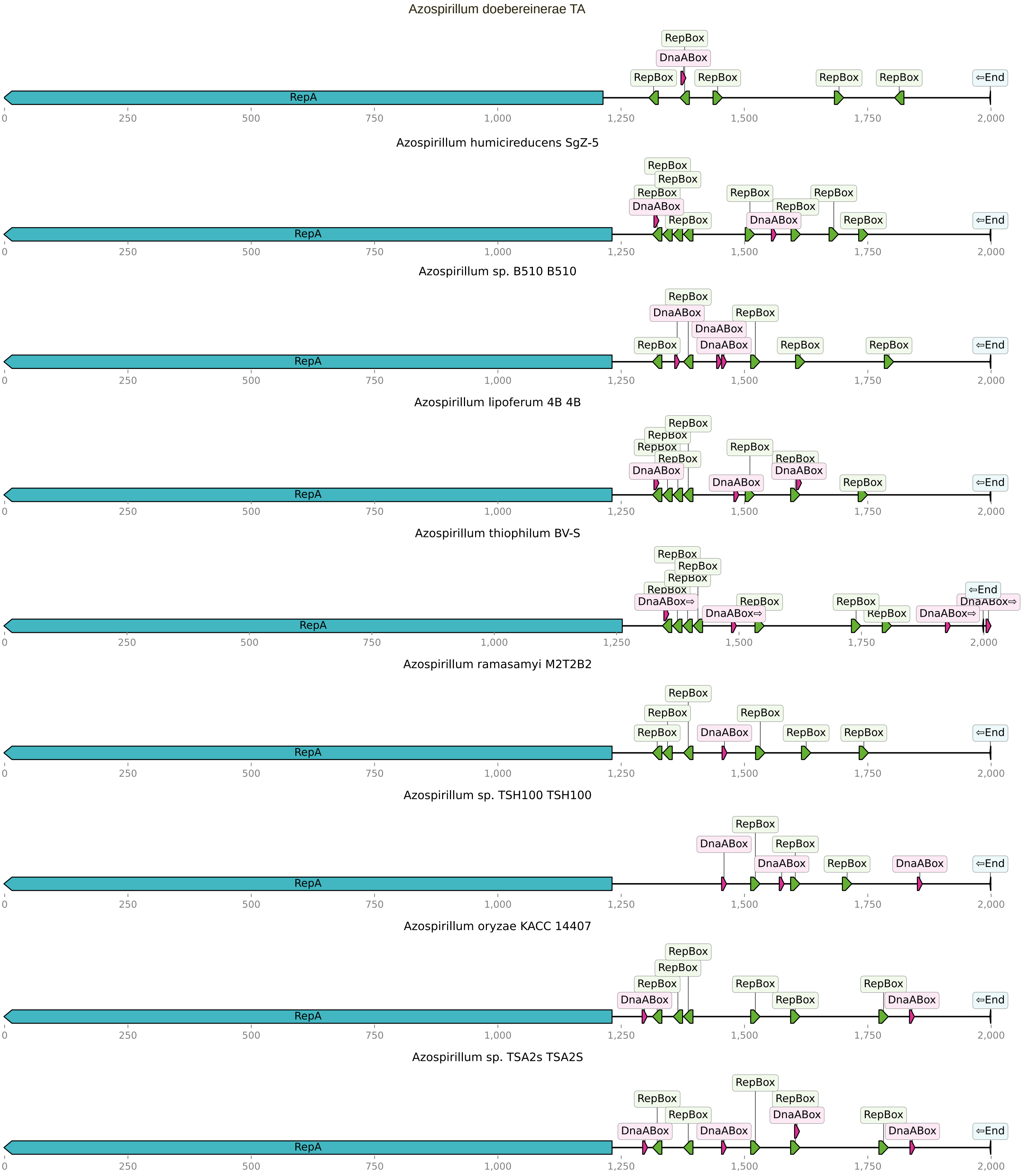
Azospirillum doebereinerae TA
RepA
RepBox
DnaABox
⇦End
Azospirillum humicireducens SgZ-5
Azospirillum sp. B510 B510
Azospirillum lipoferum 4B 4B
Azospirillum thiophilum BV-S
DnaABox⇨
Azospirillum ramasamyi M2T2B2
Azospirillum sp. TSH100 TSH100
Azospirillum oryzae KACC 14407
Azospirillum sp. TSA2s TSA2S
0
250
500
750
1,000
1,250
1,500
1,750
2,000

Group9

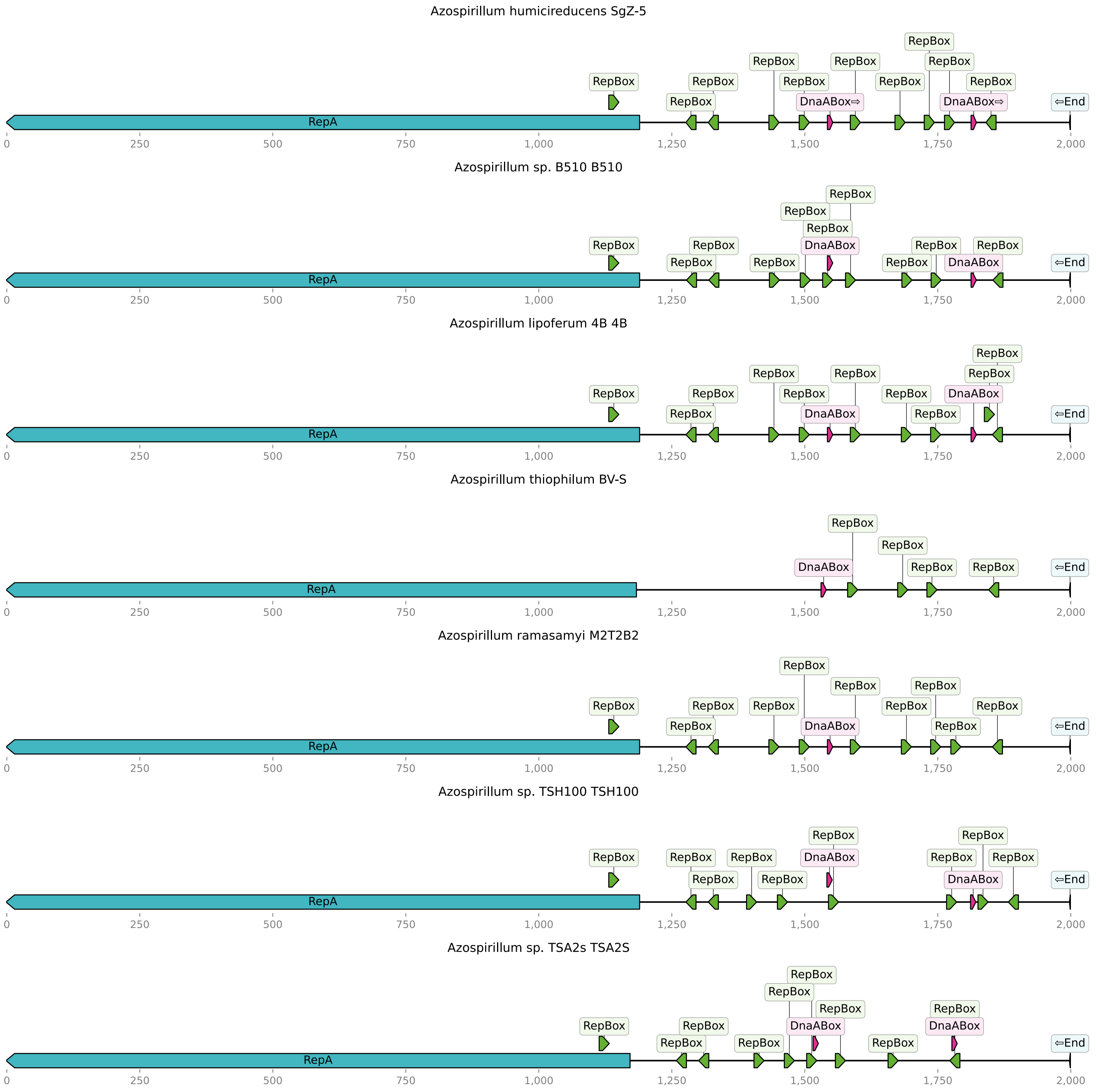
Azospirillum humicireducens SgZ-5
RepA
RepBox
DnaABox⇨
⇦End
0
250
500
750
1,000
1,250
1,500
1,750
2,000
Azospirillum sp. B510 B510
Azospirillum lipoferum 4B 4B
Azospirillum thiophilum BV-S
DnaABox
Azospirillum ramasamyi M2T2B2
Azospirillum sp. TSH100 TSH100
Azospirillum sp. TSA2s TSA2S

### Group10

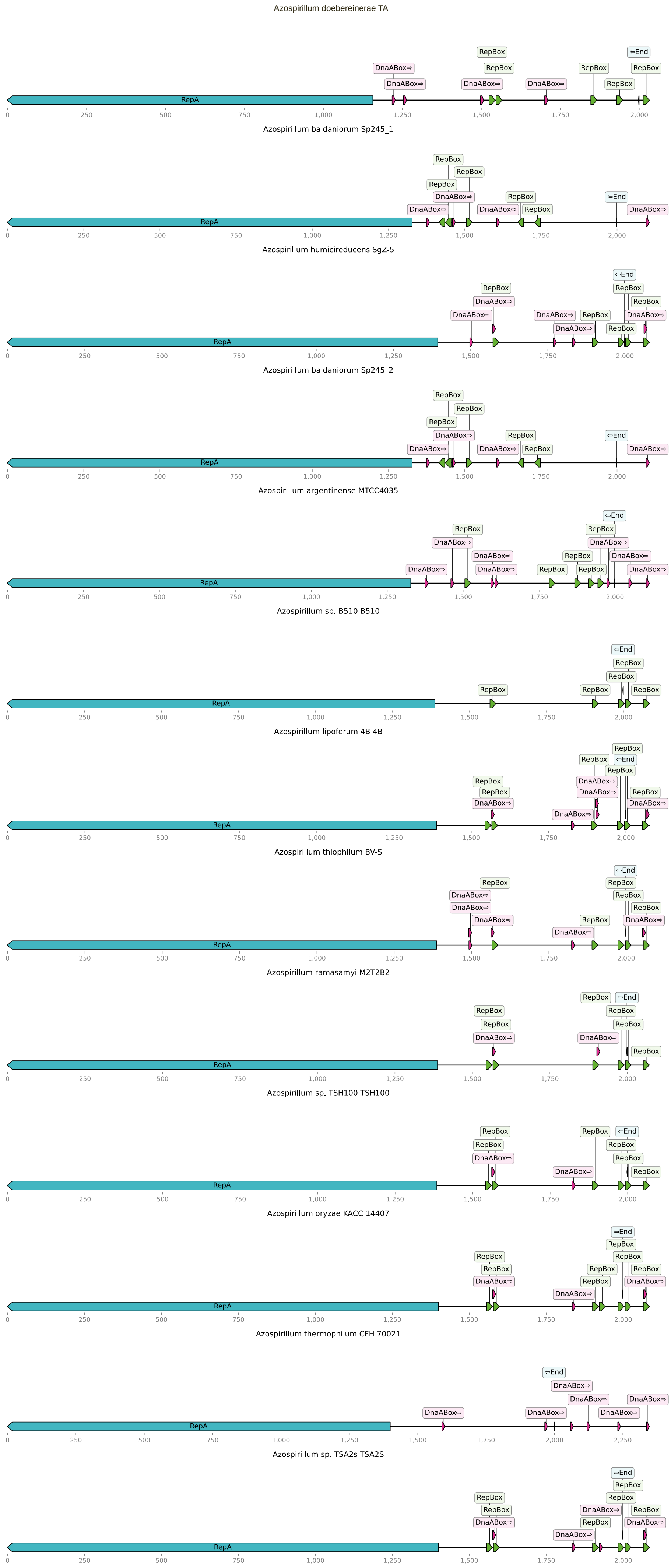

Azospirillum doebereinerae TA
Azospirillum baldaniorum Sp245_1
Azospirillum humicireducens SgZ-5
Azospirillum baldaniorum Sp245_2
Azospirillum argentinense MTCC4035
Azospirillum sp. B510 B510
Azospirillum lipoferum 4B 4B
Azospirillum thiophilum BV-S
Azospirillum ramasamyi M2T2B2
Azospirillum sp. TSH100 TSH100
Azospirillum oryzae KACC 14407
Azospirillum thermophilum CFH 70021
Azospirillum sp. TSA2s TSA2S

**Supplementary Figure 5.** Phylogenetic tree of the ParA proteins, rooted at the midpoint.

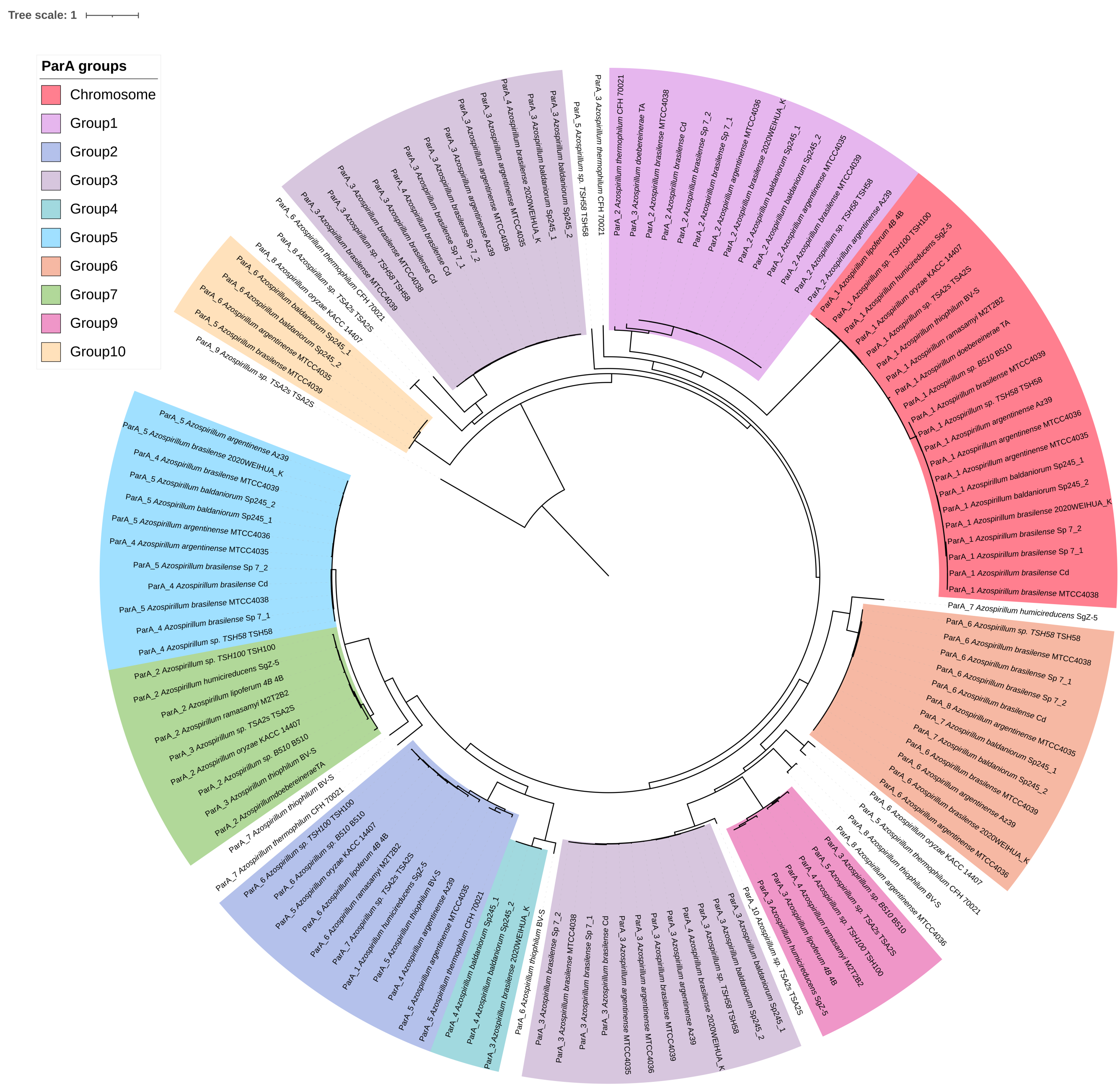

**Supplementary Figure 6.** Phylogenetic tree of the ParB proteins, rooted at the midpoint..

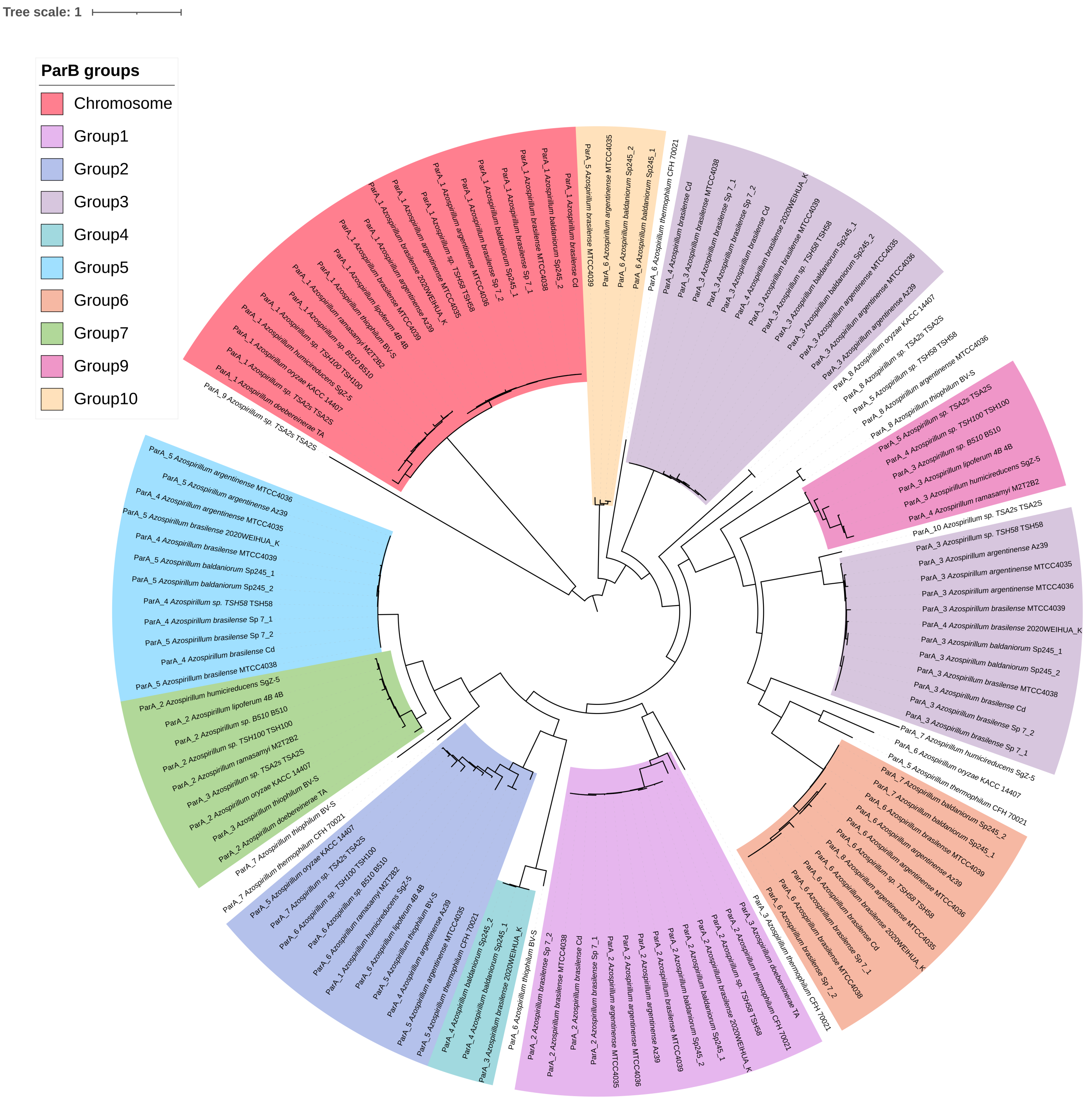

**Supplementary Figure 7.** The relationship between the size of the replicon and the distance from replicon origin to the start of the parAB operon across replicon similarity groups.

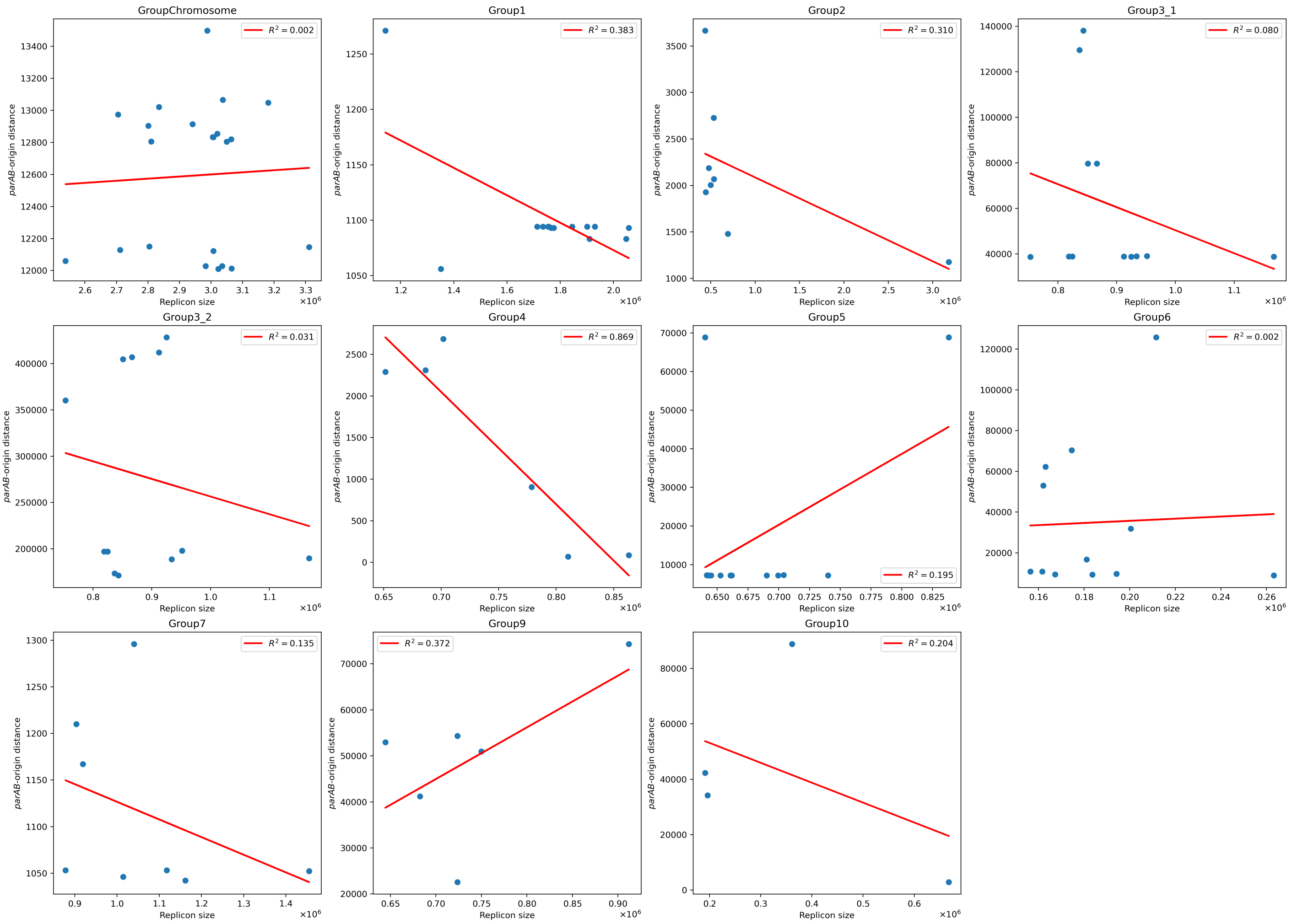

**Supplementary Figure 8.** Phylogenetic tree of site-specific XerC and XerD recombinases.

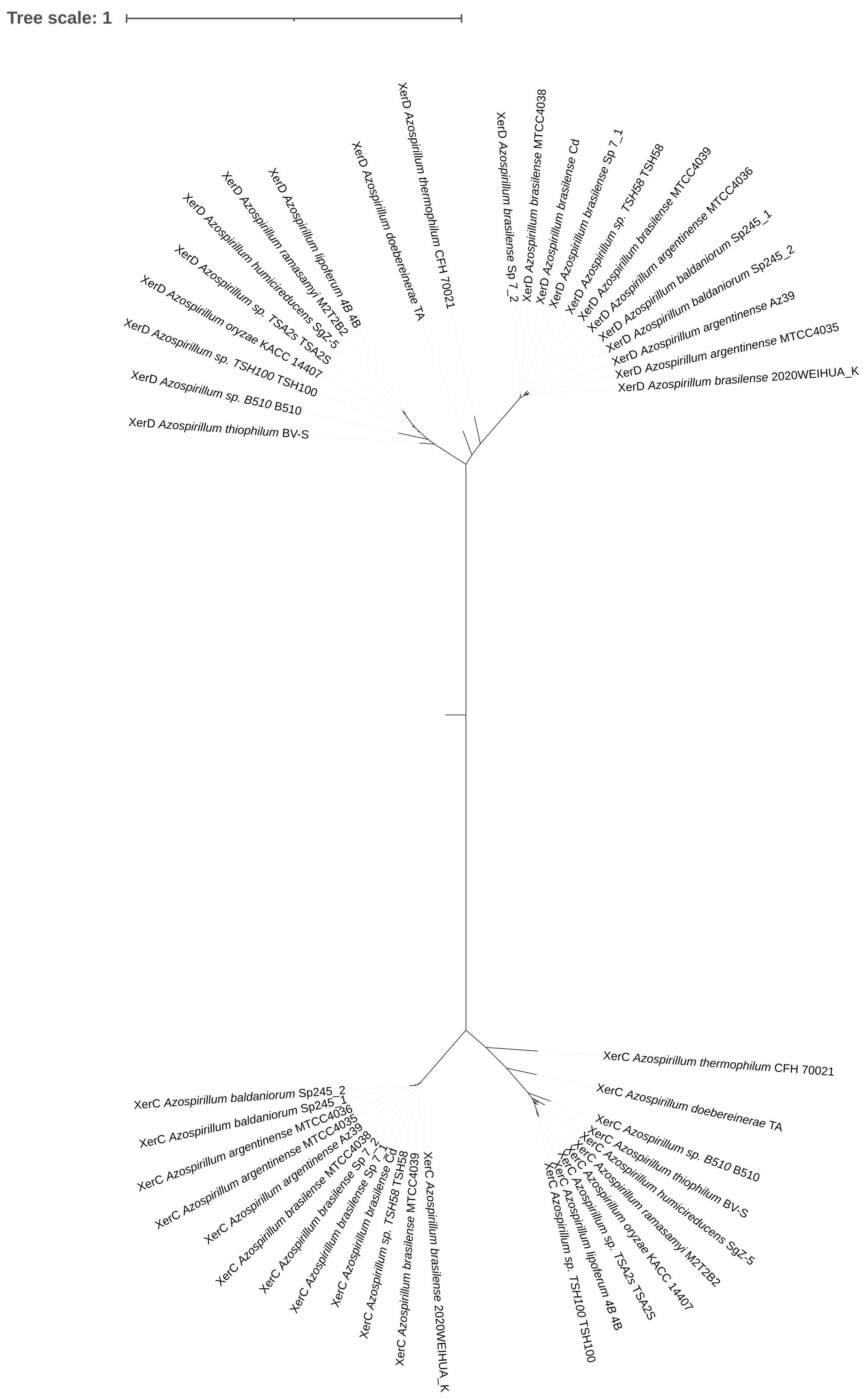

**Supplementary Figure 9.** KOPS positions for all replicons of the same group (by RepA). The color of height 1 ticks (lime and dark blue) indicates the direction of the site. The red height 2 tick indicates the position of dif. The short black tick indicates the maximum coordinate for the replicon.

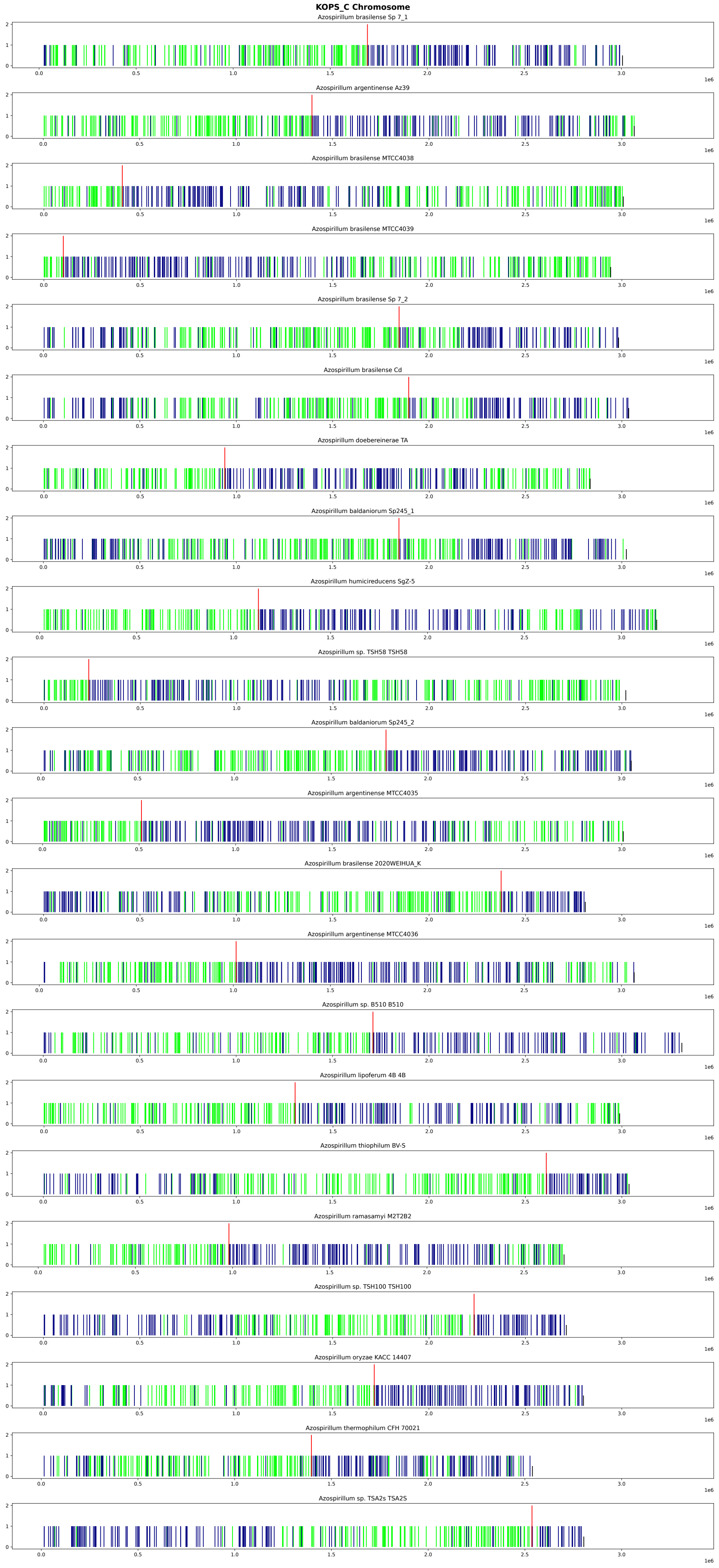

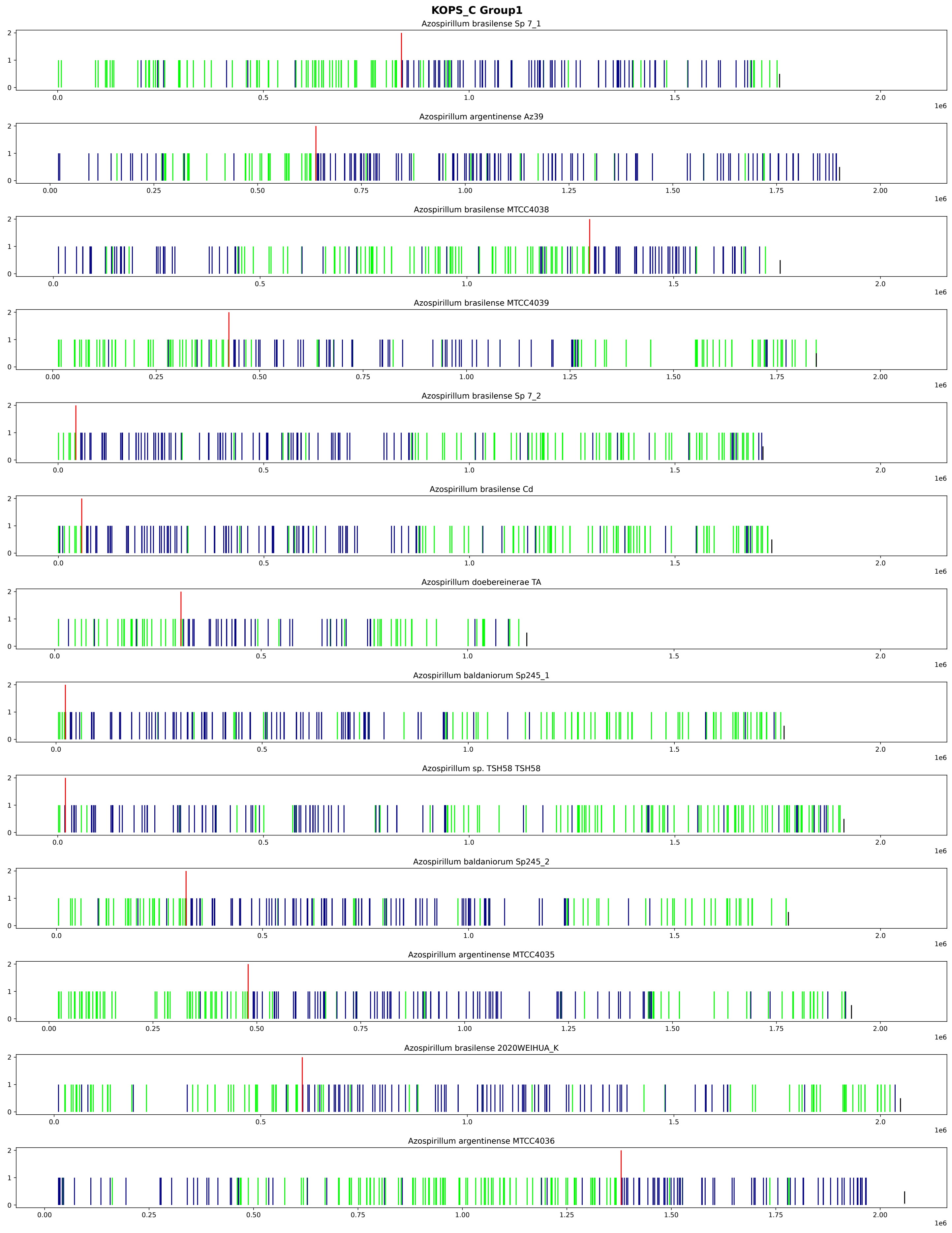
KOPS_C Group1
Azospirillum brasilense Sp 7_1
Azospirillum argentinense Az39
Azospirillum brasilense MTCC4038
Azospirillum brasilense MTCC4039
Azospirillum brasilense Sp 7_2
Azospirillum brasilense Cd
Azospirillum doebereinerae TA
Azospirillum baldaniorum Sp245_1
Azospirillum sp. TSH58 TSH58
Azospirillum baldaniorum Sp245_2
Azospirillum argentinense MTCC4035
Azospirillum brasilense 2020WEIHUA_K
Azospirillum argentinense MTCC4036
1e6

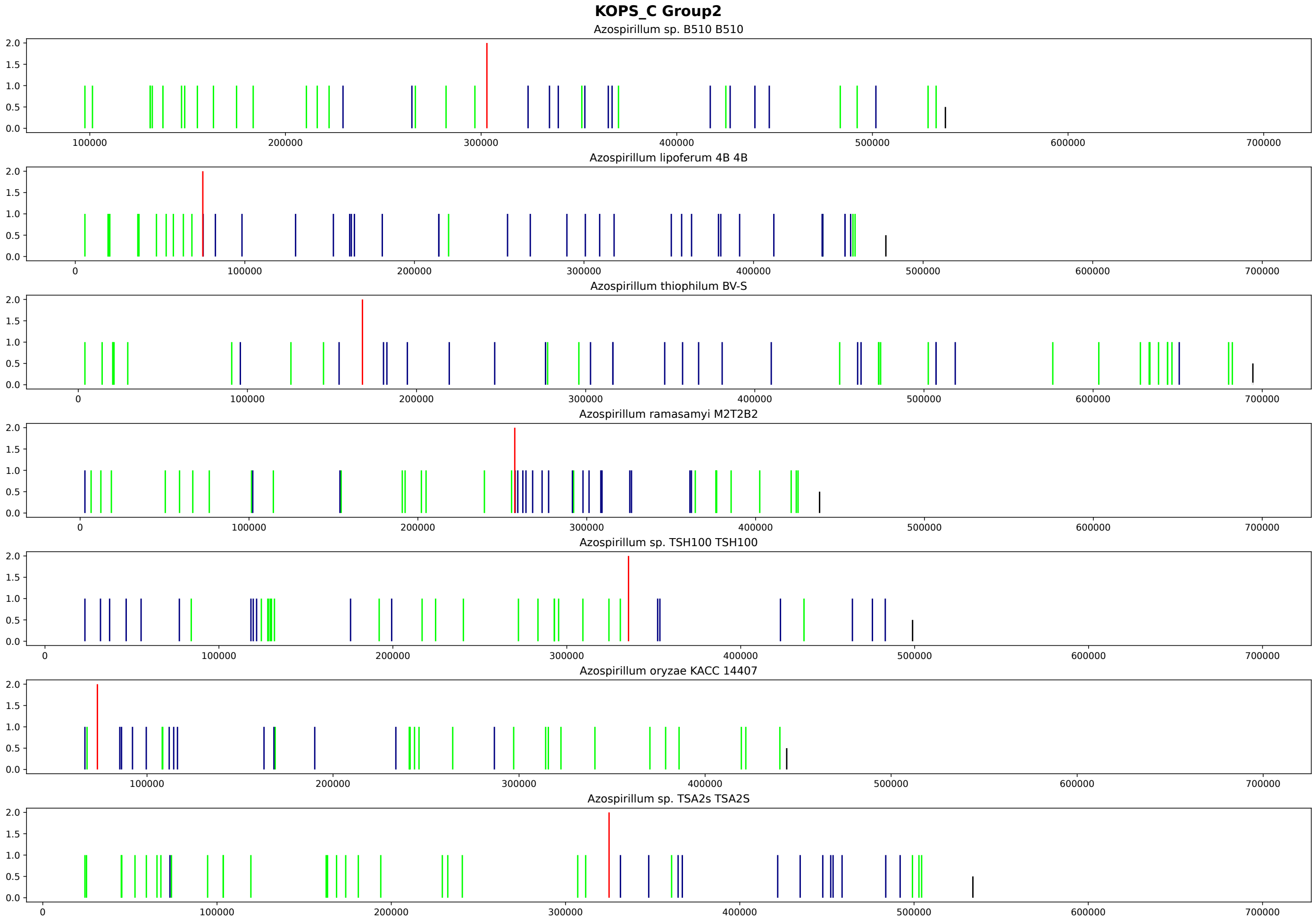
KOPS_C Group2
Azospirillum sp. B510 B510
Azospirillum lipoferum 4B 4B
Azospirillum thiophilum BV-S
Azospirillum ramasamyi M2T2B2
Azospirillum sp. TSH100 TSH100
Azospirillum oryzae KACC 14407
Azospirillum sp. TSA2s TSA2S

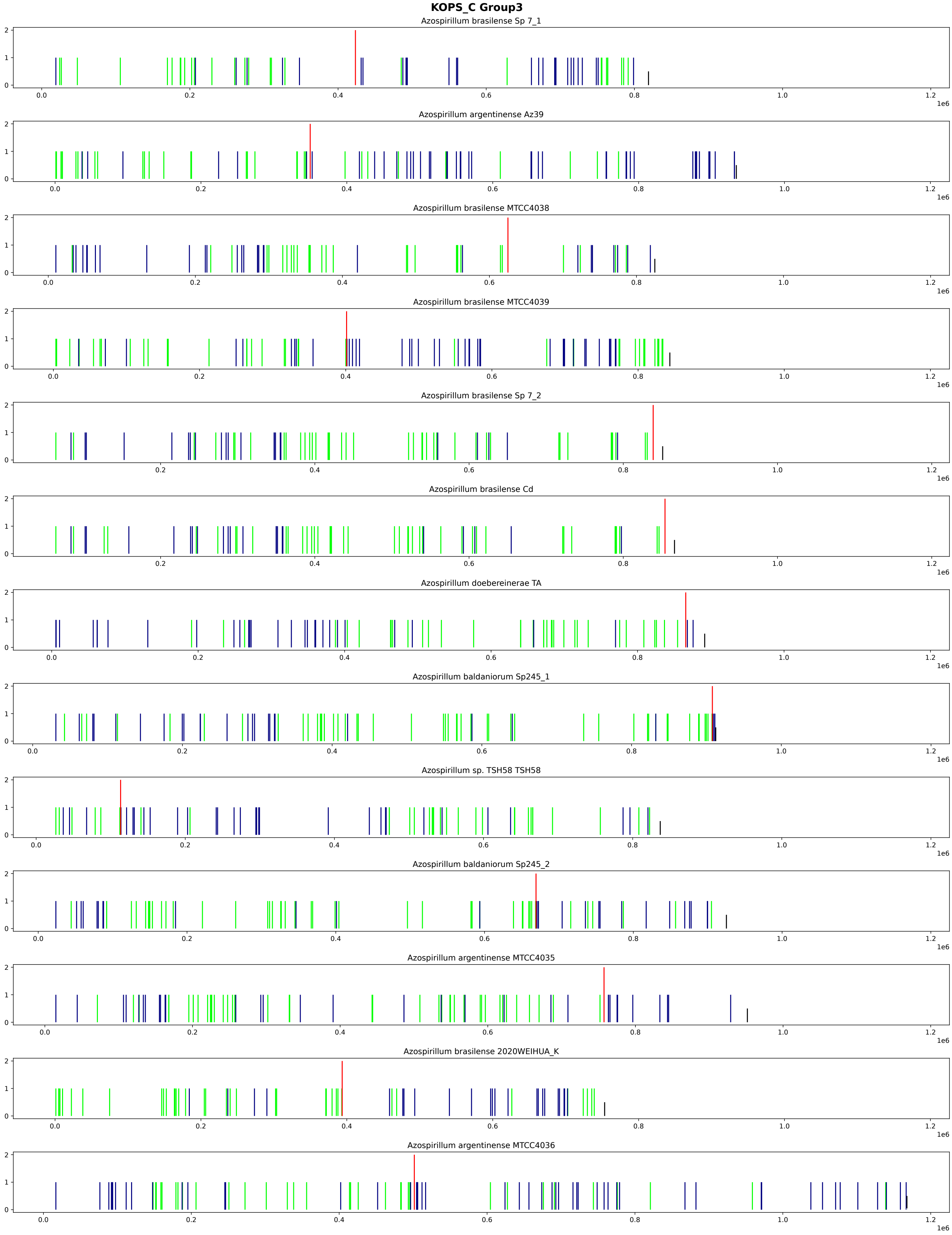

KOPS_C Group3
Azospirillum brasilense Sp 7_1
Azospirillum argentinense Az39
Azospirillum brasilense MTCC4038
Azospirillum brasilense MTCC4039
Azospirillum brasilense Sp 7_2
Azospirillum brasilense Cd
Azospirillum doebereinerae TA
Azospirillum baldaniorum Sp245_1
Azospirillum sp. TSH58 TSH58
Azospirillum baldaniorum Sp245_2
Azospirillum argentinense MTCC4035
Azospirillum brasilense 2020WEIHUA_K
Azospirillum argentinense MTCC4036
1e6

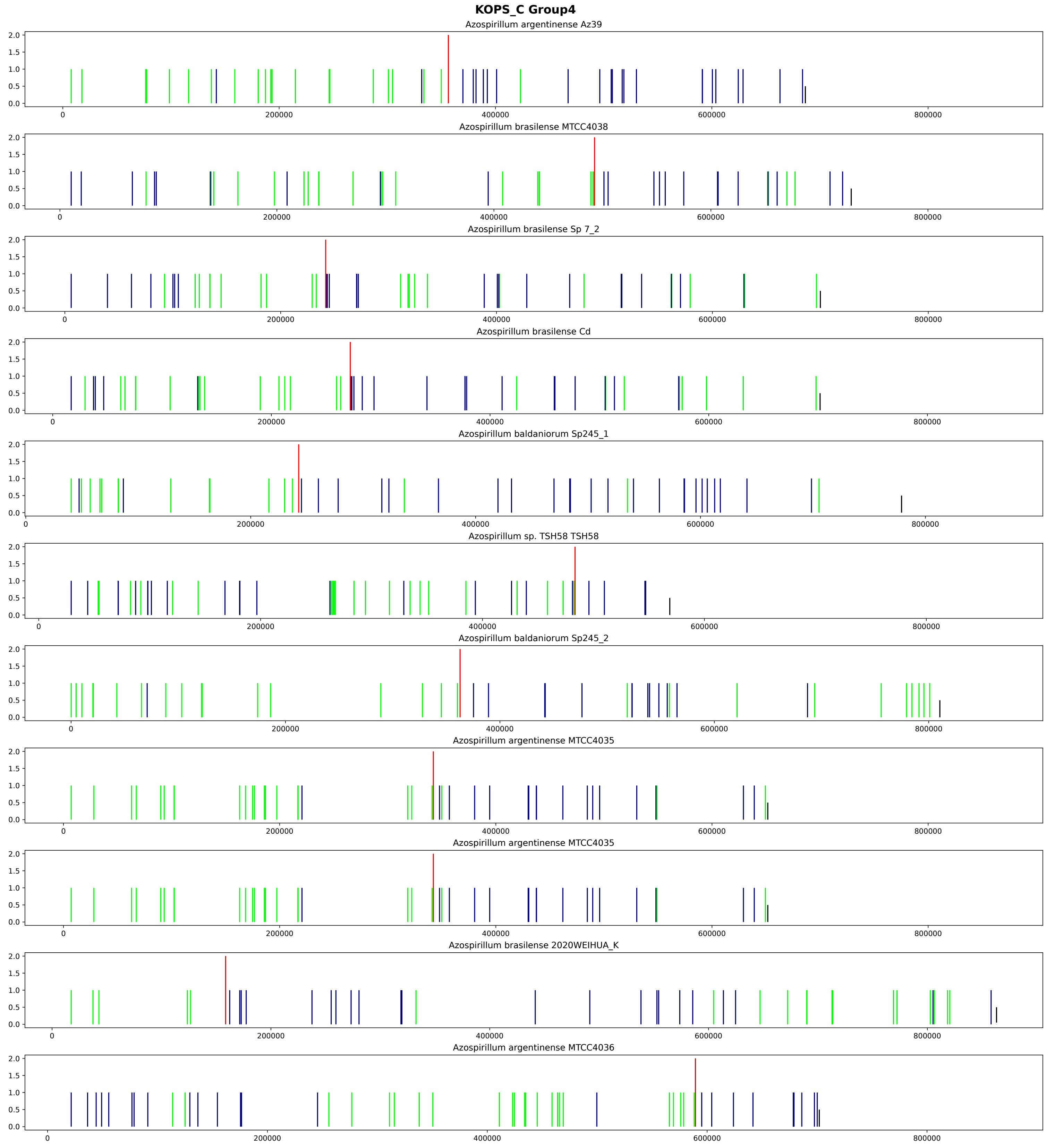
KOPS_C Group4
Azospirillum argentinense Az39
Azospirillum brasilense MTCC4038
Azospirillum brasilense Sp 7_2
Azospirillum brasilense Cd
Azospirillum baldaniorum Sp245_1
Azospirillum sp. TSH58 TSH58
Azospirillum baldaniorum Sp245_2
Azospirillum argentinense MTCC4035
Azospirillum argentinense MTCC4035
Azospirillum brasilense 2020WEIHUA_K
Azospirillum argentinense MTCC4036
0.0
0.5
1.0
1.5
2.0
0
200000
400000
600000
800000

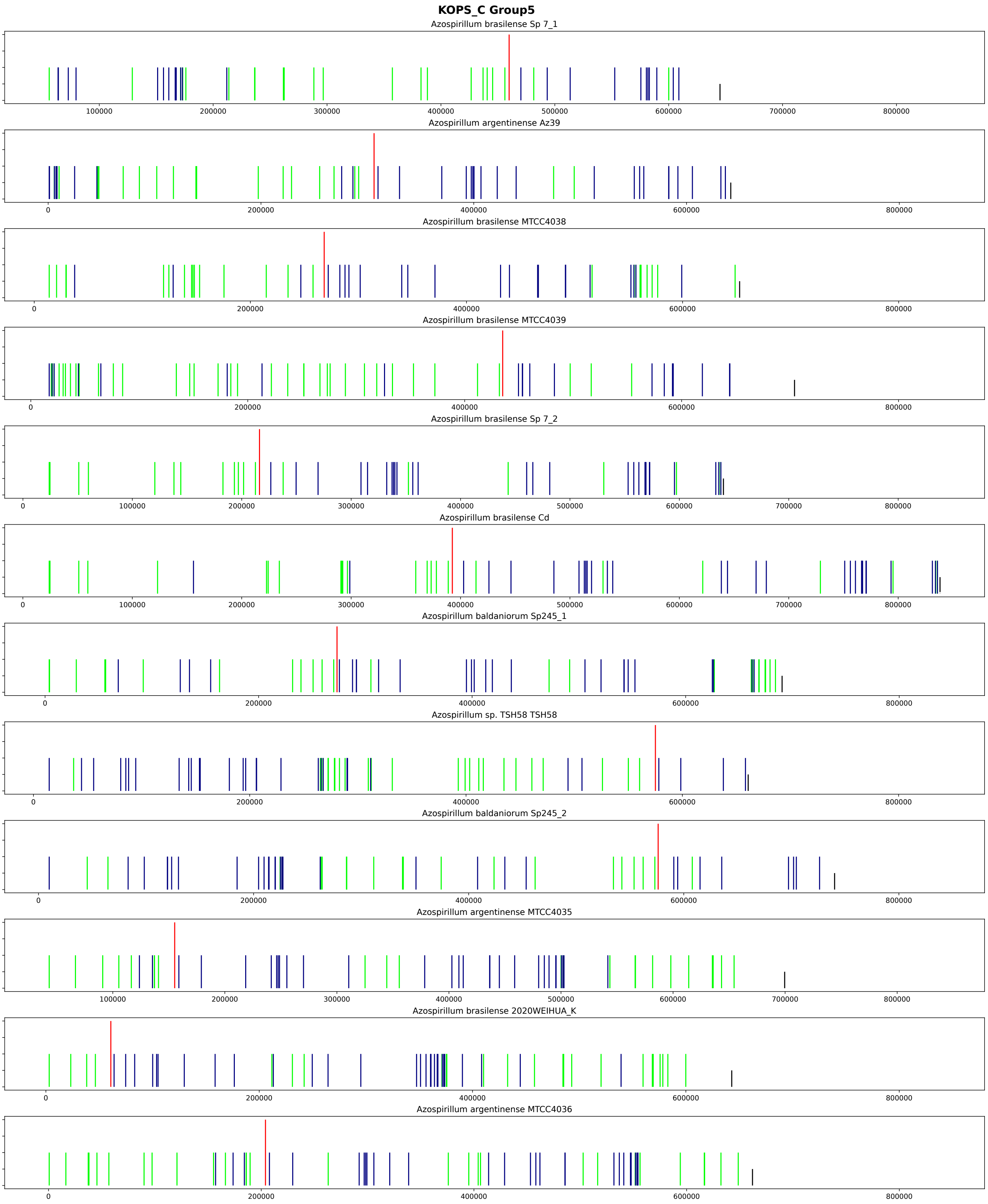
KOPS_C Group5
Azospirillum brasilense Sp 7_1
Azospirillum argentinense Az39
Azospirillum brasilense MTCC4038
Azospirillum brasilense MTCC4039
Azospirillum brasilense Sp 7_2
Azospirillum brasilense Cd
Azospirillum baldaniorum Sp245_1
Azospirillum sp. TSH58 TSH58
Azospirillum baldaniorum Sp245_2
Azospirillum argentinense MTCC4035
Azospirillum brasilense 2020WEIHUA_K
Azospirillum argentinense MTCC4036

#### KOPS_C Group6

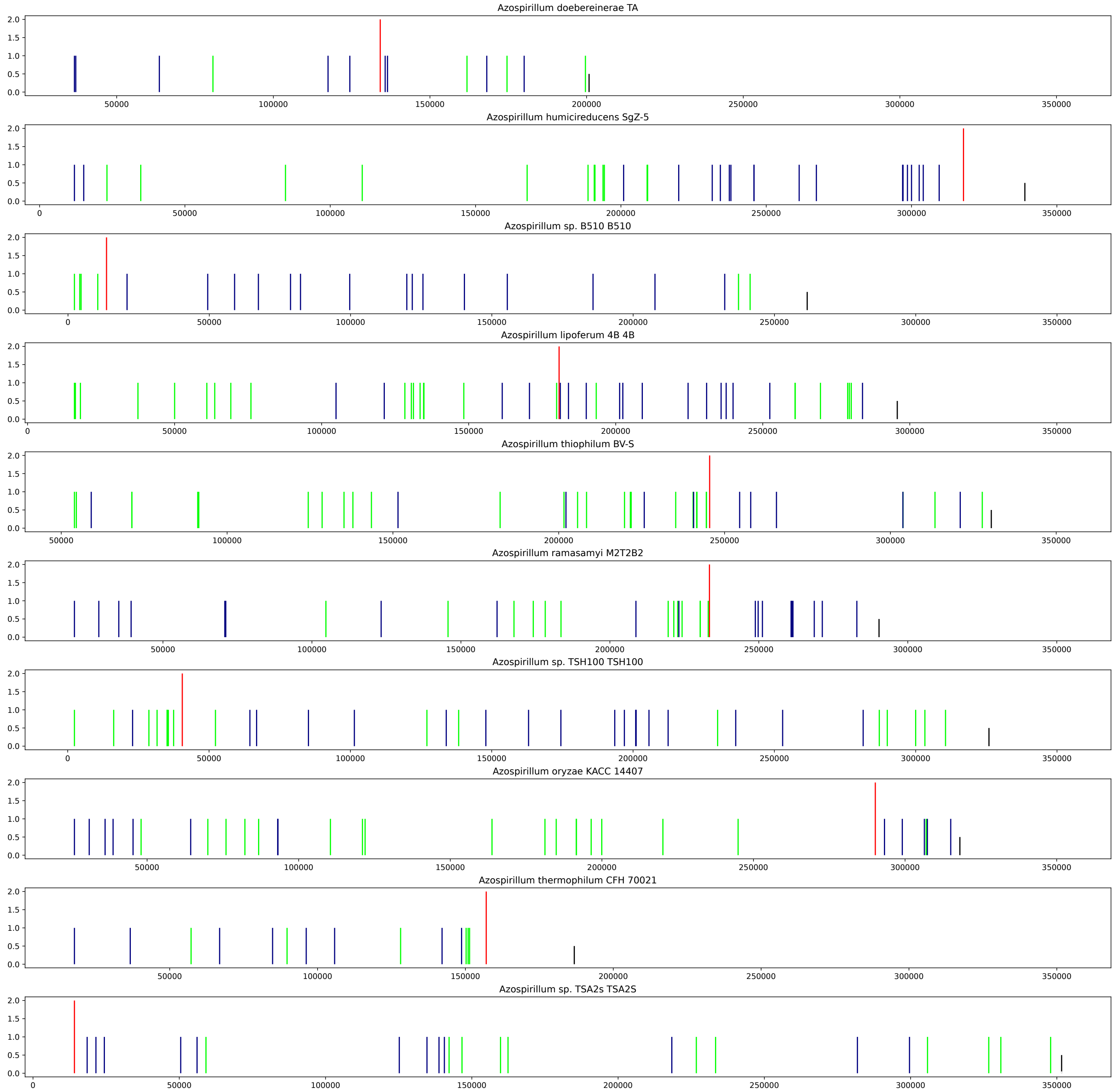

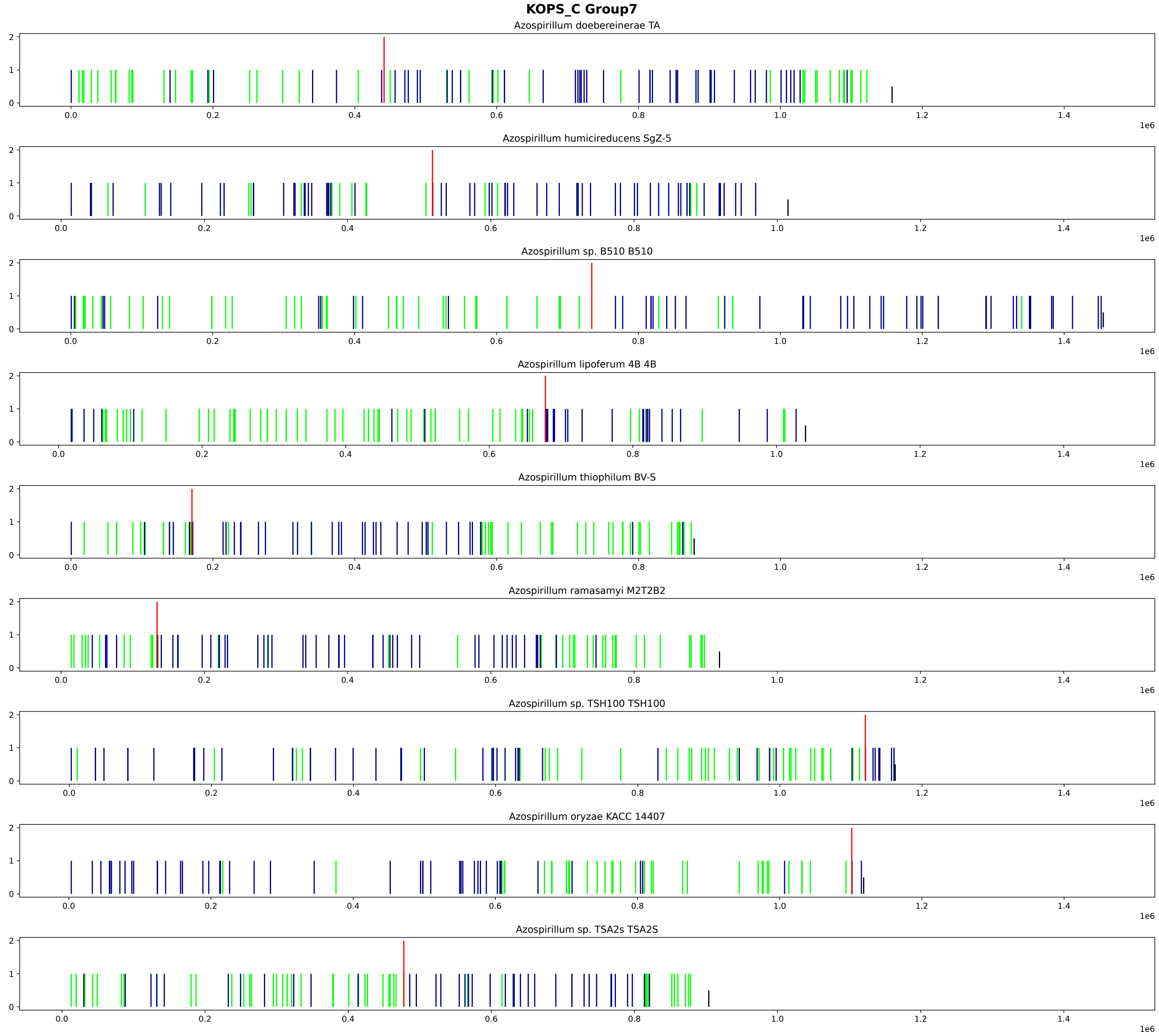
KOPS_C Group7
Azospirillum doebereinerae TA
Azospirillum humicireducens SgZ-5
Azospirillum sp. B510 B510
Azospirillum lipoferum 4B 4B
Azospirillum thiophilum BV-S
Azospirillum ramasamyi M2T2B2
Azospirillum sp. TSH100 TSH100
Azospirillum oryzae KACC 14407
Azospirillum sp. TSA2s TSA2S
1e6

### KOPS_C Group8

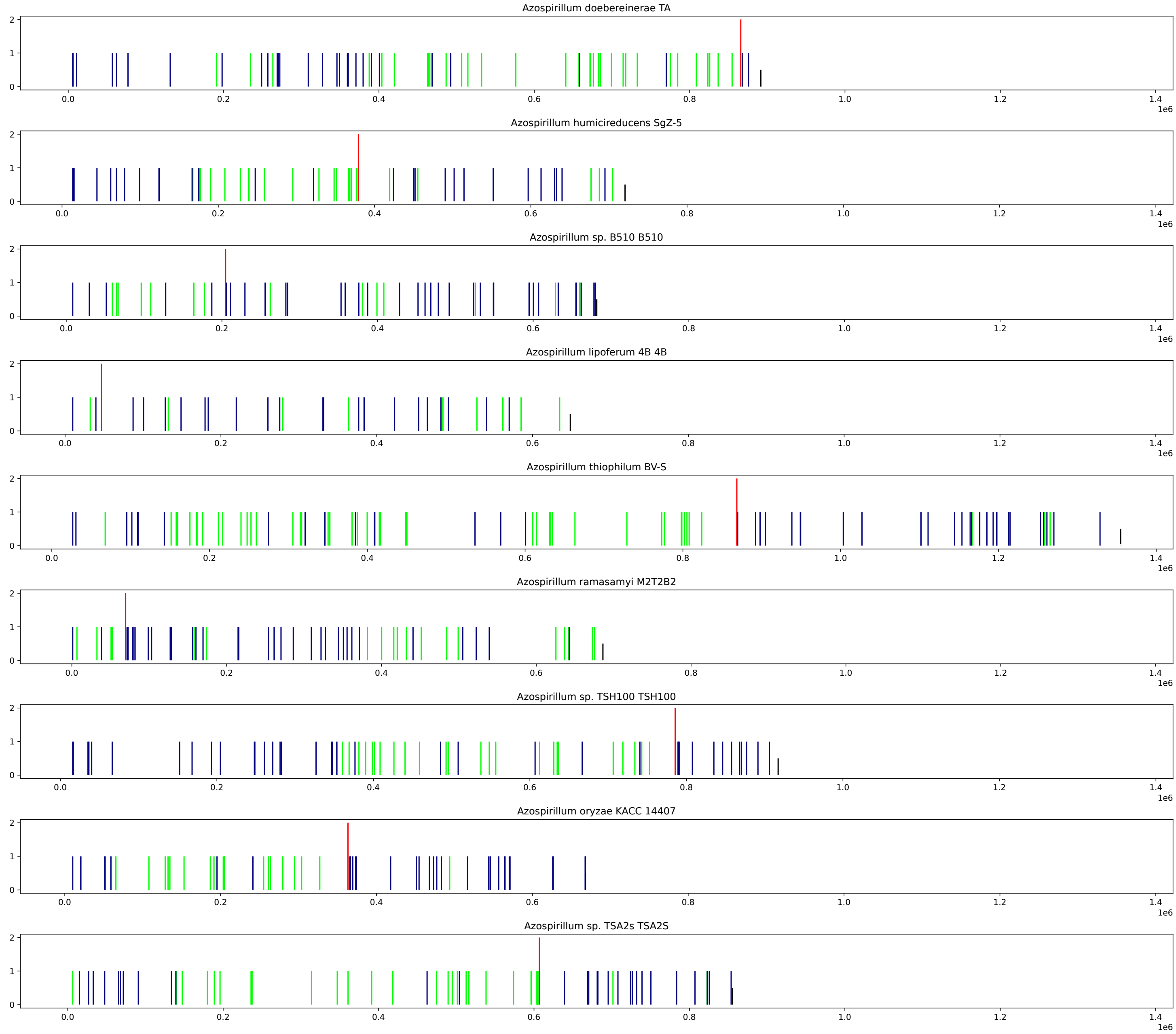

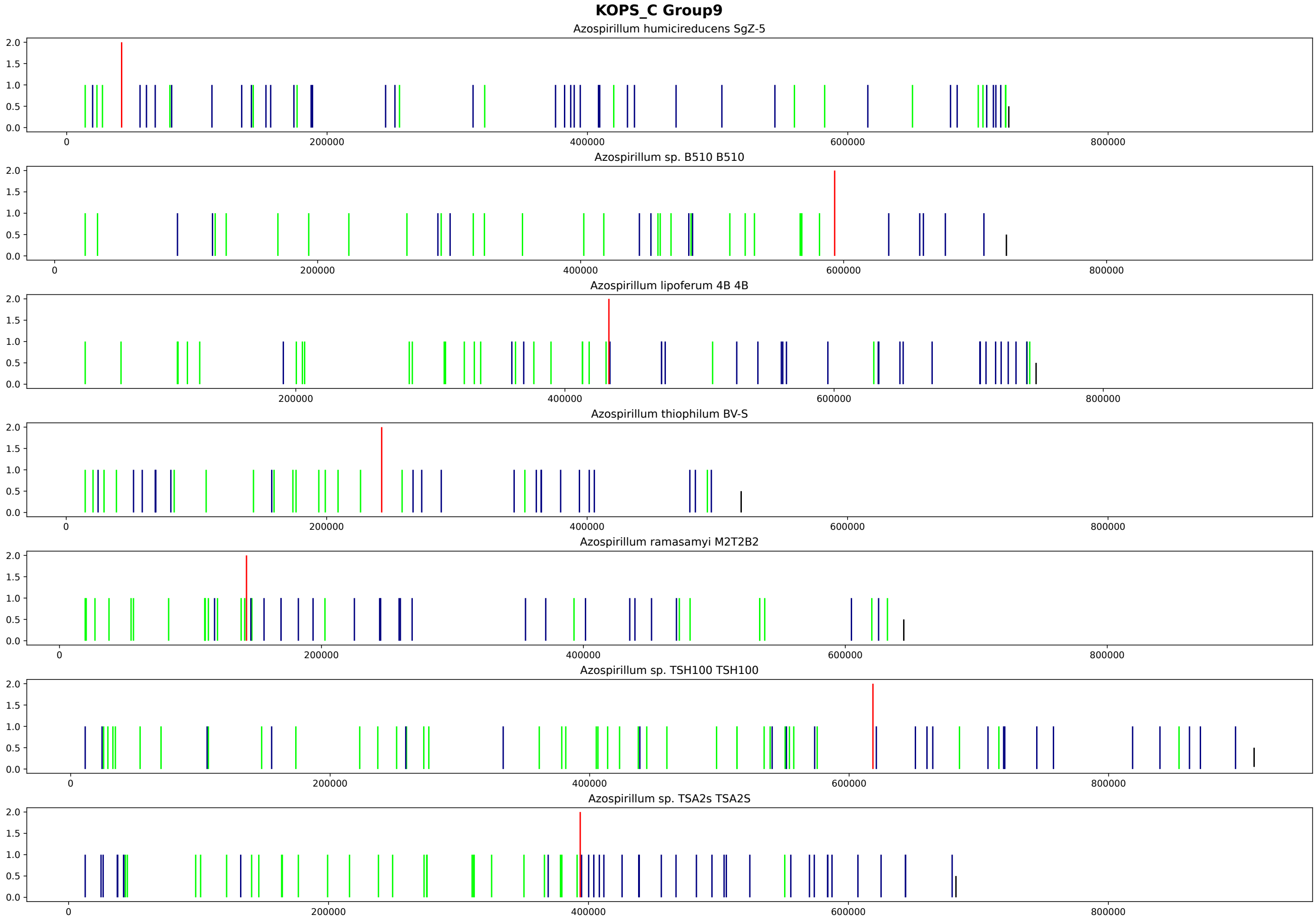
KOPS_C Group9
Azospirillum humicireducens SgZ-5
Azospirillum sp. B510 B510
Azospirillum lipoferum 4B 4B
Azospirillum thiophilum BV-S
Azospirillum ramasamyi M2T2B2
Azospirillum sp. TSH100 TSH100
Azospirillum sp. TSA2s TSA2S
0.0
0.5
1.0
1.5
2.0
0
200000
400000
600000
800000

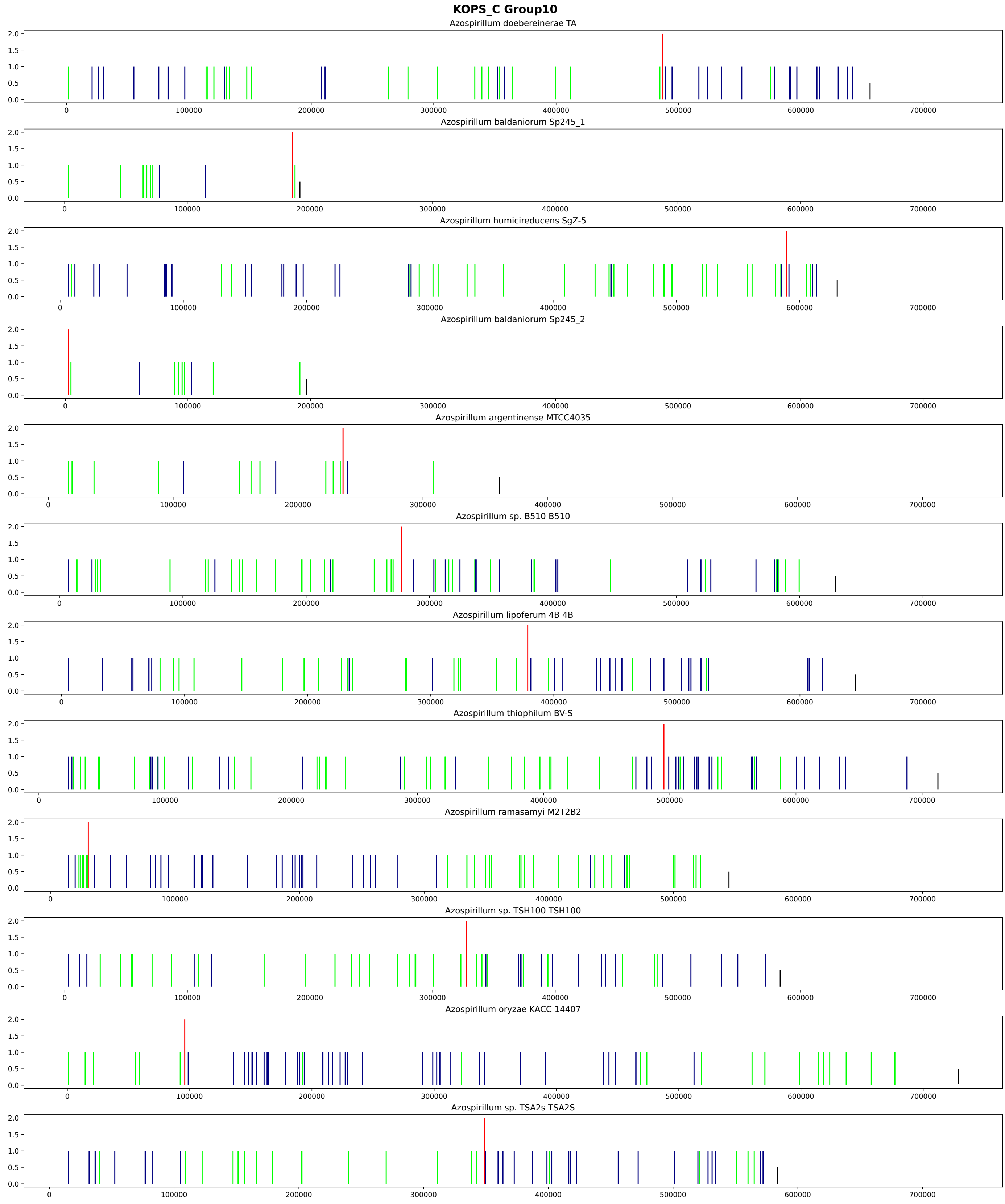
KOPS_C Group10
Azospirillum doebereinerae TA
Azospirillum baldaniorum Sp245_1
Azospirillum humicireducens SgZ-5
Azospirillum baldaniorum Sp245_2
Azospirillum argentinense MTCC4035
Azospirillum sp. B510 B510
Azospirillum lipoferum 4B 4B
Azospirillum thiophilum BV-S
Azospirillum ramasamyi M2T2B2
Azospirillum sp. TSH100 TSH100
Azospirillum oryzae KACC 14407
Azospirillum sp. TSA2s TSA2S
0.0
0.5
1.0
1.5
2.0
0
100000
200000
300000
400000
500000
600000
700000

**Supplementary Figure 10.** Cumulative GC skew, chi-square statistics for a sliding window with a step of 100, and the position of KOPS in both directions for all replicons of the strain, grouped by strain.

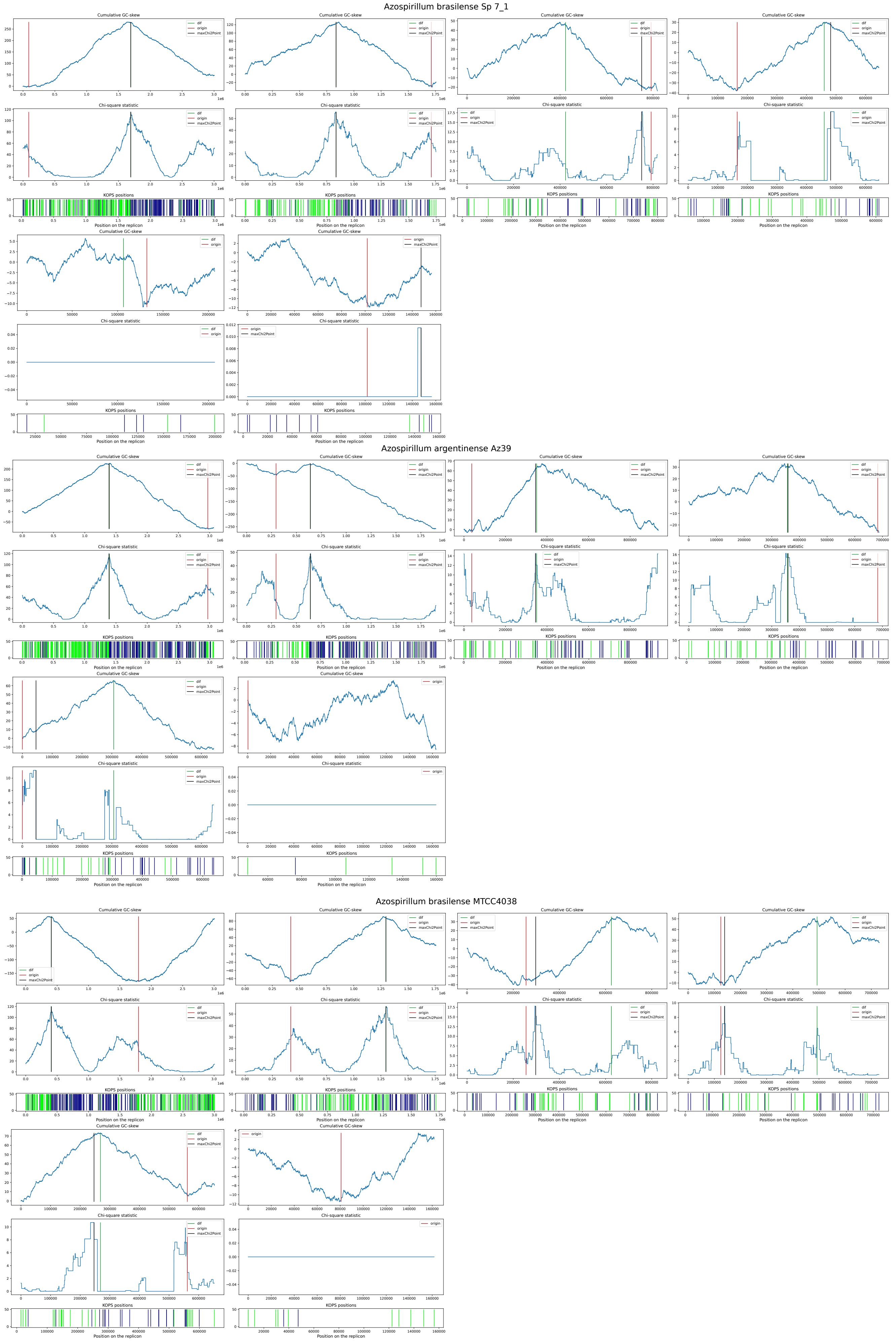

#### Azospirillum brasilense MTCC4039

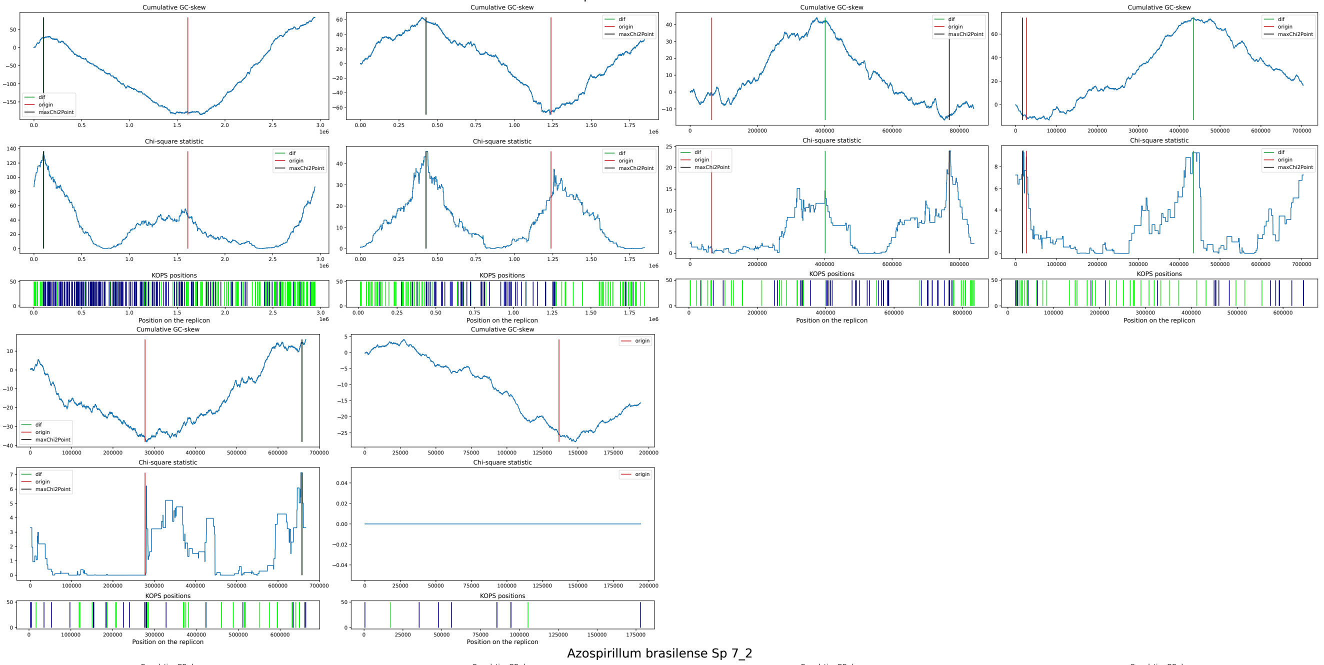


#### Azospirillum brasilense Sp 7_2

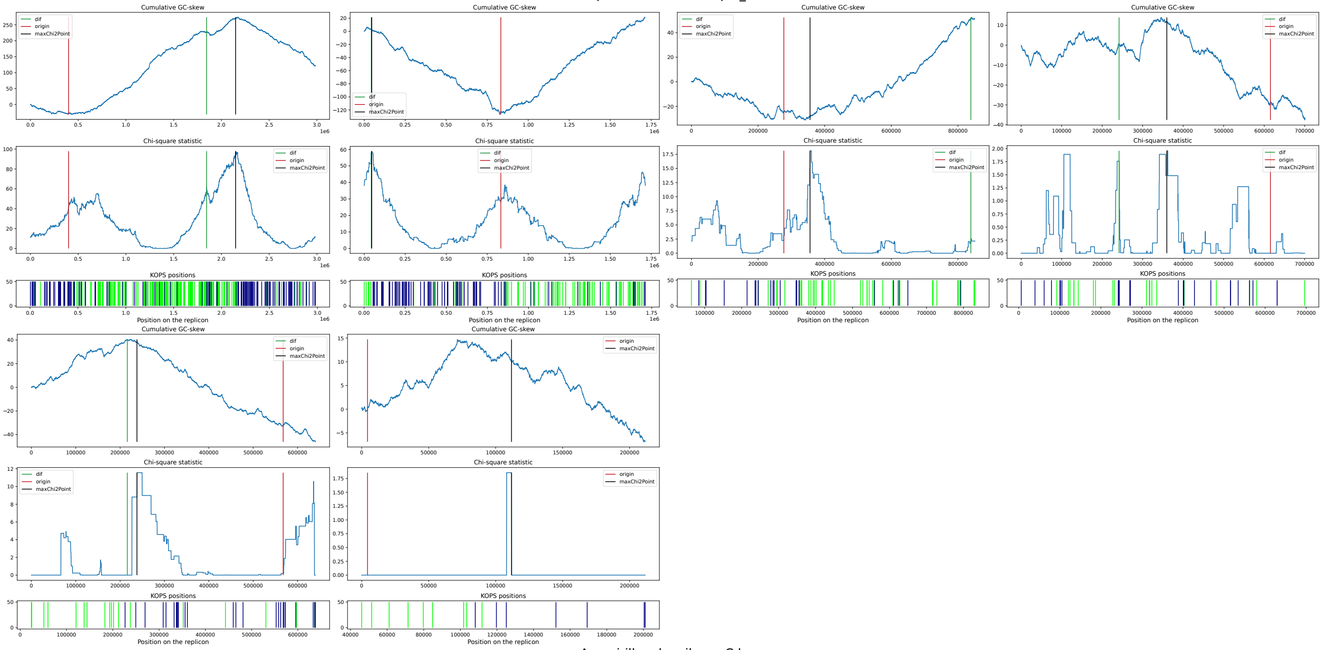


#### Azospirillum brasilense Cd

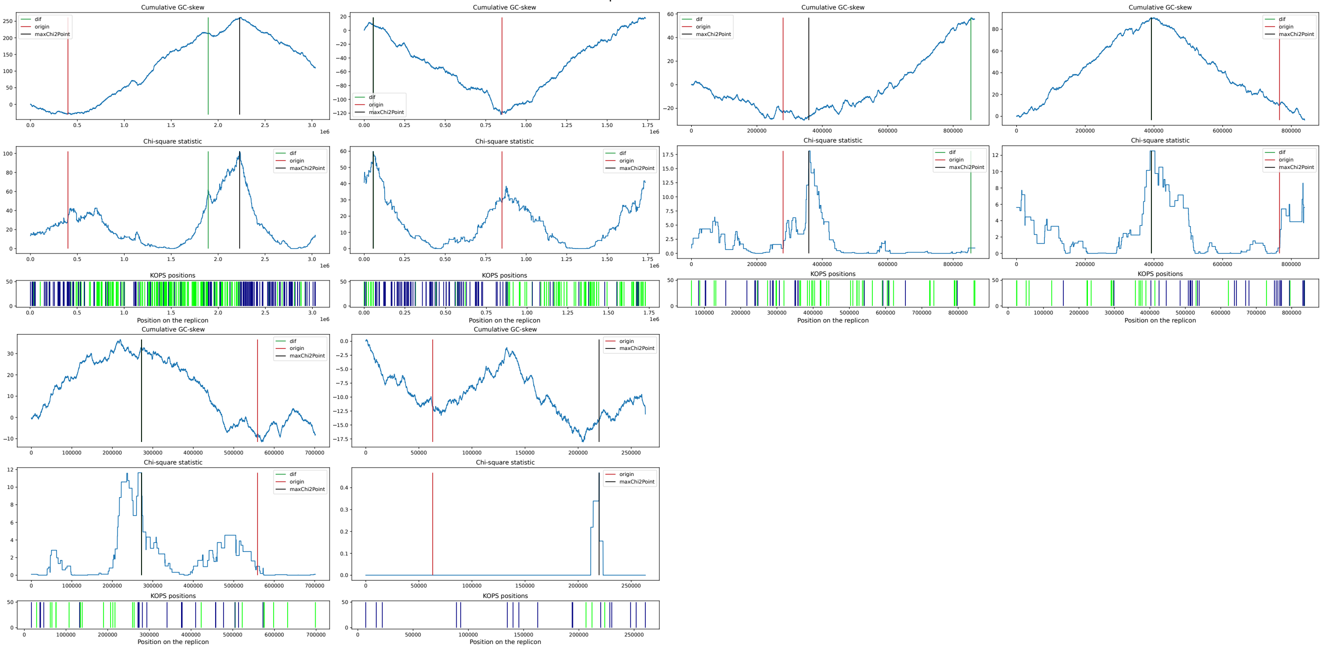

#### Azospirillum doebereinerae TA

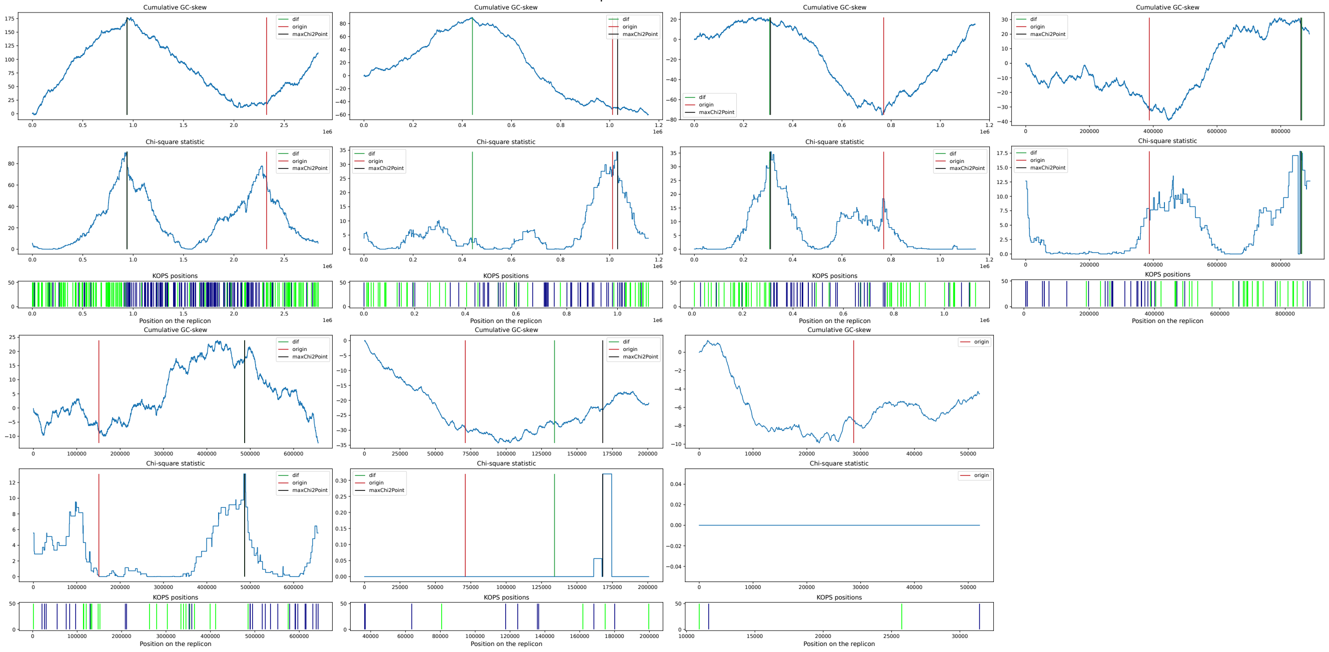


#### Azospirillum baldaniorum Sp245_1

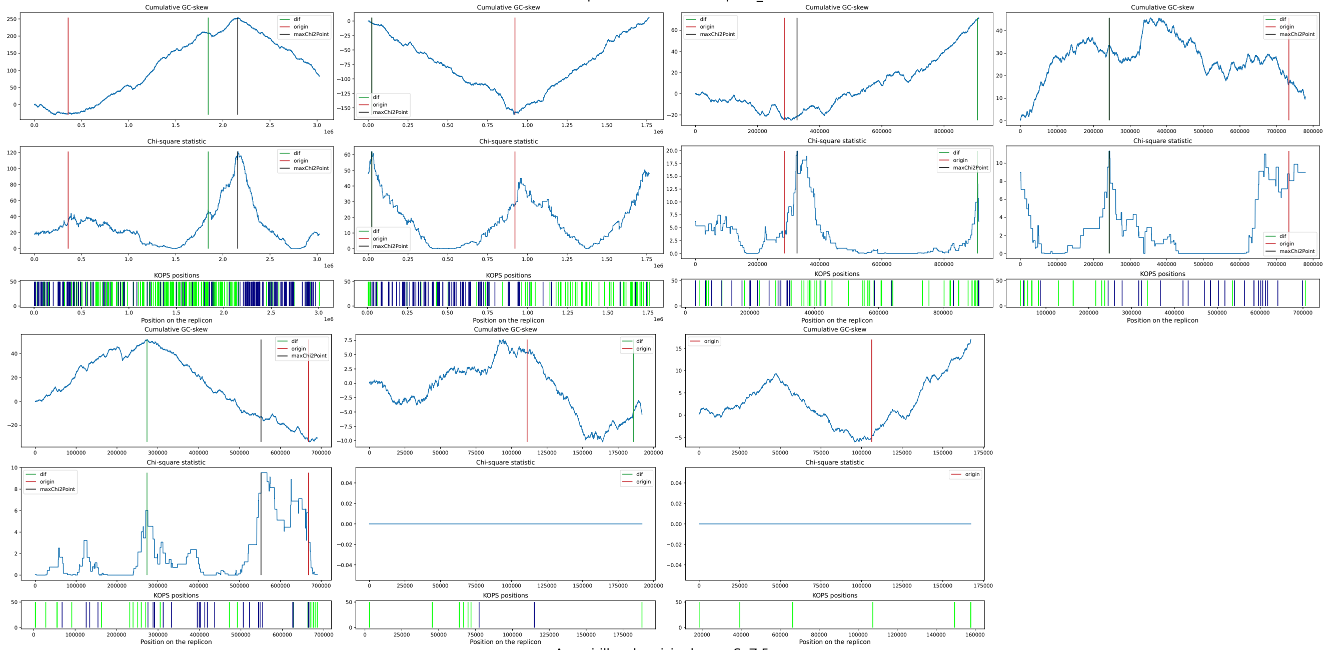


#### Azospirillum humicireducens SgZ-5

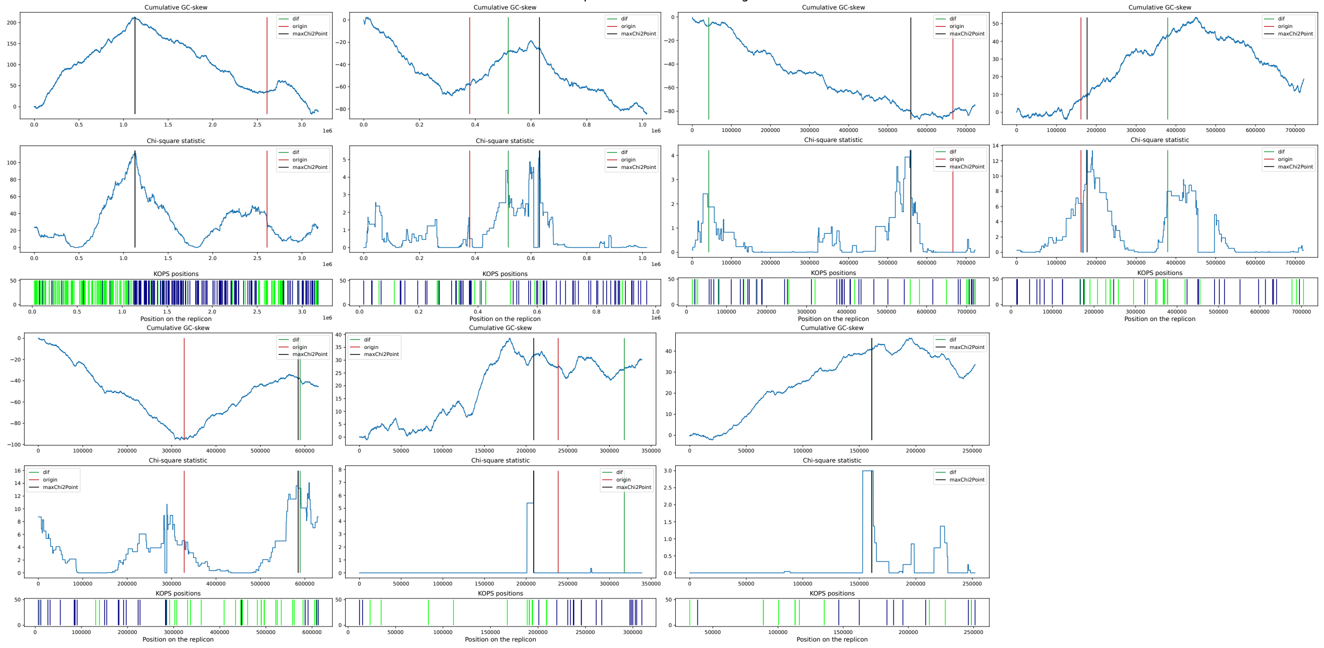

#### Azospirillum sp. TSH58 TSH58

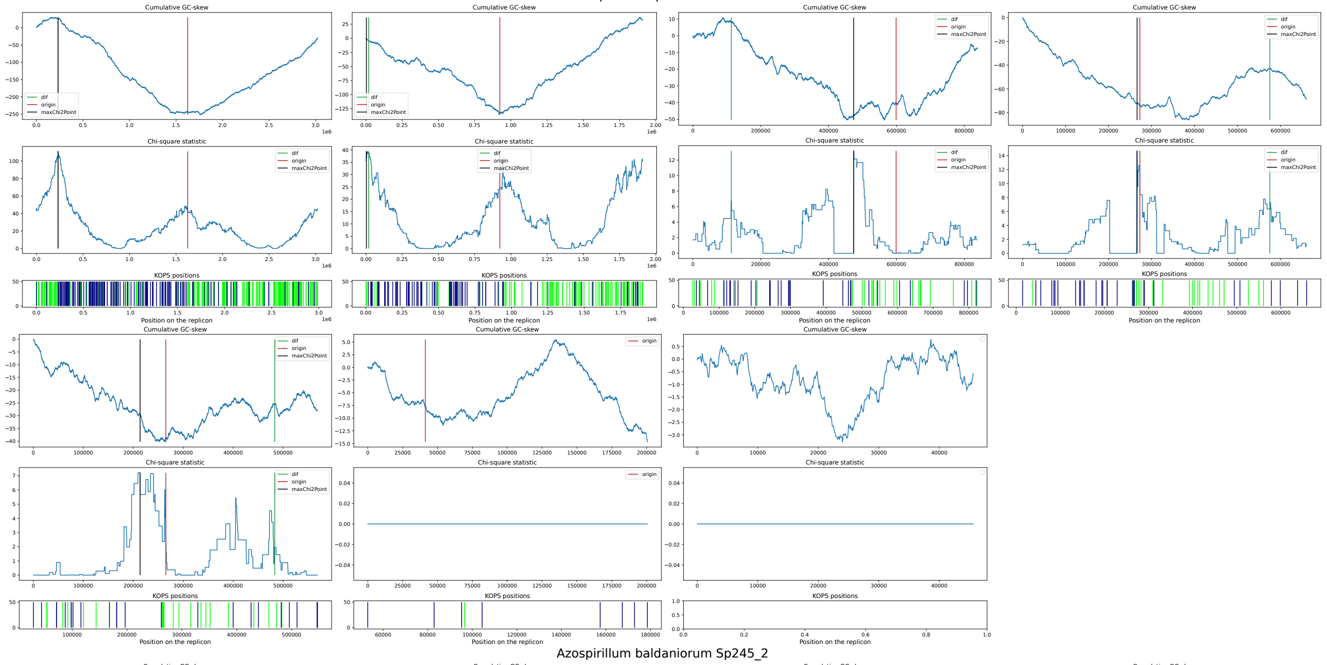


#### Azospirillum baldaniorum Sp245_2

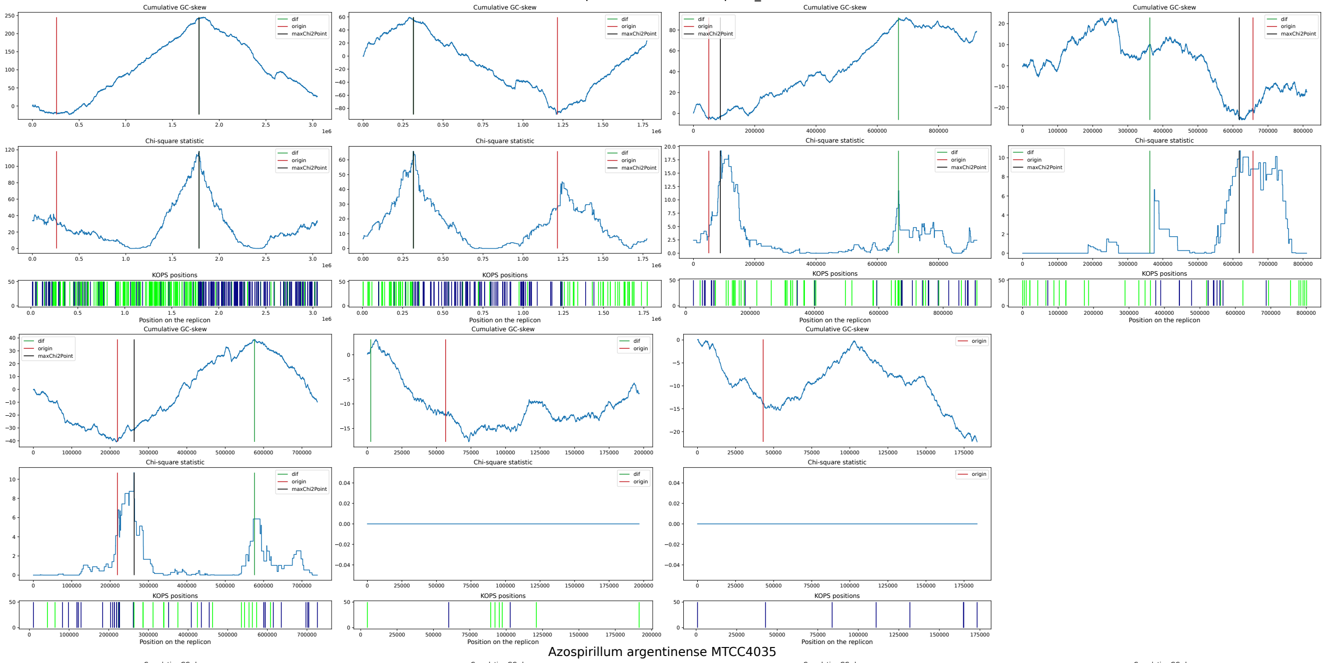


#### Azospirillum argentinense MTCC4035

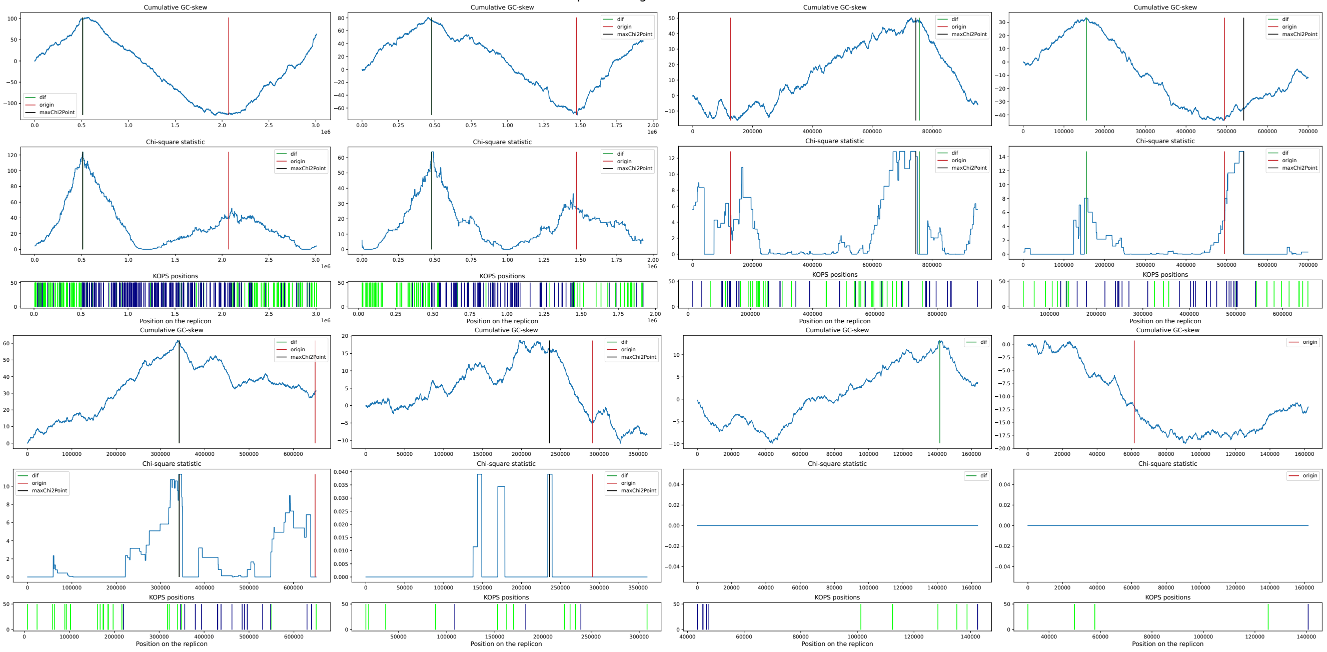

#### Azospirillum brasilense 2020WEIHUA_K

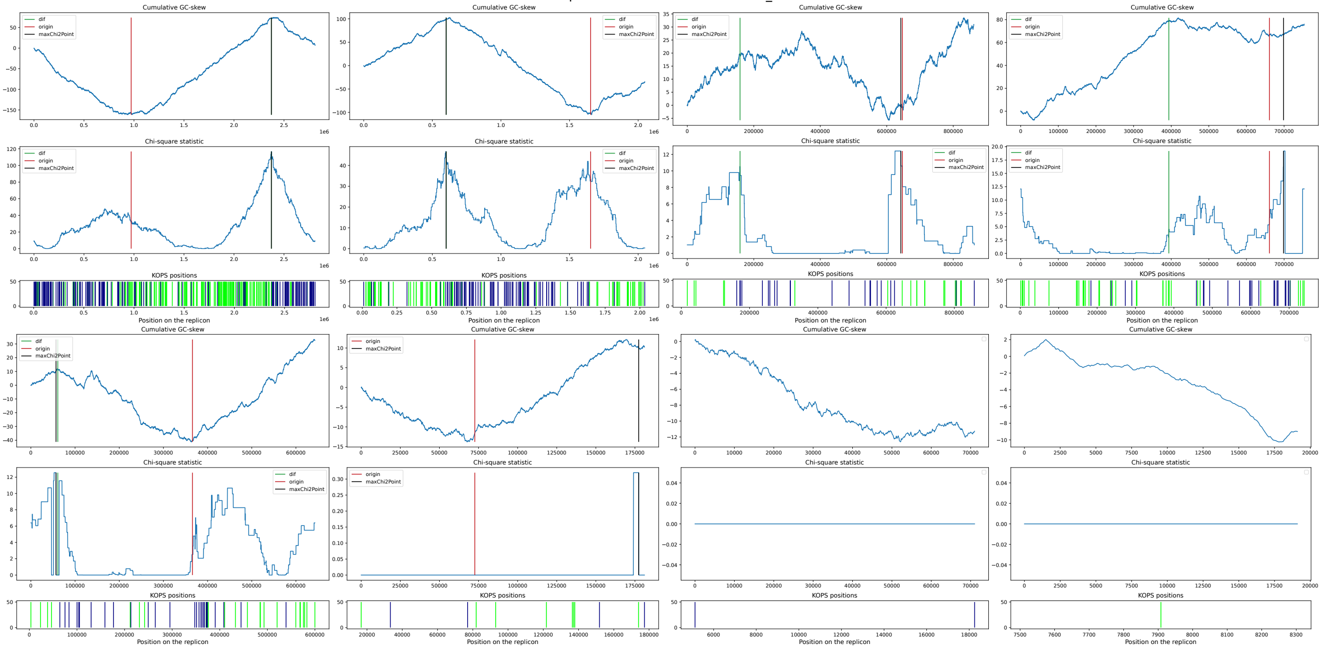


#### Azospirillum argentinense MTCC4036

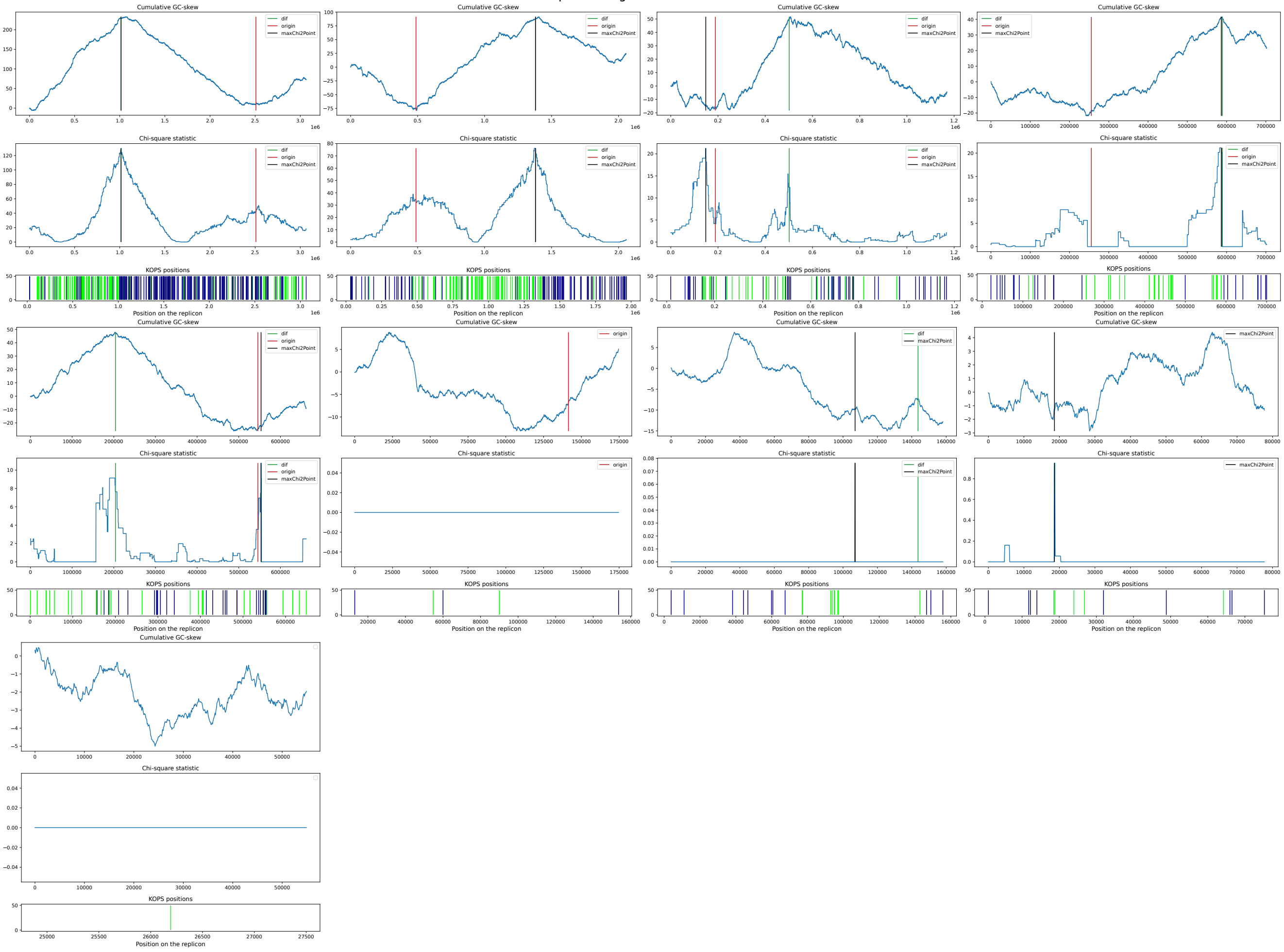

#### Azospirillum sp. B510 B510

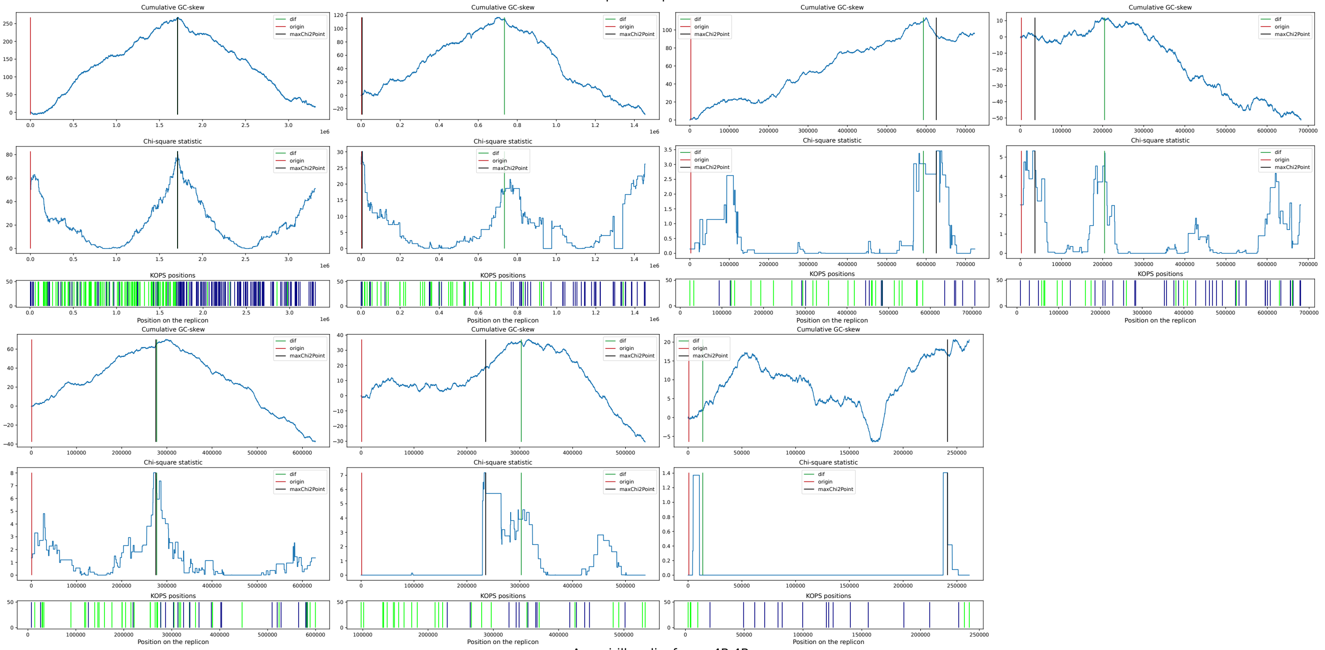


#### Azospirillum lipoferum 4B 4B

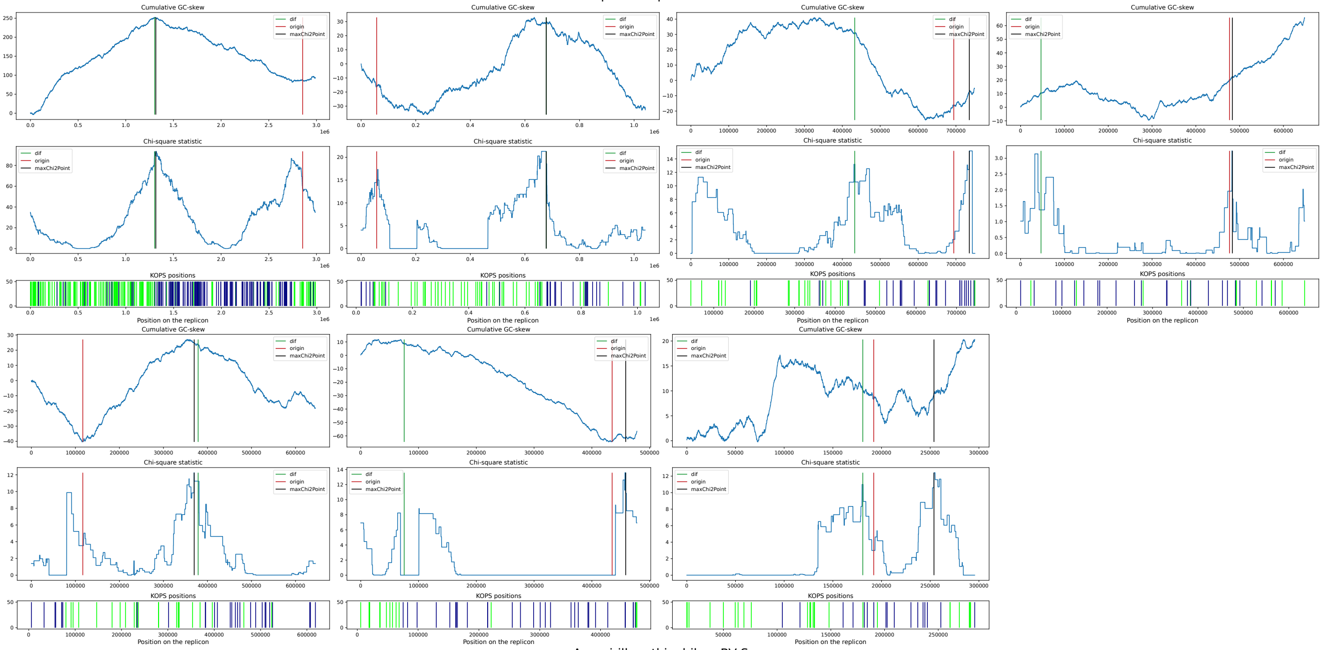


#### Azospirillum thiophilum BV-S

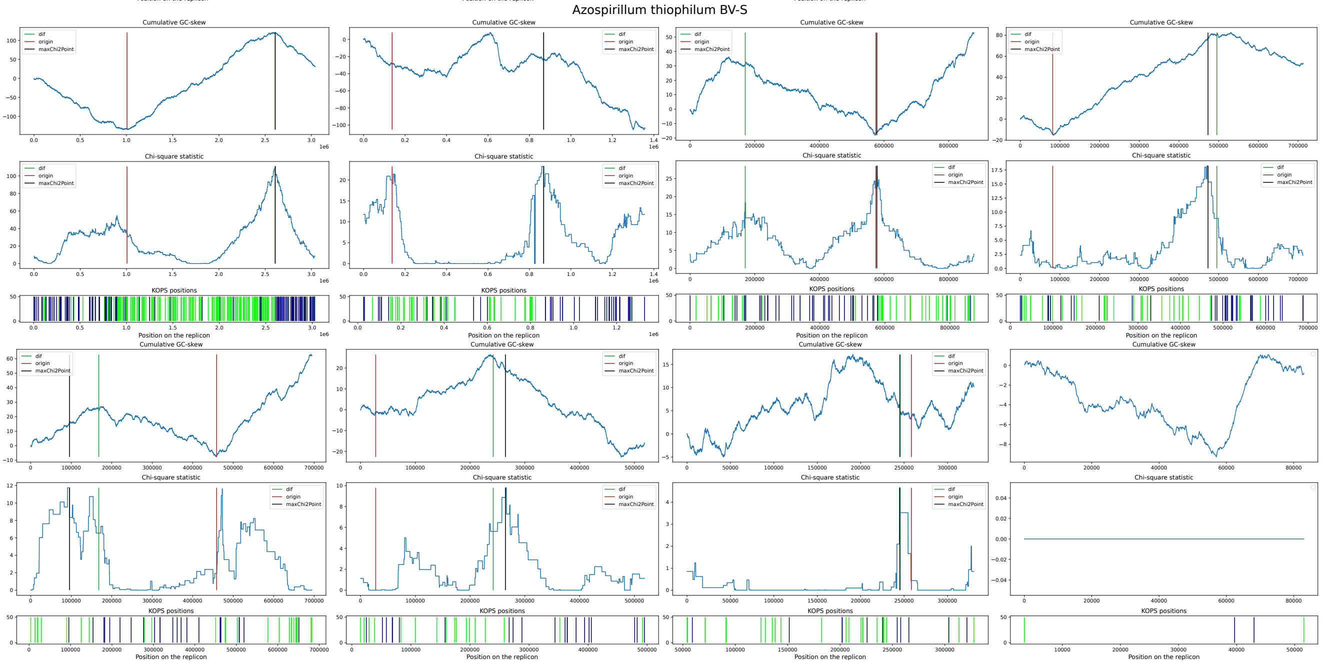

#### Azospirillum ramasamyi M2T2B2

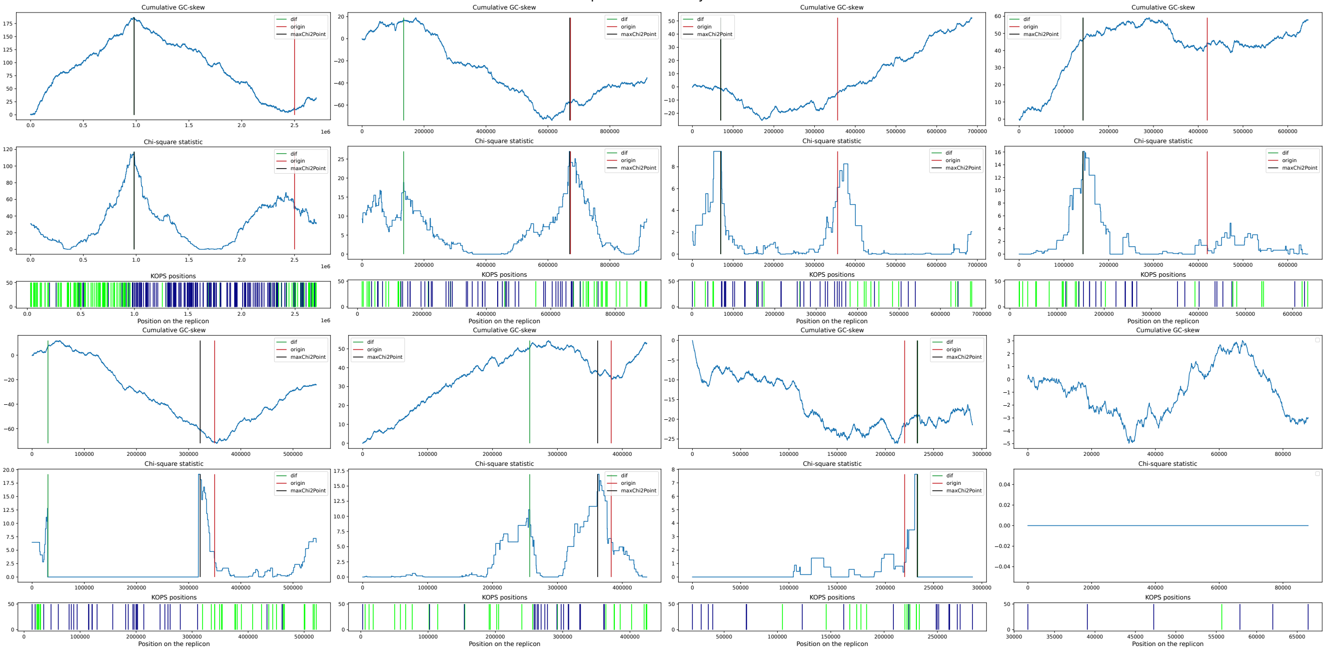


#### Azospirillum sp. TSH100 TSH100

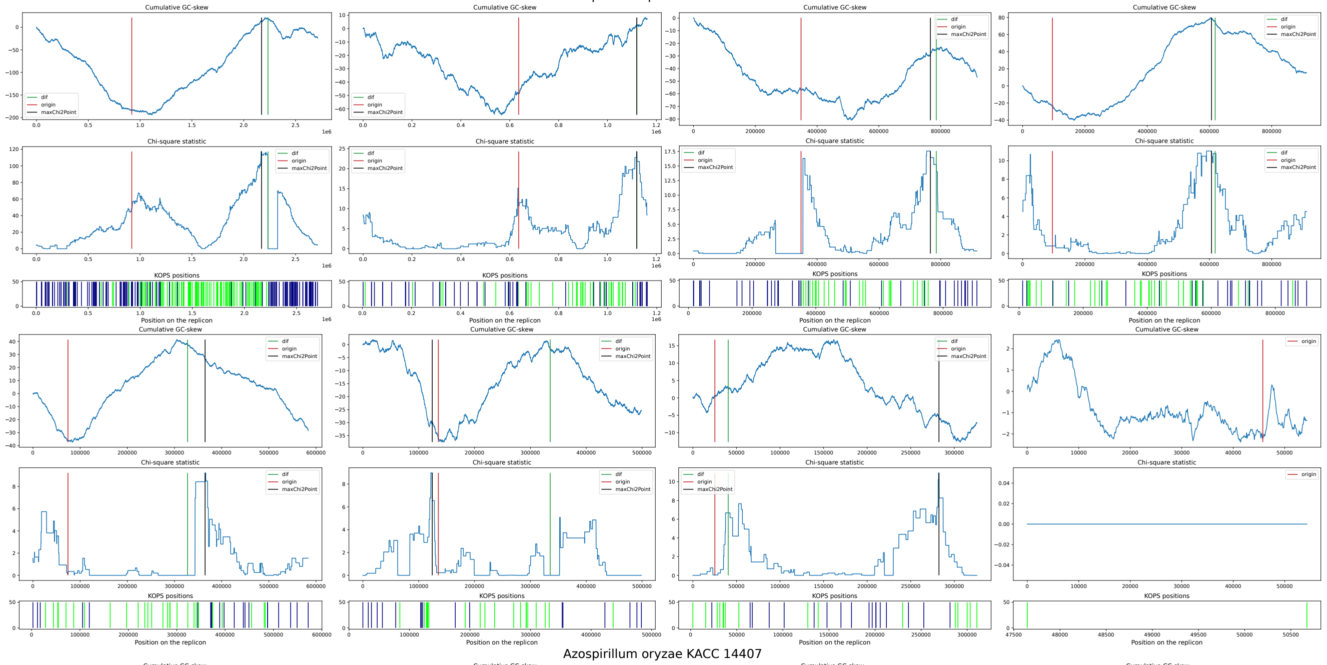


#### Azospirillum oryzae KACC 14407

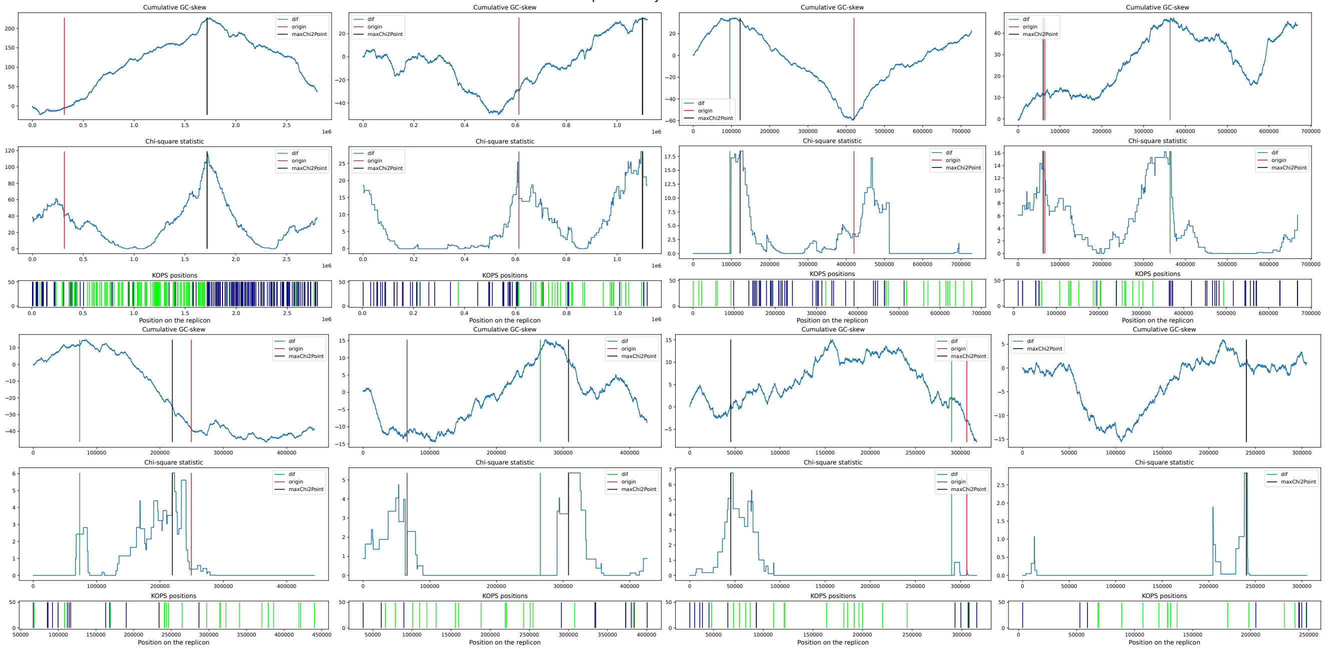

#### Azospirillum thermophilum CFH 70021

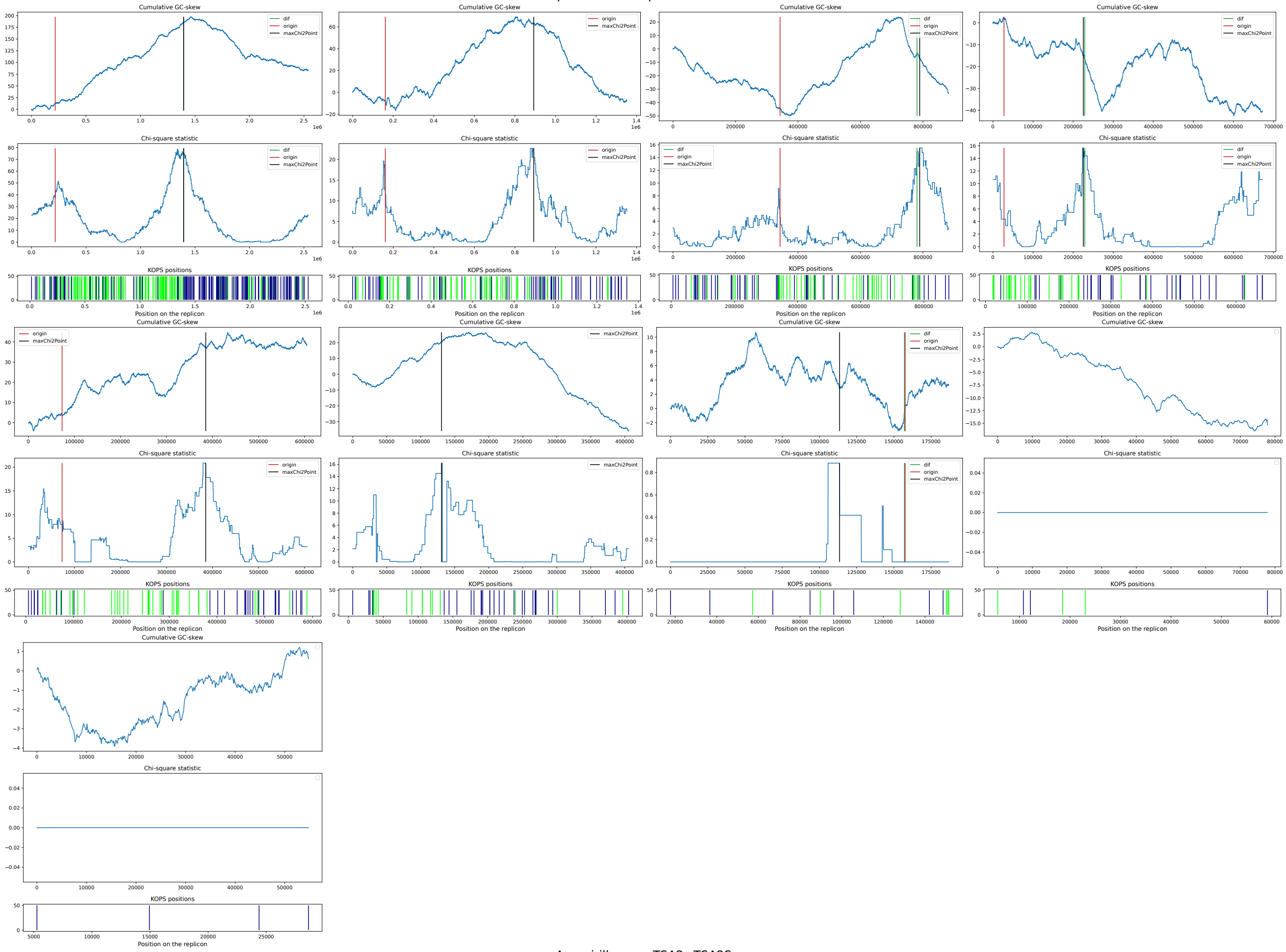


#### Azospirillum sp. TSA2s TSA2S

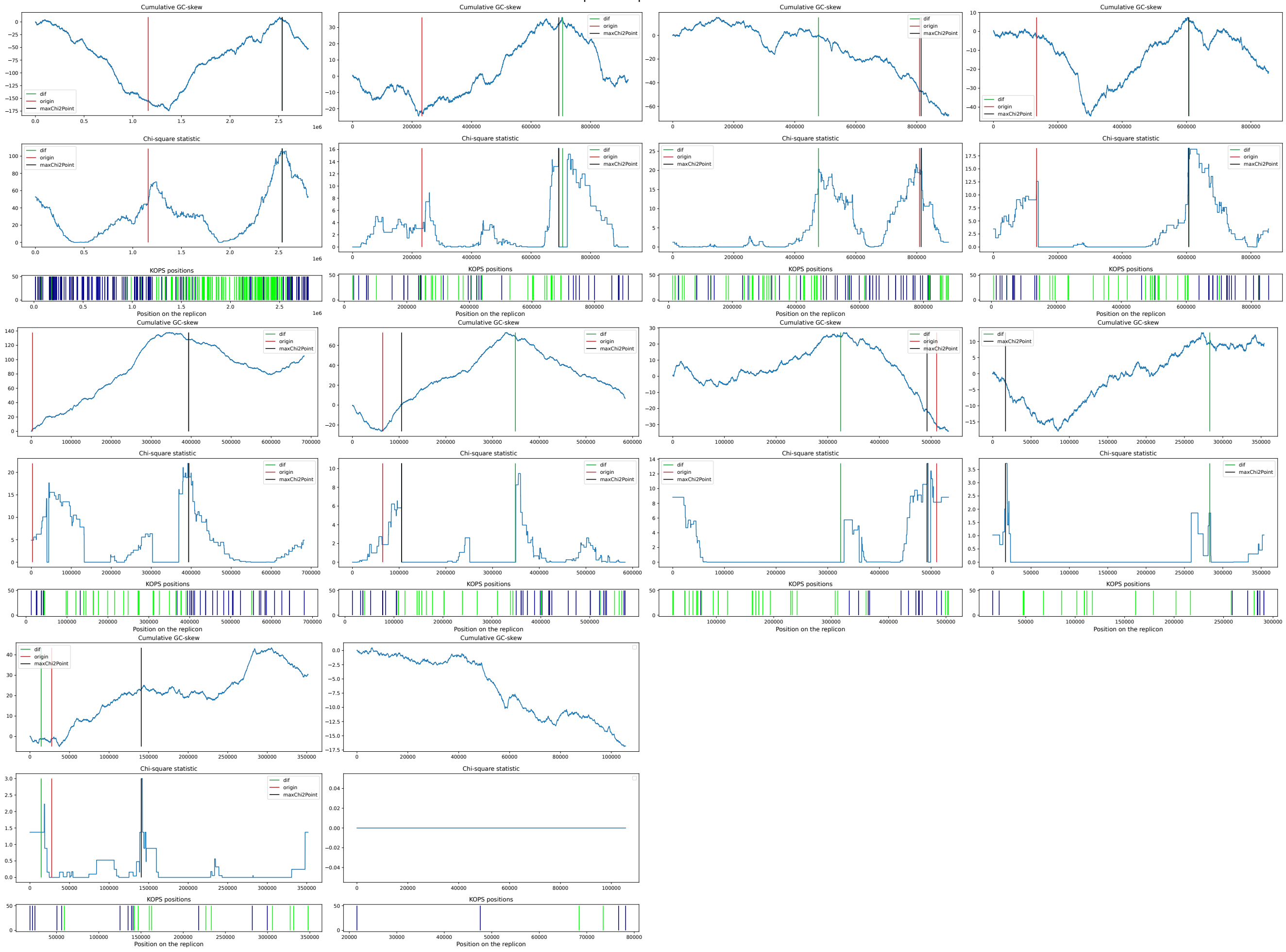

**Supplementary Figure 11.** KOPS positions in all chromosomes. The green and blue ticks of height 1 indicate KOPS in different directions. The pink and blue ticks of height 1.5 indicate the origin and terminus of replication. The red tick of height 2 indicates the position of the predicted dif site.

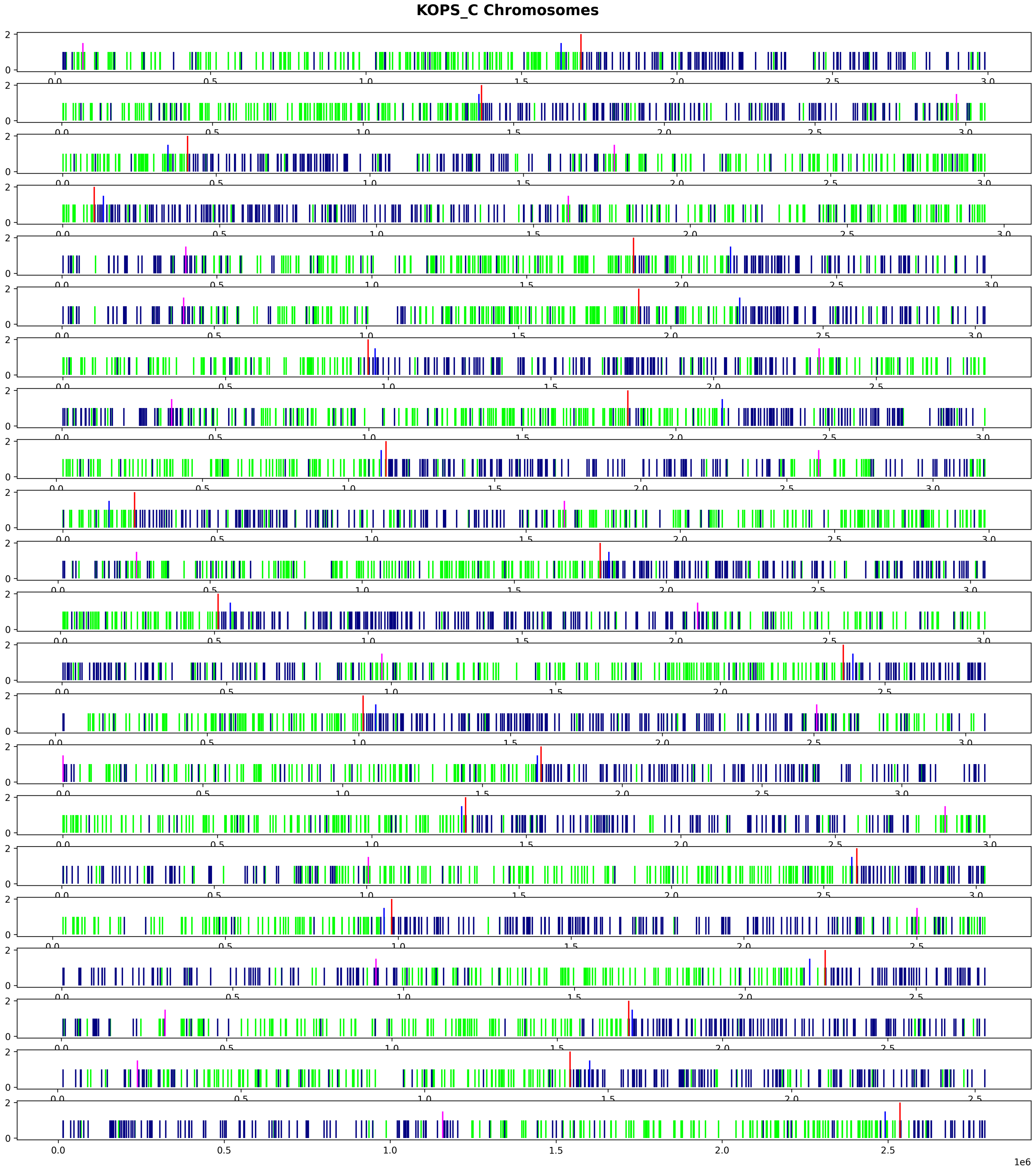
